# Inferring transcriptional bursting dynamics from scSLAM-seq data

**DOI:** 10.64898/2026.11.26.769093

**Authors:** Teresa Rummel, Yiliam Cruz Garcia, Bhupesh K. Prusty, Christophe Toussaint, Julius Deynet, Markus Vogt, Juliane Müller, Dirk Schadendorf, Bastian Schilling, Thomas Hennig, Lars Dölken, Antoine-Emmanuel Saliba, Elmar Wolf, Florian Erhard

**Author notes:** Correspondence: Elmar Wolf and Florian Erhard.

## Abstract

Transcription is a bursty, stochastic process. How transcriptional regulators reshape bursting dynamics during cellular responses remains poorly understood, largely because most single-cell RNA sequencing experiments measure only steady-state RNA abundance.

Here, we present buRst, a probabilistic framework that infers burst frequency, burst size, and RNA degra-dation rates from metabolically labeled single-cell RNA sequencing (scSLAM-seq) during acute perturbations, without assuming steady state. It jointly models labeling measurements and RNA dynamics, propagating uncertainty from sparse labeling data into kinetic estimates.

We applied buRst to acute MYC depletion in two melanoma cell lines. MYC amplified burst size in a manner scaling with its promoter occupancy, whereas it reduced burst frequency genome-wide, independently of occupancy. As a result, only strongly bound genes showed net activation, reconciling models of MYC as a selective activator and as a global amplifier of transcription.

## 1 Introduction

Even genetically identical cells in the same environment can differ substantially in RNA abundance because transcription and RNA degradation are probabilistic processes [1–3]. An important source of this variation is transcriptional bursting, in which RNA synthesis occurs intermittently rather than at a constant rate [3–6]. This behavior can be described by two effective parameters: burst frequency, describing how often bursts occur, and burst size, describing the average number of transcripts produced per burst. Together with the RNA degradation rate, these parameters determine RNA dynamics and describe the distribution of RNA counts across cells over time.

Single-cell RNA sequencing has enabled transcriptome-wide studies of transcriptional heterogeneity, revealing systematic variation in bursting properties across genes [7]. However, conventional scRNA-seq provides only a snapshot of total RNA abundance, reflecting both RNA production and degradation. Distinct combinations of burst size, burst frequency, and RNA turnover can therefore generate similar steady-state distributions. Inference requires strict steady-state assumptions, and even then individual kinetic parameters are only weakly identifiable from total RNA alone [7, 8].

Metabolic RNA labeling provides temporal information about RNA dynamics. In SLAM-seq, 4-thiouridine is incorporated into newly synthesized RNA during a defined labeling pulse and chemically modified to produce a characteristic T-to-C signature, allowing labeled transcripts to be identified by sequencing [9]. Because of limited 4sU incorporation, typically labeling less than 5% of uridines in the total RNA pool, and sequencing errors, newly synthesized molecule counts must be inferred from conversion patterns [10, 11]. Combining metabolic labeling with single-cell sequencing, including scSLAM-seq and sci-fate, enabled measurements of recent RNA synthesis and turnover at single-cell resolution [12–15]. However, single-cell sparsity exacerbates the challenge of estimating newly synthesized RNA fractions, resulting in substantial uncertainty.

Crucially, distinguishing strong from frequent bursts relies on the true variability of RNA abundance across cells. Measurement uncertainty obscures this variability, challenging unbiased estimation of bursting parameters. Edwards *et al.* [8] demonstrated that combining total and newly synthesized RNA can resolve kinetic parameters that are difficult to infer from total RNA alone. However, their Bayesian framework models newly synthesized and pre-existing RNA relative to an underlying steady-state distribution. This leaves a need for computationally tractable frameworks that propagate labeling uncertainty across many genes and conditions, particularly when steady-state assumptions cannot be made.

MYC provides a relevant system for distinguishing alternative modes of transcriptional regulation. As a central regulator of cellular growth, metabolism, and biosynthetic activity, MYC controls broad transcriptional programs, yet its mode of action remains debated [16–19]. MYC has been proposed to amplify the output of active genes, partly by promoting RNA polymerase II pause release [20–23]. Other studies support selective activation and repression of defined gene sets and suggest that MYC also influences transcription initiation [24, 25]. Acute MYC degradation experiments linked immediate transcriptional dependence to quantitative MYC occupancy and promoter context [26], while live-cell measurements found that MYC overexpression increased burst duration rather than frequency [27]. However, how these effects translate into transcriptome-wide changes in effective burst frequency, burst size, and RNA degradation following acute MYC depletion remains unclear.

Here, we introduce buRst, a likelihood-based framework for inferring transcriptional bursting kinetics from scSLAM-seq data. The framework combines total and newly synthesized RNA to estimate burst frequency, burst size, and RNA degradation, including when the perturbed condition is not at steady state. Through latent-state marginalization, buRst propagates uncertainty in newly synthesized RNA assignments rather than treating estimated counts as exact observations. We validate the framework using simulated data and compare its accuracy, uncertainty estimates, and computational performance with burstMCMC [8]. Applying buRst to acute MYC depletion, we identify coordinated changes in burst size and frequency, with the strongest transcriptional responses and promoter-associated MYC occupancy more clearly linked to burst size changes. Together, these results provide a transcriptome-wide view of how MYC influences transcriptional bursting.

## 2 Results

### buRst: A probabilistic framework for inferring transcriptional bursting kinetics from scSLAM-seq data

We developed buRst, a probabilistic framework for inferring transcriptional bursting kinetics directly from scSLAM-seq measurements (Figure 1). Gene expression is modeled by transcriptional bursts occurring at rate *k_m_*, each producing a geometrically distributed number of RNA molecules with mean burst size *B*, and independent RNA degradation at rate *d* [3, 4, 6, 28]. At steady state, RNA counts follow a negative binomial distribution. For this and other previously used models [7, 29, 30], the degradation rate is structurally unidentifiable from total RNA alone. Even when burst frequency is expressed in units of the RNA half-life, burst size and frequency are practically unidentifiable for relevant parameter ranges with noisy data (Supplementary Figure S1a).

**Figure 1:**
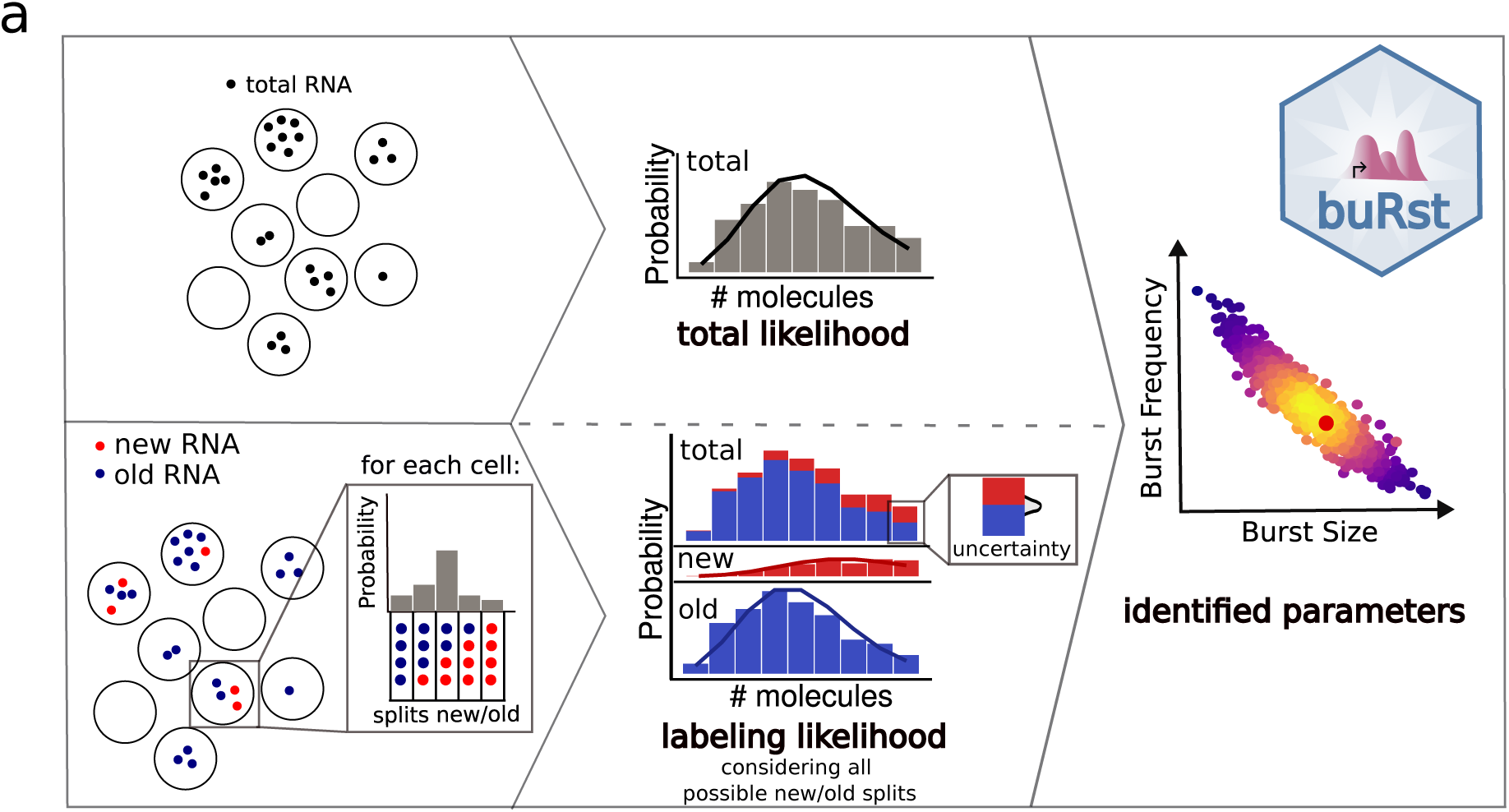
Schematic overview of the probabilistic framework for inference of transcriptional bursting kinetics from scSLAM-seq data.

To address these identifiability challenges and infer parameters under non-steady-state conditions, we incorporated metabolic RNA labeling information. In scSLAM-seq, newly synthesized RNA is labeled with 4-thiouridine, typically for 2 hours, and detected through T-to-C mismatches [9, 12]. The distribution of RNA transcribed during labeling helps distinguish kinetic parameter combinations unresolved by total RNA alone (Supplementary Figure S1a). We derived analytical probability-generating functions for the RNA-count distributions and developed an efficient numerical evaluation based on the fast Fourier transform (Methods), enabling parameter inference.

Because sequencing coverage is sparse and conversion events are rare, newly synthesized RNA counts are not observed directly. Rather than assigning transcripts to discrete “new” and “old” states, buRst considers all possible partitions into newly synthesized and pre-existing populations. For each partition, we combine the likelihood of the observed mismatch patterns with the corresponding bursting-model likelihood. Marginalizing over these partitions propagates labeling uncertainty directly into kinetic parameter inference.

At steady state, mean RNA abundance constrains the product of burst frequency and size (*λ*), whereas variability across cells helps distinguish large, infrequent bursts from small, frequent bursts. Propagating labeling uncertainty is therefore essential to tell apart biological variability from uncertainty in newly synthesized RNA counts and avoid biased estimates of burst size and frequency (Supplementary Figure S1b–c). By combining total and newly synthesized RNA information while accounting for labeling uncertainty, buRst infers bursting kinetics with parameter-specific uncertainty estimates (Figure 1).

### buRst accurately recovers transcriptional bursting parameters with calibrated uncertainty

To evaluate the statistical performance of buRst against known ground truth, we performed a simulation experiment. We first inferred realistic gene-specific burst sizes, burst frequencies, and degradation rates from the 0 h time point of the sci-fate dataset [13]. These parameters served as ground truth for simulating 2-hour RNA-labeling experiments for 8,254 genes and 1,151 cells. Simulations generated new and pre-existing RNA counts together with read-level labeling evidence, reproducing the inputs used by buRst. Parameters were then re-estimated and compared with their true values (Figure 2a).

**Figure 2:**
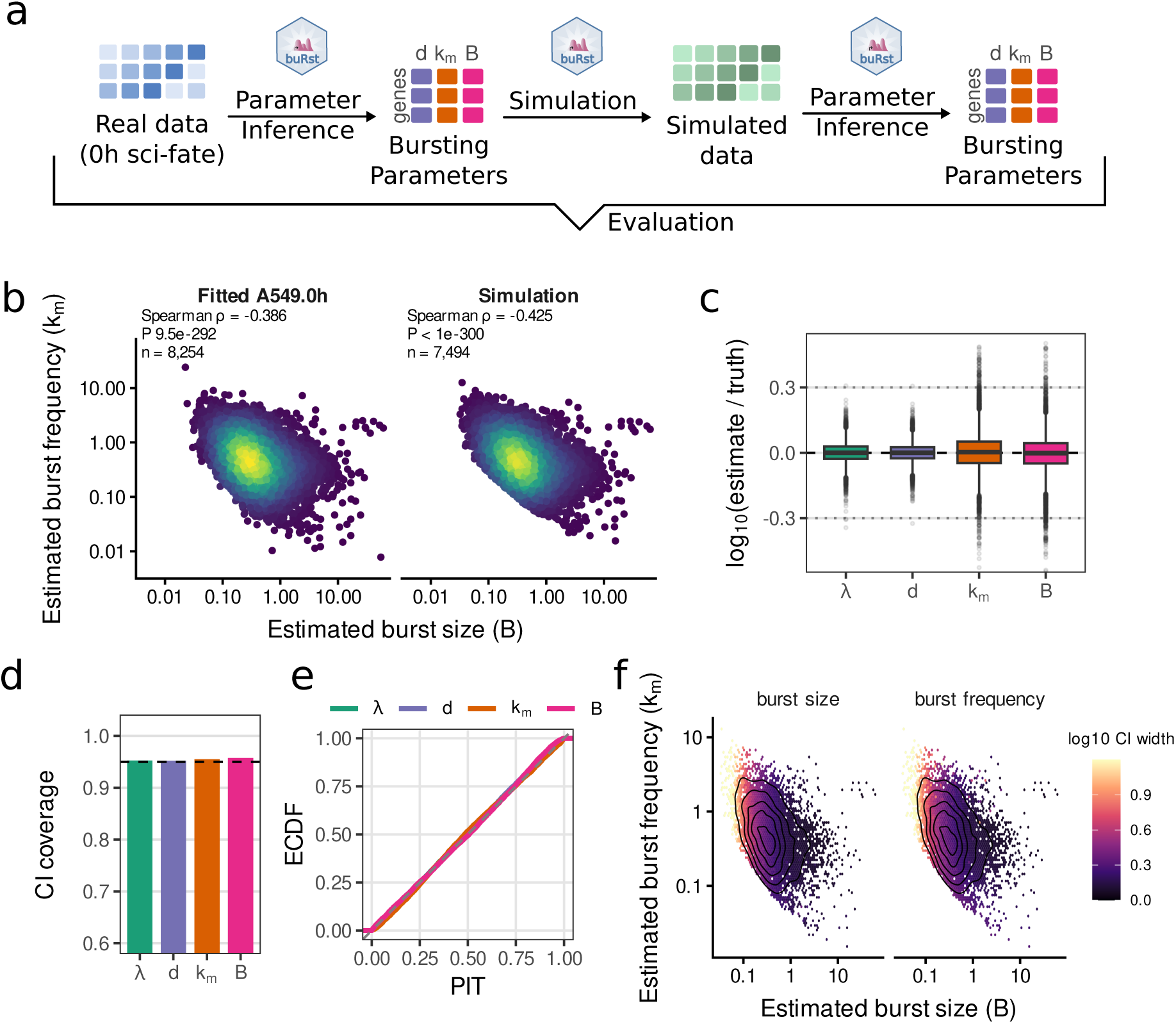
Validation of transcriptional bursting parameter inference using simulated scSLAM-seq data. (**a**) Schematic overview of the simulation and parameter re-estimation workflow. (**b**) Joint distribution of burst size (*B*) and burst frequency (*k_m_*) inferred from the sci-fate 0 h dataset (*n* = 8,254 genes; left) and from simulated data (*n* = 7,494 genes; right). Colors represent local point density. Spearman’s rank correlation coefficients were *ρ* = *−*0.386 (*P* = 9.5 *×* 10*^−^*^292^) for the sci-fate 0 h dataset and *ρ* = *−*0.425 (*P <* 10*^−^*^300^) for the simulated data. For the matched subset of 7,494 genes with successful inference, the ground-truth correlation was *ρ* = *−*0.404 (*P* = 5.48 *×* 10*^−^*^292^). (**c**) Parameter estimation errors for *n* = 7,494 genes, expressed as log_10_(estimate*/*truth). Here, *λ* = *k_m_B* denotes the mean RNA production rate, *d* the RNA degradation rate, *k_m_* the burst frequency, and *B* the burst size. The dashed line indicates exact recovery, and the dotted lines at *±*0.3 indicate approximately twofold over- or underestimation. (**d**) Empirical coverage of the 95% approximate posterior credible intervals (CI) for *n* = 7,494 genes. The dashed line indicates the nominal coverage level. (**e**) Empirical cumulative distribution functions of the probability integral transform values for *n* = 7,494 genes for each parameter. The dashed diagonal indicates the expected distribution under calibrated inference. (**f**) Hexbin plots showing estimated burst sizes and burst frequencies for *n* = 7,494 genes. Color denotes the mean log_10_ width of the 95% approximate posterior credible interval within each bin for burst size (left) or burst frequency (right). Black contour lines indicate the density of genes across the kinetic parameter landscape.

Re-estimated parameters closely matched the ground truth (Figure 2b). Re-estimated burst sizes ranged from 0.09 to 2.29 detected transcripts per burst and frequencies from 0.11 to 2.14 per hour (5th–95th percentiles), consistent with previously reported values [7, 15]. Among the 7,494 genes with successful inference, the negative correlation between burst size and frequency was preserved (Spearman’s *ρ* = *−*0.404 for ground-truth parameters restricted to these genes and *ρ* = *−*0.425 for re-estimated parameters; both *P <* 10*^−^*^290^). The full reference set of 8,254 genes, shown in the left panel, had a correlation of *ρ* = *−*0.386. Thus, buRst preserved key features of the simulated kinetic landscape.

The 90th percentiles of fold errors in expression strength *λ* (the product of burst frequency and size), decay rate *d*, burst frequency *k_m_*, and burst size *B* were 1.20, 1.18, 1.47, and 1.45, respectively. Log-error distributions were centered near zero, suggesting unbiased estimation (Figure 2c).

Approximate posterior credible intervals had nominal coverage across all four parameters (Figure 2d). Probability integral transform (PIT) values were uniformly distributed (Figure 2e). Together, these analyses indicate that the approximate posterior distributions are calibrated.

To investigate identifiability, we assessed uncertainty across the inferred kinetic landscape. Genes with small burst sizes and high burst frequencies generally exhibited greater uncertainty, whereas genes with greater RNA abundance showed better identifiability of both parameters (Figure 2f). This pattern is consistent with limited information for sparsely expressed genes and indicates that estimated uncertainty reflects read availability.

Distinguishing strong from frequent bursts requires separating true expression variance from measurement uncertainty in labeled RNA. We therefore compared our new uncertainty propagation approach (latent-state marginalization) with our previous beta approximation method [10, 31] and purely using point estimates of labeled RNA. Point estimates overestimated true expression variance, inflating burst sizes and underestimating burst frequencies (Supplementary Figure S1d). Conversely, the beta approximation apparently underestimated true expression variance, strongly biasing both parameters in the opposite direction. Both approaches also affected decay rate and expression strength estimates and substantially reduced coverage of the approximate posterior credible intervals (Supplementary Figure S1e). In contrast, the full labeling likelihood yielded estimates centered at the true values and nominal coverage, demonstrating the importance of accounting for combinatorial labeling complexity.

We further benchmarked buRst against burstMCMC, a Bayesian framework for inferring bursting kinetics from single-cell 4sU-labeling data [8]. Both methods were evaluated under identical simulated conditions across 8,254 genes. With a per-gene timeout of 10,000 seconds, 742 burstMCMC attempts (9.0%) timed out and were excluded from subsequent comparisons. Overall, 7,494 genes passed quality criteria for buRst and 5,461 for burstMCMC, with 5,218 passing for both methods and entering the direct comparison (Supplementary Figure S2a). buRst showed comparable or improved parameter recovery and interval coverage (Supplementary Figure S2b,c). Moreover, inference with buRst took less than 2 h, comparable to read mapping, whereas burstMCMC required approximately three orders of magnitude longer per-gene runtime (Supplementary Figure S2d), rendering it unsuitable for large-scale analyses. Thus, in addition to being the only tool applicable under non-steady-state conditions buRst outperformed burstMCMC in terms of accuracy, robustness and computational efficiency.

Together, these results demonstrate accurate inference of transcriptional bursting kinetics from single-cell SLAM-seq data with calibrated uncertainty estimates.

### buRst detects and classifies non-steady-state changes in transcriptional bursting

Having established accurate parameter recovery at steady state, we next assessed whether buRst detects and mechanistically classifies acute changes in transcriptional bursting.

For comparisons between conditions, buRst analyzes the joint posterior distribution of log_2_ fold changes in burst frequency and size. A change is considered supported when it is statistically well-separated from zero (origin outside the 95% joint credible ellipse) and at least 90% of the posterior mass lies outside a region of practical equivalence around zero. Changes are classified as frequency dominated, size dominated, or combined when at least 60% of the posterior mass can be attributed to one of these mechanisms; otherwise, the mechanism is considered unresolved (Figure 3a).

**Figure 3:**
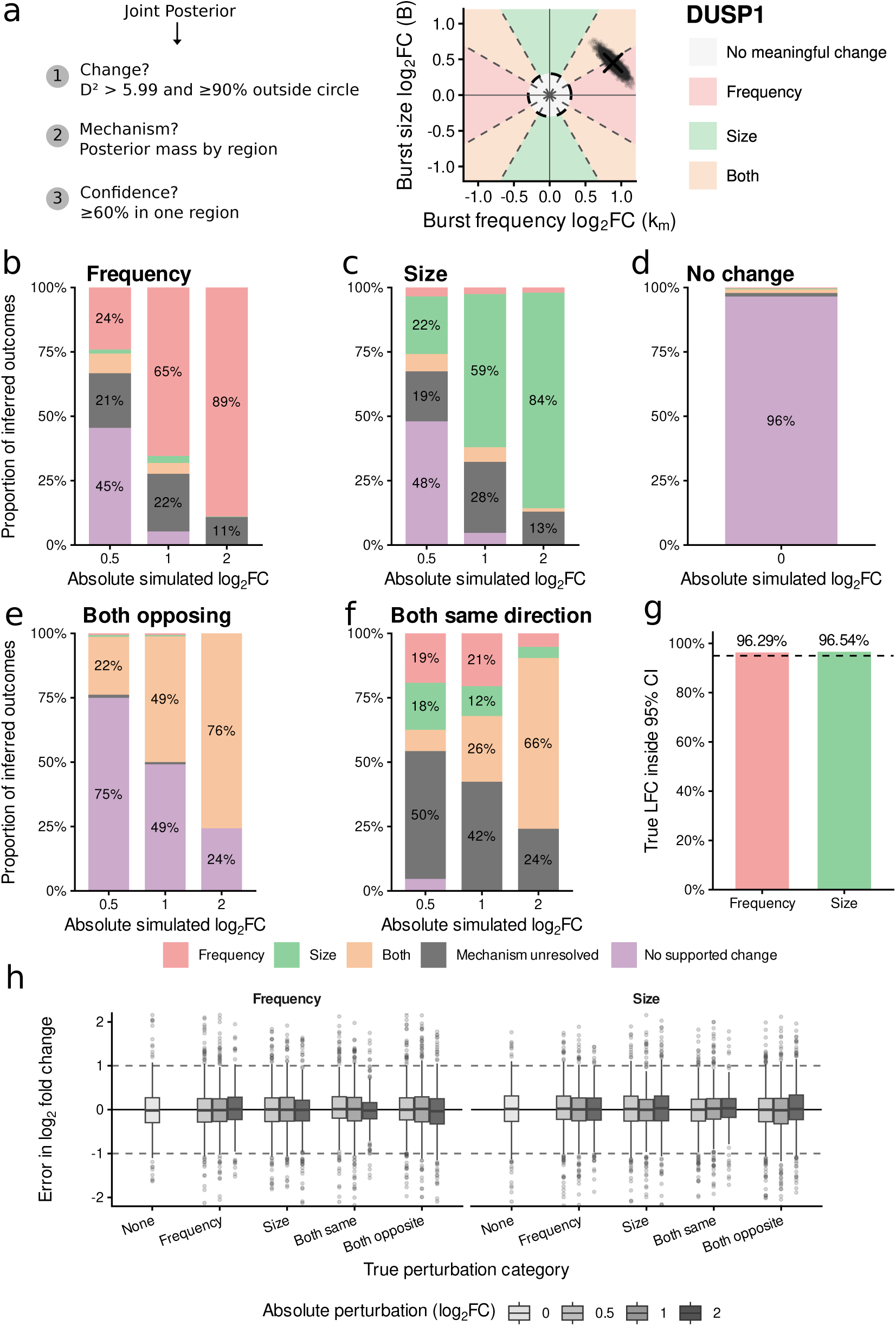
Validation of condition-specific changes in transcriptional bursting. (**a**) Strategy used by buRst to detect and classify changes in burst frequency and burst size from the joint posterior distribution of their log_2_ fold changes. A change is supported when the squared Mahalanobis distance from the origin satisfies *D*^2^ *>* 5.99 and at least 90% of the posterior mass lies outside the region of practical equivalence. Mechanistic assignments are based on posterior mass in the corresponding regions and are considered confident when at least 60% of posterior draws outside the no-change circle support one mechanistic region. (**b**) Classification outcomes for simulated burst-frequency-only perturbations of increasing absolute magnitude. Bars show the fraction of genes classified as frequency, size, both, mechanistically unresolved, or no supported change. (**c**) Corresponding classification outcomes for burst-size-only perturbations. (**d**) Classification outcomes in the absence of an imposed change. (**e**) Classification outcomes for balanced perturbations in which burst frequency and burst size changed by equal magnitudes in opposite directions. (**f**) Classification outcomes for balanced perturbations in which burst frequency and burst size changed by equal magnitudes in the same direction. (**g**) Empirical coverage of the nominal 95% approximate posterior credible intervals for burst-frequency and burst-size log_2_ fold changes. The dashed line denotes 95% nominal coverage; observed coverage was 96.29% for burst frequency and 96.54% for burst size. (**h**) Errors in burst-frequency and burst-size log_2_ fold changes across true perturbation categories and strengths, calculated as estimated minus true log_2_ fold changes. The solid horizontal line indicates exact recovery; dashed lines at *±*1 indicate twofold over- or underestimation of the true fold change.

To evaluate classification independently of biological correlations, we generated an artificial perturbation benchmark using kinetic parameters inferred from the sci-fate 0 h condition as reference values. Burst frequency *k_m_*, burst size *B*, or both were perturbed by predefined log_2_ fold changes, while degradation rates remained unchanged. The perturbed condition was simulated as a non-steady-state response initialized from the reference population, followed by parameter estimation and differential classification. Known kinetic changes allowed us to assess quantitative recovery and classification across perturbation strengths.

For frequency-only perturbations, correct classification increased from 24% at an absolute perturbation of 0.5 log_2_-fold to 65% at 1 and 89% at 2 (Figure 3b). For size-only perturbations, the corresponding fractions were 22%, 59%, and 84% (Figure 3c). Cross-mechanism assignments were uncommon; weak perturbations were generally unresolved or unsupported. Consistent with this conservative behavior, 96% of genes without an imposed change were classified as showing no supported change (Figure 3d).

For balanced perturbations in opposite directions, combined changes were correctly identified in 22%, 49%, and 76% of genes at absolute perturbations of 0.5, 1, and 2 log_2_-fold, respectively (Figure 3e). Weak perturbations were predominantly unsupported rather than assigned to an incorrect single mechanism. Balanced same-direction perturbations were also increasingly identified as combined changes, reaching 66% at 2 log_2_-fold (Figure 3f). Weaker changes often remained unresolved or were assigned to the component with the stronger estimated change. Limited identifiability thus primarily reduced resolution of weak joint perturbations.

Joint estimates showed the expected compensatory relationship arising from *λ* = *k_m_B*: deviations in one parameter can be offset by deviations in the other, producing anticorrelated errors. Nevertheless, estimates tracked the imposed changes, while uncertainty remained elongated along the compensatory frequency–size direction (Supplementary Figure S3a). The position of the joint distribution therefore reflected the perturbation, whereas its shape reflected the intrinsic identifiability tradeoff.

Despite this coupling, fold changes were recovered accurately across perturbation categories and strengths, with estimated-minus-true log_2_ fold-change distributions centered near zero (Figure 3h). The median absolute error was 0.26 log_2_ units for both parameters, corresponding to a 1.20-fold error. For 90% of genes, absolute errors were below 0.88 and 0.89 log_2_ units for burst frequency and burst size, respectively, corresponding to 1.84-fold and 1.86-fold errors. Empirical coverage of the nominal 95% approximate posterior credible intervals was 96.29% for burst frequency and 96.54% for burst size (Figure 3g).

Together, these simulations demonstrate that buRst recovers and mechanistically classifies non-steady-state changes in bursting. Classification became more specific with stronger perturbations, whereas weak or poorly identifiable changes were generally unresolved or unsupported. The frequency–size tradeoff limits mechanistic resolution but is reflected in the inferred uncertainty.

### The sci-fate response involves coordinated changes in burst frequency and burst size

We next applied buRst to compare the sci-fate 0 h and 2 h conditions, focusing on early dexamethasone responders with a log_2_ fold change in new RNA abundance greater than one.

Of the 69 analyzed upregulated genes, the mechanism remained unresolved for 15 (21.7%). Among the remaining 54 genes, 30 (55.6%) were classified as frequency changes, 8 (14.8%) as size changes, and 16 (29.6%) as combined changes (Figure 4a,b; Supplementary Figure S4). Of the 16 genes with combined changes, 14 showed compensatory increases in burst frequency and decreases in burst size. Thus, among early responders with a resolved mechanism, burst-frequency changes were most common.

**Figure 4:**
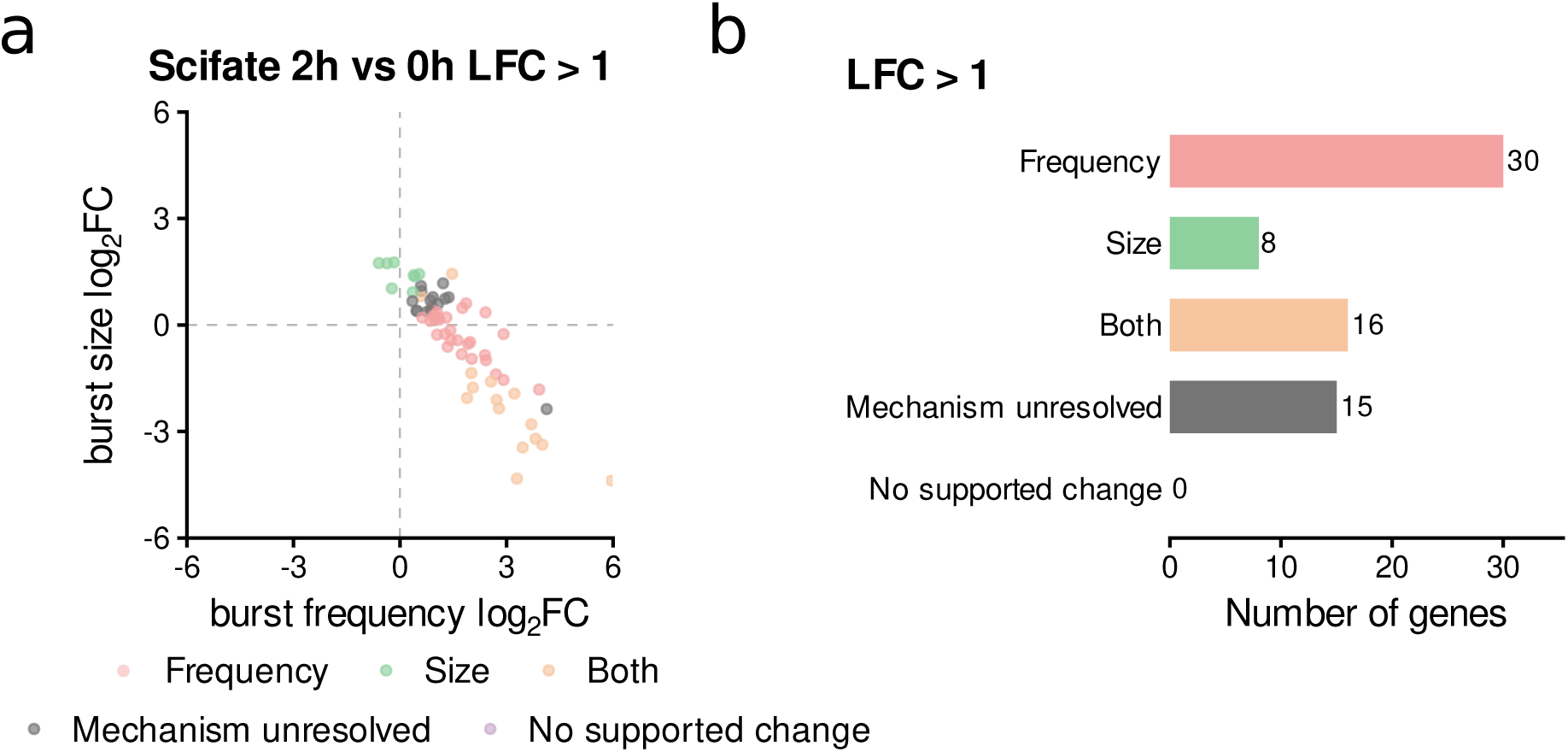
Transcriptional bursting changes between the sci-fate 0 h and 2 h conditions. (**a**) Joint log_2_ fold changes in burst frequency and burst size for genes with a log_2_ fold change in new RNA abundance greater than one (*n* = 69). (**b**) Numbers of genes in each mechanistic class.

### Acute MYC depletion induces distinct transcriptional responses in melanoma cells

To investigate how acute modulation of transcriptional output affects burst frequency and burst size, we focused on MYC, given its central role in tumorigenesis and the longstanding interest in its molecular mechanisms of action. We established an inducible degradation system for acute and reversible manipulation of endogenous MYC in two human melanoma cell lines with distinct basal MYC expression: A375, with high expression, and MaMel63, with comparatively low expression. An auxin-inducible degron (AID) tag was introduced into the endogenous MYC locus in both cell lines by genome editing [32], followed by stable lentiviral expression of TIR1 to enable auxin-dependent ubiquitination and degradation of AID-tagged MYC.

Cells were treated with DMSO or auxin for 2 h before addition of 4sU, followed by a further 2 h of metabolic labeling and subsequent scSLAM-seq profiling (Figure 5a). In separate immunoblot experiments, MYC protein was strongly depleted in both cell lines by 3 h of auxin treatment and remained low at 6 h (Figure 5b).

**Figure 5:**
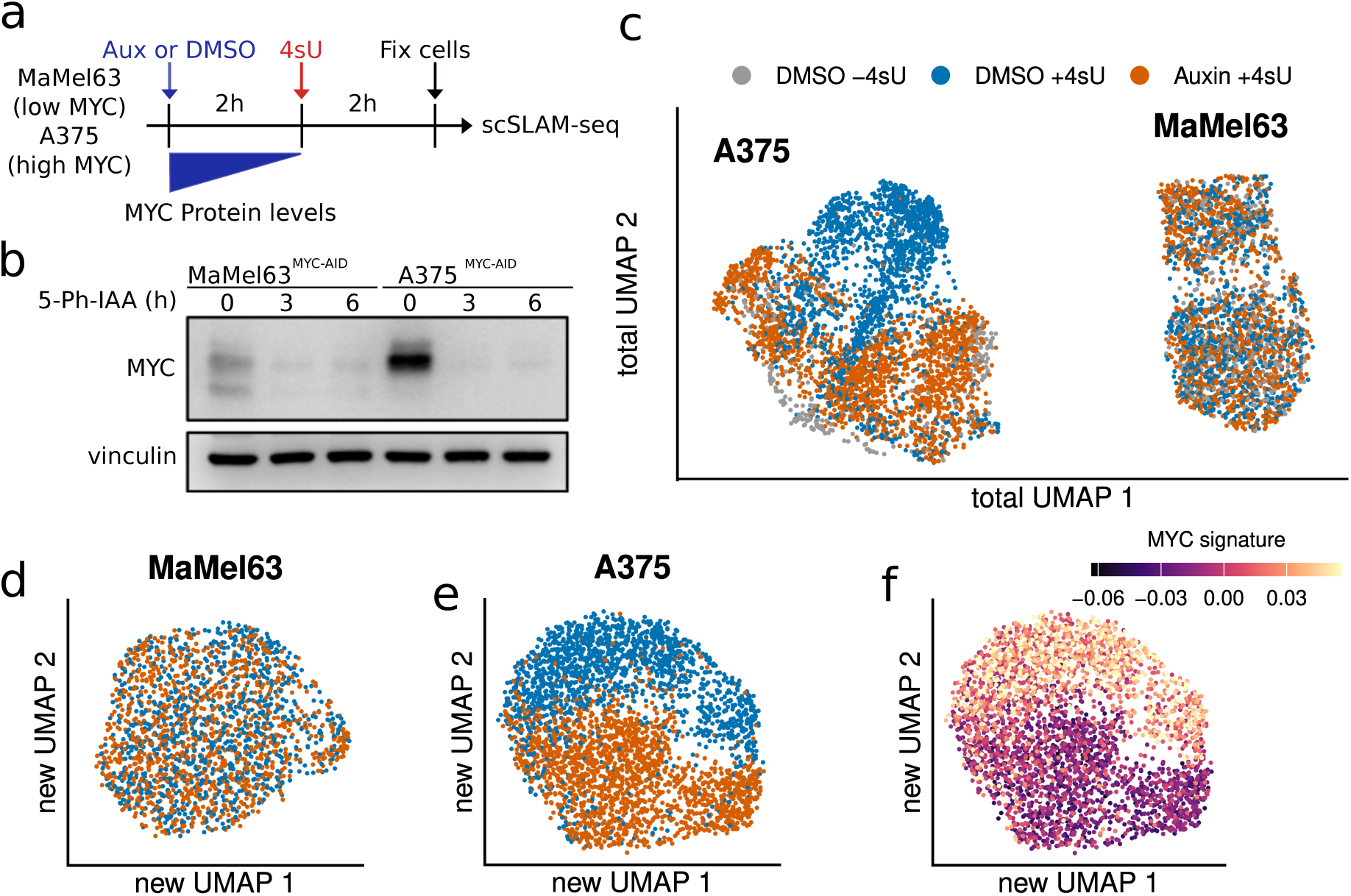
scSLAM-seq profiling of MYC-dependent transcriptional responses in MaMel63 and A375 cells. (**a**) Experimental design. MaMel63 and A375 melanoma cells were treated with auxin or DMSO for 2 h, labeled with 4sU for 2 h, and profiled by scSLAM-seq following fixation. (**b**) Immunoblot analysis of MYC protein levels in MaMel63^MYC^*^−^*^AID^ and A375^MYC^*^−^*^AID^ cells after 0, 3, and 6 h of 5-Ph-IAA treatment. Vinculin served as a loading control. (**c**) Total RNA UMAP embedding of all retained cells, colored by treatment and labeling status. (**d,e**) New RNA UMAP embeddings of MaMel63 (**d**) and A375 (**e**) cells, colored by treatment and labeling status. (**f**) MYC target-gene signature score projected onto the A375 new RNA UMAP embedding.

MULTI-seq demultiplexing assigned 7,044 cells. Increased exonic T-to-C mismatch frequencies in labeled samples relative to no-4sU controls, together with GRAND3 estimates separating labeling-associated conversions from background mismatches, confirmed successful labeling (Supplementary Figure S5a–d). Quality control excluded 657 cells from low-quality populations in total or new RNA embeddings and 122 with transcriptomic profiles inconsistent with their assigned cell line, leaving 6,265 cells for downstream analyses. Across retained samples, median total RNA counts ranged from 14,106 to 38,192 UMIs and median detected genes from 4,453 to 6,754 per cell; median mitochondrial RNA fractions ranged from 4.9% to 10.5% (Supplementary Figure S5).

UMAP embeddings of total RNA primarily separated cells by cell-line identity (Figure 5c). DMSO- and auxin-treated MaMel63 cells largely overlapped, whereas A375 cells showed partial treatment-associated separation. No-4sU controls overlapped with corresponding labeled DMSO-treated cells, suggesting that labeling did not dominate the global transcriptional structure.

To focus on acute transcriptional responses, we constructed embeddings from newly synthesized RNA. MaMel63 cells remained broadly intermixed across treatments, whereas A375 cells showed clear separation (Figure 5d,e). Treatment-associated states were therefore more distinguishable in A375, although embedding separation alone does not quantify the magnitude or number of transcriptional changes.

We assessed MYC-dependent transcription using a published 100-gene signature [26]. In the A375 new RNA embedding, scores were lower in the auxin-enriched region and higher in the DMSO-enriched region, consistent with reduced MYC-dependent transcription following depletion (Figure 5f).

Together, these analyses reveal cell-line-specific transcriptional responses to acute MYC depletion, with a distinct new RNA state and reduced MYC signature activity in A375 cells. This dataset provides a system for testing whether buRst resolves condition-dependent transcriptional responses at single-cell resolution.

### Pseudobulk analysis links MYC-dependent transcription to quantitative promoter binding

To assess MYC’s role as a general amplifier or selective activator, we jointly analyzed published MYC perturbation data from K562 and HCT116 cells [26] and our pseudobulk MaMel63 and A375 data. The top 100 MYC response genes showed higher transcription with MYC present in all four cell lines (median log_2_ fold changes: A375, 1.17; MaMel63, 0.52; K562, 2.03; HCT116, 1.02; Figure 6a). The stronger response in A375 than MaMel63 was consistent with its higher endogenous MYC levels. These genes were previously considered representative selective MYC targets [26].

**Figure 6:**
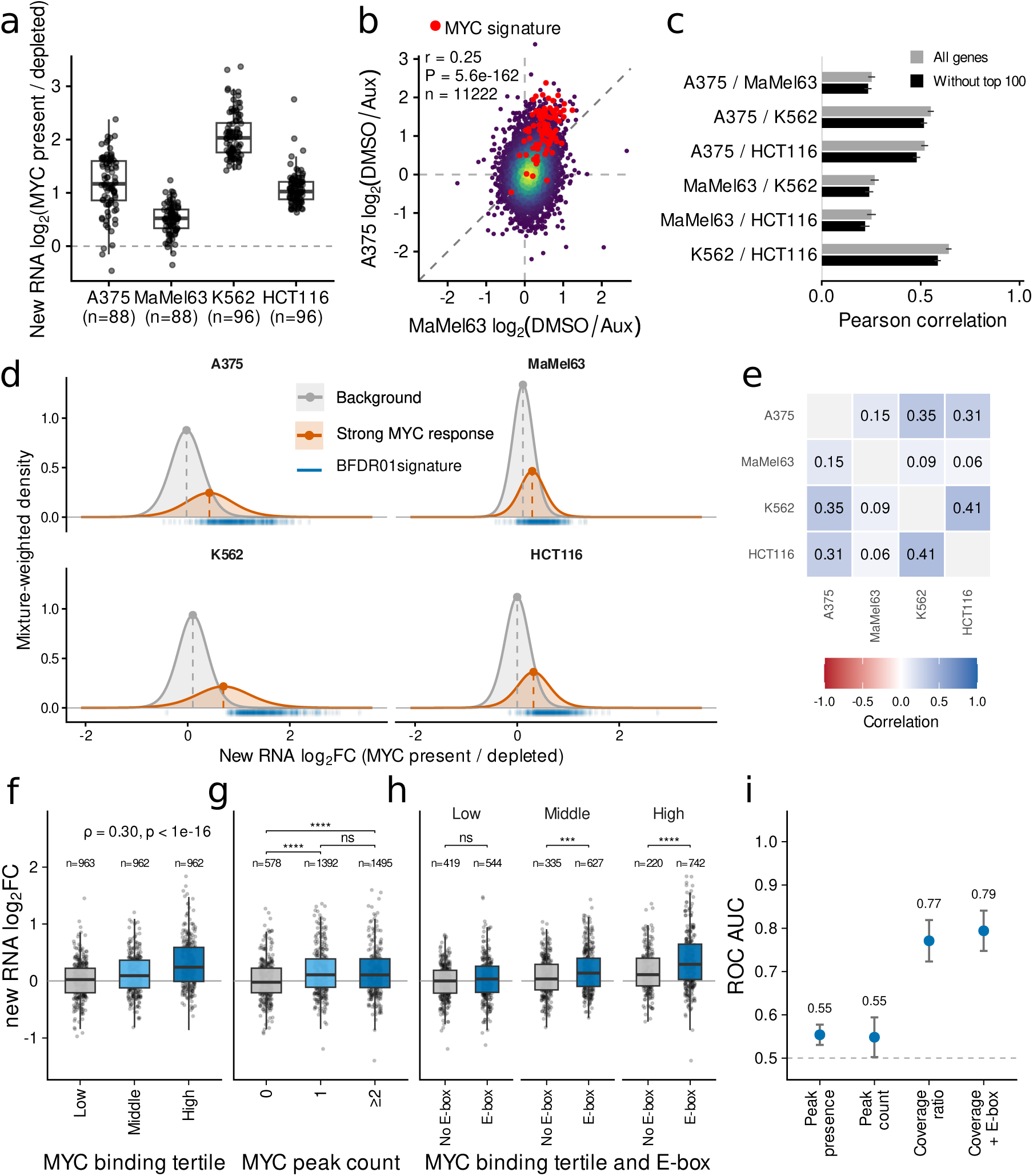
Pseudobulk analysis of acute MYC-dependent transcriptional responses and promoter-associated MYC binding. (**a**) New RNA log_2_(MYC present*/*depleted) fold changes for the 100-gene MYC-response signature in A375, MaMel63, K562, and HCT116 cells. A375 and MaMel63 data are from this study; K562 and HCT116 data were re-analyzed from published datasets [26]. Points represent signature genes; boxes indicate medians and interquartile ranges. Numbers below boxes indicate the number of quantified signature genes. (**b**) Comparison of pseudobulk new RNA log_2_(DMSO*/*Aux) fold changes between MaMel63 and A375 cells. MYC-response signature genes are highlighted [26]. Annotations report Pearson’s correlation coefficient (*r*), *P* value, and gene count (*n*). (**c**) Pearson correlations of new RNA responses for all six cell-line pairs, calculated using all eligible genes (grey) or after excluding the Muhar top-100 signature genes (black). Error bars show 95% confidence intervals. Each comparison uses the genes available for that cell-line pair. (**d**) Marginal distributions of new RNA responses in each cell line from a jointly fitted two-component, four-dimensional multivariate *t* mixture with component-specific scale matrices and degrees of freedom. Grey and orange curves show mixture-weighted densities for the background and strong MYC-response components, respectively. Dashed lines and points mark component means. Blue ticks mark responses of genes selected by the joint model at 1% Bayesian FDR (BFDR01; *n* = 438). (**e**) Cross-cell-line correlations derived from the separately estimated covariance matrix of the near-zero component of the same *t* mixture. Values are fitted component correlations. (**f**) New RNA response across tertiles of quantitative MYC promoter binding, defined by log_10_ MYC/Input coverage among genes with assigned MYC peaks. Spearman’s *ρ* and *P* value describe the association between quantitative MYC binding and new RNA response. (**g**) New RNA response by promoter-associated MYC-peak count. (**h**) New RNA response for promoters with and without an E-box across low, intermediate, and high MYC-binding tertiles, restricted to genes with assigned promoter MYC peaks. Numbers indicate gene counts. For **g** and **h**, two-sided Wilcoxon rank-sum tests used Benjamini–Hochberg correction across displayed comparisons within each panel. Adjusted *P* values: ns, *P ≥* 0.05; *∗*, *P <* 0.05; *∗∗*, *P <* 0.01; *∗ ∗ ∗*, *P <* 0.001; *∗ ∗ ∗∗*, *P <* 0.0001. (**i**) ROC AUC for classifying strongly MYC-dependent genes (new RNA log_2_(DMSO*/*Aux) *≥* 1) using promoter-associated MYC-peak presence, peak count, or MYC-to-input coverage ratio. Quantitative occupancy and E-box presence were combined by logistic regression with predicted probabilities from stratified five-fold cross-validation. Points indicate ROC AUC estimates; error bars, 95% confidence intervals; dashed line, chance performance (ROC AUC = 0.5).

To further challenge the selective activator hypothesis, we examined whether responses to MYC depletion were shared more broadly across genes. Newly synthesized and total RNA yielded largely similar results, although total RNA preferentially reflects responses of short-lived transcripts (Supplementary Figure S6a). New RNA responses correlated between A375 and MaMel63 (*r* = 0.25, *P* = 5.6 *×* 10*^−^*^162^, *n* = 11,222 genes; Figure 6b), with stronger responses in A375. Despite differences in cell type and protocol, A375 responses also correlated with K562 (*r* = 0.55) and HCT116 (*r* = 0.52); correlations were weaker for MaMel63 (*r* = 0.27 and *r* = 0.25, respectively; Supplementary Figure S6b). These associations persisted after excluding the Muhar top-100 signature genes: the A375–MaMel63 correlation remained *r* = 0.23 (*n* = 11,134), and correlations with K562 and HCT116 remained *r* = 0.52 and *r* = 0.48 for A375 and *r* = 0.24 and *r* = 0.22 for MaMel63, respectively (all *P <* 10*^−^*^74^; Figure 6c). Thus, shared MYC-dependent responses extended beyond the established top-100 signature.

To define a shared strong-response signature, we modeled new RNA responses across all four cell lines using a two-component, four-dimensional multivariate *t* mixture with component-specific scale matrices and degrees of freedom (Figure 6d). This model accommodates heavy-tailed response distributions. Selection at a Bayesian false discovery rate (BFDR) of 1% yielded a 438-gene signature, termed BFDR01. We interpret this signature as an operationally defined strongly responsive subset rather than a natural boundary between MYC-dependent and independent genes. Cross-cell-line correlations remained positive within the near-zero component (Pearson’s *r* = 0.06–0.41; Figure 6e).

We next related transcriptional responses to promoter-associated MYC binding using A375 MYC ChIP-seq and matched input data [32]. Among genes with an assigned promoter peak, increasing occupancy, quantified as the log_10_ MYC-to-input signal ratio, was associated with stronger responses to MYC depletion (*ρ* = 0.30, *P <* 10*^−^*^16^; Figure 6f).

Genes with promoter-associated MYC peaks responded more strongly than genes without peaks, but multiple peaks were not associated with an additional response (Figure 6g). Median responses were *−*0.02, 0.11, and 0.11 log_2_(DMSO*/*Aux) for genes with no peak, one peak, or at least two peaks, respectively. Peak presence was therefore informative, whereas peak number added little information.

We next asked whether E-box-containing promoters responded more strongly within MYC-binding tertiles. E-box-containing promoters showed significantly higher responses at intermediate and high occupancy, whereas no significant difference was detected at low occupancy (Figure 6h). Promoter sequence context therefore provided information beyond quantitative occupancy.

Finally, we assessed prediction of strongly MYC-dependent genes, defined by a new RNA response of at least 1 log_2_(DMSO*/*Aux). Peak presence and count showed modest classification performance (AUROC = 0.55 for both), whereas quantitative MYC-to-input coverage performed better (AUROC = 0.77; Figure 6i). Adding E-box presence in a five-fold cross-validated logistic-regression model modestly increased AUROC to 0.79. Consistent with previous studies [26, 33], quantitative binding was more informative than binary peak classification for predicting acute MYC dependence.

Together, these results support shared MYC-dependent responses extending beyond a strongly responsive gene subset, consistent with general amplification. Response magnitude was associated with quantitative promoter occupancy and E-box context, and stronger responses in A375 than MaMel63 coincided with higher endogenous MYC levels.

### Acute MYC depletion induces coordinated changes in burst size and burst frequency

Bulk or pseudobulk analyses of responses to MYC depletion can only quantify net changes in mean RNA synthesis, leaving the underlying changes in burst frequency and burst size unresolved. We used buRst to identify kinetic mechanisms underlying acute MYC depletion responses in A375. Because MYC-dependent new RNA responses were broadly similar across cell-cycle phases among the 397 evaluable BFDR01 signature genes per cell line, with absolute median paired differences below 0.11 log_2_ units, we pooled cells across phases for bursting analyses (Supplementary Figure S7).

In A375, 2,116 of 3,485 analyzed genes showed no supported kinetic change, 1,077 changed both burst size and frequency, 77 changed frequency, and 41 changed size. For 174 genes, a change was supported but the mechanism remained unresolved (Figure 7a,b).

**Figure 7:**
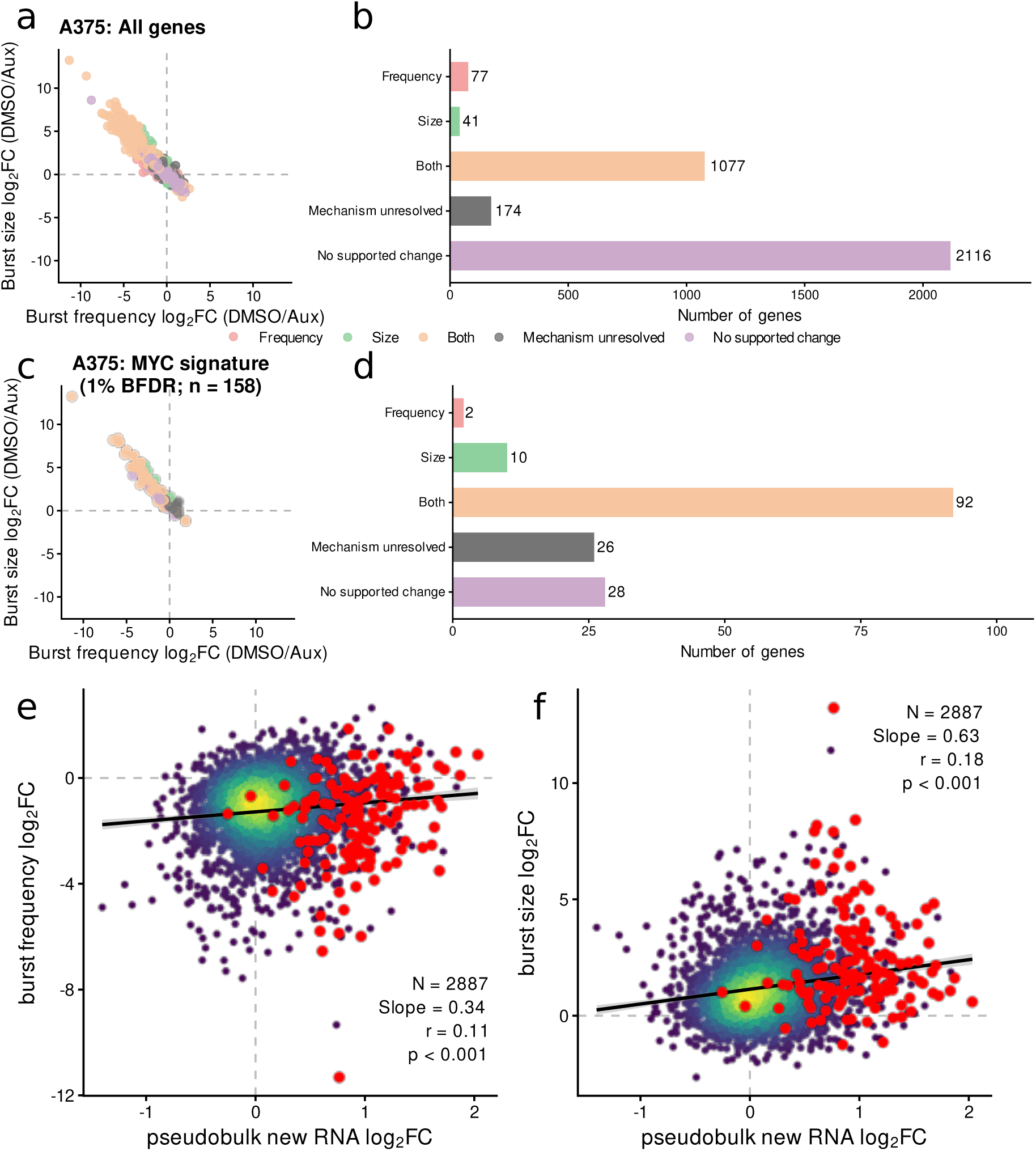
MYC-dependent changes in transcriptional bursting kinetics. All fold changes are expressed as log_2_(DMSO*/*Aux). (**a**,**b**) Joint burst-frequency and burst-size fold changes and mechanistic class counts for all analyzed A375 genes (*n* = 3,485). Colors indicate inferred mechanisms; dashed lines indicate no change. (**c**,**d**) Corresponding plots for the 158 BFDR01 signature genes with evaluable comparisons. (**e**,**f**) Pseudobulk new RNA responses versus burst-frequency (**e**) or burst-size (**f**) responses for *n* = 2,887 genes with finite responses in all three measurements and a finite, positive promoter MYC-to-input coverage ratio. Colors indicate local point density; BFDR01 signature genes are highlighted in red. Black lines and shading show linear regression fits and 95% confidence intervals. Annotations report slopes, Pearson’s *r*, and corresponding *P* values.

Among 158 BFDR01 signature genes with evaluable bursting comparisons in A375, 92 changed both parameters, 10 changed size, and 2 changed frequency; 26 were unresolved and 28 showed no supported change (Figure 7c,d). Unlike the sci-fate comparison, where the strongest expression changes were more frequently associated with burst frequency, supported changes in MYC-signature genes predominantly involved both parameters.

MaMel63 showed a similar inverse relationship between burst size and frequency, with combined changes predominating among genes with supported changes (Supplementary Figure S8a,b). Among 251 BFDR01 signature genes, 108 changed both parameters, 25 changed size alone, and 1 changed frequency alone; 22 were unresolved and 95 showed no supported change (Supplementary Figure S8c,d). Thus, coordinated changes occurred in both cell lines.

The net changes in new RNA synthesis were more strongly associated with burst-size changes (slope = 0.63, 95% CI [0.51, 0.76], *r* = 0.18) than burst-frequency changes (slope = 0.34, 95% CI [0.23, 0.46], *r* = 0.11; Figure 7e,f). Although modest, these associations suggest that acute MYC-dependent transcriptional responses are more closely linked to burst size, often accompanied by coordinated frequency changes. Across the 2,887 genes included in this analysis, the presence of MYC was associated with globally increased burst sizes (mean log_2_ fold change: 1.24) and compensatory decreased burst frequencies (mean log_2_ fold change: *−*1.23).

### Quantitative MYC binding is preferentially associated with burst-size changes

To determine how promoter-associated MYC binding relates to acute MYC dependence, we analyzed its association with response magnitude and kinetic mechanism. Quantitative promoter-associated MYC occupancy was positively associated with the new RNA log_2_(DMSO*/*Aux) response (Figure 8a; slope = 0.551, 95% CI [0.493, 0.609], *R*^2^ = 0.108, *P <* 2 *×* 10*^−^*^16^). Stronger binding therefore corresponded to greater reductions in RNA synthesis after MYC depletion, although it explained only part of the variation.

**Figure 8:**
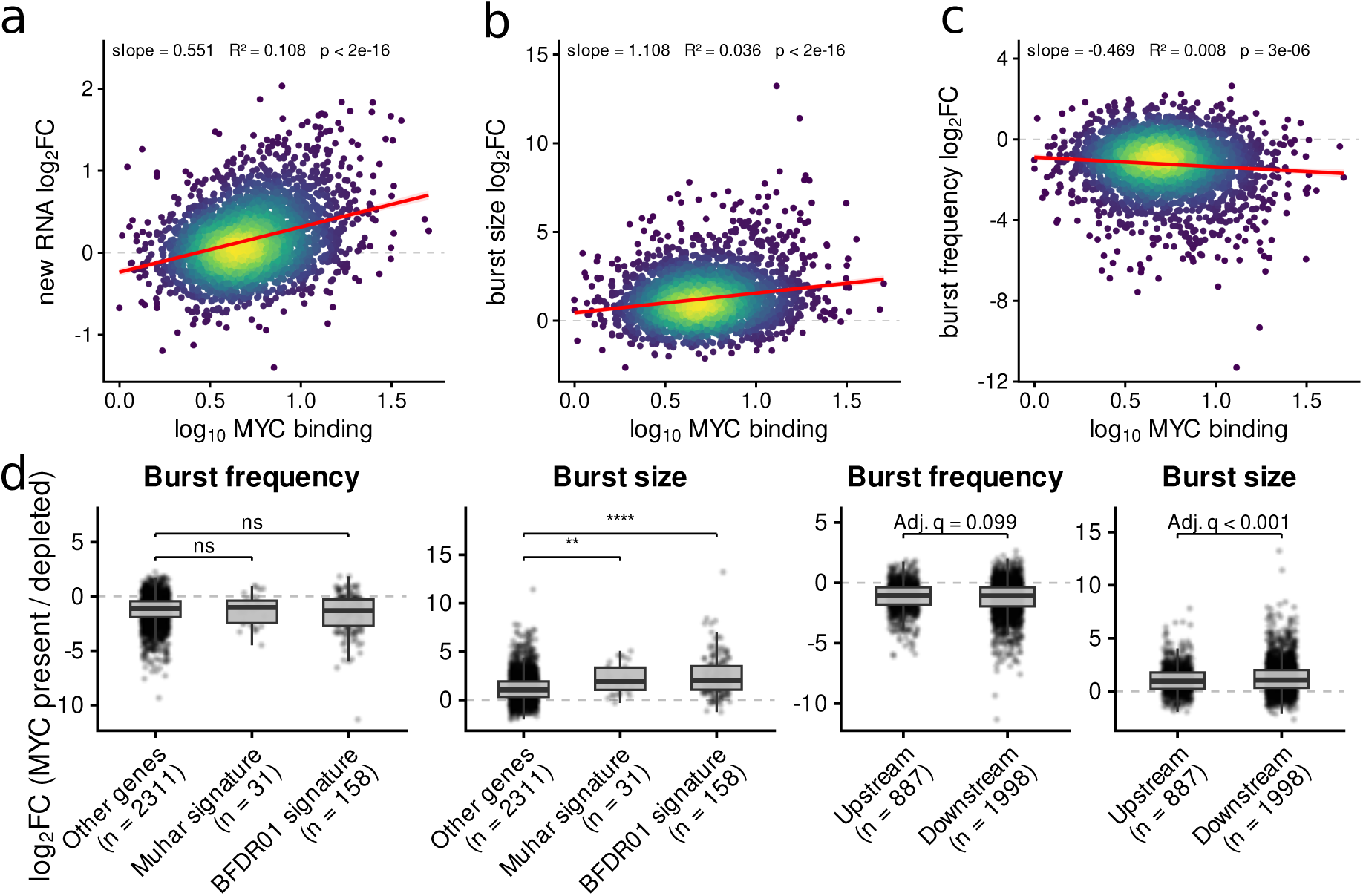
Association of quantitative MYC binding with transcriptional bursting changes. (**a**) Relationship between quantitative promoter-associated MYC binding and the pseudobulk new RNA log_2_(DMSO*/*Aux) fold change. (**b**,**c**) Relationship between MYC binding and the inferred log_2_(DMSO*/*Aux) fold change in burst size (**b**) or burst frequency (**c**). Quantitative MYC binding is shown as the log_10_ MYC-to-input promoter coverage ratio. All three panels use the same *n* = 2,887 genes with finite responses for all three measurements and a finite, positive coverage ratio. Point color indicates local density. Red lines show ordinary least-squares regression fits, and shaded regions indicate 95% confidence intervals. Annotations report the fitted slope, the coefficient of determination (*R*^2^), and the *P* value testing whether the slope differs from zero. (**d**) Burst-frequency and burst-size log_2_(MYC present*/*depleted) fold changes for other genes, genes belonging to the Muhar signature, and genes belonging to the BFDR01 signature (left), and for genes with upstream or downstream MYC peaks (right). Other genes exclude both signatures; the two signature groups overlap. Boxes indicate the median and interquartile range, and points represent individual genes. Numbers below the categories indicate the number of genes. Signature-group comparisons used two-sided Wilcoxon rank-sum tests against other genes, with Benjamini–Hochberg correction across the four comparisons. For upstream-versus-downstream comparisons, annotations show adjusted *P* values for the peak-position coefficient from multivariable linear models controlling for quantitative MYC binding, E-box presence, TATA-box presence, CpG-island overlap, and absolute peak-to-TSS distance. Benjamini–Hochberg correction was applied jointly across the 18 non-intercept coefficients from the new RNA, burst-size, and burst-frequency models. For the signature panels, significance symbols indicate adjusted *P* values: ns, *P ≥* 0.05; *∗*, *P <* 0.05; *∗∗*, *P <* 0.01; *∗ ∗ ∗*, *P <* 0.001; *∗ ∗ ∗∗*, *P <* 0.0001.

MYC occupancy was also positively associated with burst-size responses (Figure 8b; slope = 1.108, 95% CI [0.900, 1.317], *R*^2^ = 0.036, *P <* 2 *×* 10*^−^*^16^). Burst-frequency responses showed a weaker and negative association (Figure 8c; slope = *−*0.469, 95% CI [*−*0.666*, −*0.272], *R*^2^ = 0.008, *P* = 3 *×* 10*^−^*^6^). Thus, the positive association of MYC binding with the net transcriptional response was accompanied by increasing burst size, whereas burst frequency showed an opposing association.

Burst-size responses were broadly positive even outside the MYC signatures, consistent with a general amplification of burst sizes by MYC (Figure 8d). Both Muhar and BFDR01 signature genes showed larger burst-size responses than other genes, whereas burst-frequency responses did not differ significantly. When MYC peaks were located downstream of the transcription start site, they were associated with modestly larger burst-size responses after accounting for quantitative binding and other promoter features, whereas the association with burst-frequency responses was not significant (Supplementary Figure S9). In conclusion, MYC globally amplifies burst size in a manner that scales with promoter occupancy and is modulated by promoter positioning, whereas burst frequency decreases in an opposing manner.

## 3 Discussion

Inferring transcriptional bursting kinetics from scSLAM-seq is challenging, since the identifiability of burst size and frequency is strongly dependent on an accurate estimation of the true variability of newly synthesized RNA across individual cells. Thus, it is essential to separate estimation uncertainty from true biological variance. buRst accomplishes this by marginalizing over the likelihood profile of new RNA counts per cell. Reducing the labeling evidence to point estimates or beta approximations distorted this variability, resulting in biased kinetic estimates and substantial interval undercoverage. In contrast, the full labeling likelihood recovered the generating parameters with little systematic bias and close-to-nominal uncertainty coverage. Explicitly accounting for labeling uncertainty is therefore required not only for reliable inference of transcriptional activity, but also for resolving the kinetic mechanisms underlying differences in RNA production.

While steady-state transcript count distributions are well characterized for common transcriptional bursting models, calculating transient distributions under non-steady-state conditions is more challenging. The steady-state assumption is inappropriate for acute perturbations such as drug treatment or auxin-induced degradation of a regulatory factor. To address this, we developed a mathematical approach to numerically evaluate the time-dependent probability distribution for a given initial condition and set of kinetic parameters using the fast Fourier transform, as well as an approximation based on moments. Using these numerically evaluated distributions for likelihood calculations, we were able to infer transient changes in bursting kinetics directly, rather than interpreting them as differences between two independent steady states.

We first applied our approach to the dexamethasone-induced response in the sci-fate dataset [13]. Dexamethasone activates the glucocorticoid receptor (GR), which then translocates to the nucleus and binds to glucocorticoid response elements in the genome to induce hundreds of genes transcriptionally [34]. We found that the immediate transcriptional responders were predominantly regulated through burst frequency. This frequency-dominated response is consistent with previous studies identifying promoter activation frequency as an important target of transcriptional regulation. Genome-wide inference of mammalian bursting kinetics showed that enhancer activity mainly modulates burst frequency, whereas core-promoter architecture has a stronger influence on burst size [7]. Likewise, experimental manipulation of transcription-factor binding in a controlled mammalian system primarily altered burst frequency while leaving burst size largely unchanged [35]. Together, these observations establish modulation of burst frequency as a recurrent mode through which transcriptional regulatory inputs can alter gene expression.

While modulation of burst frequency is a common regulatory paradigm for transcription factors, the response to MYC presents a strikingly different and more complex picture. MYC plays a central role in cellular proliferation and metabolism, and its transcriptional targets include genes involved in cell growth, cell cycle progression, ribosome biogenesis, and metabolism [36–38]. MYC is also an important oncogene, and its dysregulation is a hallmark of many cancers. Despite the large body of work on MYC, its exact role in transcriptional regulation still remains debated: MYC has been described to act as a selective transcriptional regulator, or as a global amplifier of transcription. Early studies proposed that increased MYC broadly amplifies the output of genes that are already transcriptionally active [21, 22], whereas subsequent work emphasized selective regulation of MYC target genes and argued that global increases in RNA output can arise indirectly from MYC-induced changes in cellular state [18, 24, 25]. The non-steady-state formulation of buRst allowed us to investigate this long-standing question from a new perspective, suggesting that these apparently opposing models can be reconciled at the level of bursting kinetics. Our shared BFDR01 signature identifies genes with strong, consistent MYC-dependent responses across four cell lines [26]. We interpret this signature as a strongly responsive subset rather than an exclusive set of selective MYC targets: correlations between cell lines persisted after excluding the Muhar top-100 signature genes, and positive correlations remained within the near-zero component of the multivariate *t* mixture. Together, these observations support an interpretation of graded MYC dependence. Quantitative promoter occupancy was associated with response magnitude, suggesting that the strength of MYC dependence varies with promoter-associated binding.

In stark contrast to the GR response, MYC-dependent transcriptional activation was reflected predominantly in the effective burst size, consistent with live-cell measurements of a handful of genes showing that increased MYC prolongs productive transcriptional bursts without increasing their frequency [27]. Consistent with this interpretation, both the Muhar and BFDR01 signatures showed larger inferred burst-size responses than other genes, whereas no significant differences in burst-frequency responses were detected. Our data extend this observation across the transcriptome, revealing a global increase in burst size, with the magnitude of amplification further scaling with promoter-associated MYC occupancy. Thus, in contrast to the predominantly frequency-mediated response observed after dexamethasone treatment and to regulatory systems in which TF binding primarily alters promoter activation frequency [35], MYC acts mainly on the productivity of transcriptionally active periods. Importantly, this behavior is not restricted to a set of selective targets, but applies broadly across genes depending on the binding affinity of MYC to promoters. Promoter context also shaped MYC dependence. At comparable MYC occupancy, E-box-containing promoters showed stronger transcriptional responses, suggesting more productive binding and complementing previous work on sequence specificity and promoter affinity [33, 39]. While the kinetic basis of this E-box-associated response remained unresolved, downstream MYC peaks were associated with modestly larger burst-size responses after accounting for occupancy and other promoter features.

In contrast to the occupancy-dependent increase in burst size, burst frequency decreased globally in the presence of MYC and showed only a weak relationship with local MYC occupancy. We therefore propose that this reduction is not primarily a direct promoter-specific effect of MYC, but a consequence of the increased productivity of MYC-dependent transcriptional bursts. If MYC prolongs or intensifies productive transcriptional periods, RNA polymerase II or other limiting factors required for transcription initiation may remain engaged for longer at highly active promoters. Their increased sequestration at productive transcription sites could consequently reduce their availability for initiating new bursts elsewhere in the genome, resulting in a global reduction in burst frequency.

Such a mechanism would provide a direct buffering effect on MYC-driven transcriptional amplification. At strongly MYC-bound promoters, the increase in burst size is sufficiently large to compensate for the accompanying reduction in burst frequency, resulting in a net increase in transcriptional output. In contrast, genes receiving little or no MYC-dependent burst-size amplification would still experience the global decrease in burst frequency and could consequently become transcriptionally downregulated. At least part of the gene repression observed in the presence of MYC may therefore represent an indirect consequence of MYC activity rather than direct transcriptional repression, arising from competition for limiting components of the transcriptional machinery.

Together, these findings support a model in which MYC acts as an affinity-dependent global amplifier of transcription in the literal sense: it increases the amplitude but not the frequency of transcriptional bursting. Instead, this occupancy-dependent amplification is accompanied by a genome-wide reduction in burst frequency, which buffers MYC-driven increases in transcriptional output across genes and can indirectly reduce transcription at genes receiving little burst-size amplification. In this view, selective MYC dependence and global transcriptional amplification are not mutually exclusive: MYC globally increases transcriptional productivity, but the magnitude of this effect is quantitatively encoded by local promoter occupancy.

## 4 Methods

### Experimental procedures

#### Generation of MYC-AID knock-in cells

An auxin-inducible degron (AID) system was established in MaMel63 melanoma cells [40] to enable conditional degradation of endogenous MYC. The homology-directed repair (HDR) template was assembled in pJET (Thermo Fisher Scientific) by inserting the AID-V5-P2A-Blasticidin cassette between PCR-amplified homology arms corresponding to the genomic sequences flanking the *MYC* stop codon. An sgRNA targeting the *MYC* stop codon region was cloned into PX458 (Addgene, #48138). Both plasmids were co-transfected into MaMel63 cells using Lipofectamine 2000 (Thermo Fisher Scientific). After 72 h, cells were selected with blasticidin (5 *µ*g*/*mL; InvivoGen) for 15 days. Single-cell-derived colonies were screened by PCR using primers binding outside the homology arms, and PCR products were analyzed by agarose gel electrophoresis to assess the integration status of the *MYC* locus. Correct knock-in was confirmed by Sanger sequencing (LGC Genomics), and a homozygous MaMel63 MYC-AID clone was selected.

#### Generation of TIR1^F74G^-expressing cells and induction of degradation

To confer auxin responsiveness, TIR1^F74G^ was introduced into MaMel63 MYC-AID cells by lentiviral transduction using a construct based on pRRLSin.cPPT.SFFV-IRES.WPRE. Transduced cells were selected with hygromycin (InvivoGen) for 7 days. Degradation of AID-tagged MYC was induced by treatment with 1 *µ*M 5-Ph-IAA (30-003-10; BioAcademia), and DMSO was used as the vehicle control.

A375 MYC-AID cells were generated using a similar approach, as previously described [32].

#### Sample preparation and immunoblotting

MaMel63-Myc-AID and A375-Myc-AID cells were seeded at a density of 1 *×* 10^6^ cells per 10-cm dish. The following day, cells were treated with either DMSO (vehicle control) or 1 *µ*M 5-Ph-IAA for 3 and 6 h. Cells were then lysed for 20 min at 4*^◦^*C in RIPA buffer (50 mM HEPES, pH 7.9, 140 mM NaCl, 1 mM EDTA, 1% Triton X-100, 0.1% SDS, and 0.1% sodium deoxycholate) supplemented with phosphatase inhibitor cocktails (P5726 and P0044, Sigma-Aldrich) and protease inhibitor cocktail (P8340, Sigma-Aldrich). Lysates were clarified by centrifugation, and protein concentrations were determined using a BCA assay. Equal amounts of protein were separated by Bis-Tris PAGE, transferred to PVDF membranes, and blocked with 5% (w/v) non-fat dry milk in TBS-T for 1 h at room temperature. Membranes were incubated overnight at 4*^◦^*C with anti-MYC antibody (1:1000; ab32072, Abcam), followed by an HRP-conjugated anti-rabbit IgG secondary antibody (1:7500; 1079-4347/GEHENA934, GE Healthcare). Vinculin (1:5000; V9131, Sigma-Aldrich) served as the loading control. Protein bands were detected using Immobilon Western HRP substrate (WBKLS0500, Merck Millipore) and imaged using a LAS-4000 Mini imaging system (Fujifilm).

#### MYC depletion and 4sU labeling

For final single-cell SLAM-seq experiments, 4sU concentration for each cell type was optimized using bulk SLAM-seq experiments. 200 *µ*M 4sU was used for A375 cells whereas 500 *µ*M 4sU was used for MaMel63 cells. For Myc depletion, 1 *µ*M of Auxin (5-Ph-IAA) concentration was used. Auxin was added to the cells 2 h prior to the 4sU addition. Afterwards, cells were incubated with 4sU for another 2 h for RNA metabolic labelling. MaMel63 cells were grown and seeded in RPMI media, whereas A375 cells were grown in DMEM media.

After 2 h of 4sU labelling, cells were trypsinized and were collected in 10 ml of culture media containing fetal calf serum (FCS). Cells were centrifuged at 500 *× g* at 4*^◦^*C for 5 minutes. Media was removed carefully and the cells were washed once in pre-warmed PBS. 1 ml of cold 1*×*PBS was added to the cells and the pellet was gently broken down by tapping to create a homogeneous cell suspension. Cells were transferred to ice immediately. Four ml of ice-cold methanol was then added to the cells dropwise keeping the tubes on a vortex. Methanol-fixed cells were incubated on ice for 20 minutes and then aliquoted in 1.5 ml Eppendorf tubes and stored at *−*80*^◦^*C freezer.

#### Metabolic RNA labeling and scSLAM-seq

All buffers used for cell preparation were prepared from 20*×* SSC buffer (Invitrogen, #AM9763), UltraPure BSA (Invitrogen, #AM2616), and nuclease-free water (Invitrogen, #AM9932). Unless otherwise stated, buffers and cells were kept on ice at all times. Cells treated with 4-thiouridine were protected from light until the end of the alkylation step.

Methanol-fixed cells were removed from the *−*80*^◦^*C freezer and incubated on ice for 15 min to equilibrate at 4*^◦^*C before centrifugation at 1000 *× g* for 5 min. Methanol was carefully removed, and cell pellets were resuspended in 100 *µ*L of 3*×* SSC + 0.04% BSA buffer. For alkylation, 100 *µ*L of 20 mM iodoacetamide (Thermo Scientific, #A39271) in DMSO were added to the cells, followed by 30 min of incubation on ice for rehydration and 30 min of incubation at 37*^◦^*C. The alkylation reaction was quenched by adding 20 *µ*L of 1 M DTT and incubating cells on ice for 5 min before adding 500 *µ*L of 3*×* SSC + 0.04% BSA buffer. Cells were washed to remove BSA by centrifugation at 1000 *× g* for 5 min and resuspension in 500 *µ*L of 3*×* SSC buffer, followed by a second centrifugation and resuspension in 180 *µ*L of 3*×* SSC buffer. MULTI-seq anchor LMO-barcode mixes specific to each cell sample were prepared by combining 1.6 *µ*L of 50 *µ*M LMO (Sigma, #LMO001A), 8 *µ*L of 10 *µ*M barcode DNA oligo (IDT), and 10.4 *µ*L of 1*×* DPBS without Ca^2+^*/*Mg^2+^ (Gibco, #14190144). MULTI-seq co-anchor solution was prepared by combining 1.6 *µ*L of 50 *µ*M co-anchor (Sigma, LMO001B) and 18.4 *µ*L of 1*×* DPBS for each cell sample. Cells were labeled by adding 20 *µ*L of anchor LMO-barcode mix, followed by gentle mixing and incubation on ice for 5 min. The labeling reaction was continued by adding 20 *µ*L of co-anchor solution, followed by gentle mixing and incubation on ice for 5 min. The labeling reaction was finally quenched by adding 500 *µ*L of 3*×* SSC + 1% BSA, followed by centrifugation at 1000 *× g* for 5 min. The washing step with 3*×* SSC + 1% BSA was repeated for a total of two washes. Cell samples were finally resuspended in 100 *µ*L of 0.125*×* SSC + 0.04% BSA buffer. Cell samples were filtered through 70 *µ*m cell strainers (pluriSelect, #43-10070-50), enumerated with an automatic cell counter (Countess II), and pooled to achieve the desired representation.

The cell pools were loaded onto a Chromium X Controller (10x Genomics, #1000331) in separate lanes, aiming to recover 6000 cells per lane. Samples were processed using the Chromium Next GEM Single Cell 3’ v3.1 kit (10x Genomics, #1000268) to prepare the gene expression libraries according to the manufacturer’s protocol, with the following modifications: (1) in Step 2.2.a, 1 *µ*L of 2.5 *µ*M MULTI-seq primer was added to the cDNA amplification master mix; and (2) in Step 2.3.d, the supernatant was saved, as it contains the MULTI-seq barcodes.

MULTI-seq libraries were prepared according to the protocol of the original authors. A total of 80 *µ*L of saved supernatant containing MULTI-seq barcodes was mixed with 260 *µ*L of SPRIselect beads (Beckman, #B23318) and 180 *µ*L of isopropanol (Sigma, #33539). After 5 min of incubation, the bead pellets were washed twice with 500 *µ*L of 80% EtOH. After air-drying, bead pellets were resuspended in 50 *µ*L of EB buffer (Qiagen, #1014609). Next, libraries were constructed by PCR: 3.5 ng of eluted cDNA barcodes were mixed with 2.5 *µ*L of Universal I5 primer (10 *µ*M), 2.5 *µ*L of RPI primer (10 *µ*M), 26.25 *µ*L of Kapa HiFi HotStart ReadyMix (Roche, #KK2601), and nuclease-free water up to 50 *µ*L. PCR reactions were run in a thermocycler with the following program: (1) 95*^◦^*C for 5 min; (2) 8–12 cycles of 98*^◦^*C for 15 s, 60*^◦^*C for 30 s, and 72*^◦^*C for 30 s; and (3) 72*^◦^*C for 1 min. PCR products were cleaned with 80 *µ*L of SPRI beads, two washes with 200 *µ*L of 80% EtOH, and a final resuspension in 25 *µ*L of EB buffer.

General quality control of all libraries was performed using a Qubit 3.0 Fluorometer with the dsDNA High Sensitivity assay (Thermo Fisher Scientific, #Q33231) and a 2100 Bioanalyzer with the High Sensitivity DNA kit (Agilent, #5067-4626). Libraries were pooled aiming at sequencing depths of 160,000 reads per cell for gene expression libraries and 5,000 reads per cell for MULTI-seq libraries. Gene expression libraries were sequenced on a DNBSEQ-G400 platform (BGI) in paired-end mode (Read 1: 28 bp; Read 2: 90 bp), whereas MULTI-seq libraries were sequenced on an Illumina NovaSeq 6000 platform using an S2 flow cell (Read 1: 28 cycles; Read 2: 90 cycles; Index 1: 10 cycles; Index 2: 10 cycles).

### Data processing and datasets

We developed buRst, a likelihood-based framework for inferring transcriptional bursting kinetics from single-cell SLAM-seq data. The method was evaluated by simulation and then applied to scSLAM-seq measurements of acute MYC depletion.

#### Processing and analysis of the sci-fate dataset

For simulation-based validation, we used the previously published sci-fate dataset to obtain realistic gene-specific kinetic parameters [13]. FASTQ files from the dexamethasone-treatment dataset were downloaded from the Sequence Read Archive under accession SRX5848248 and processed as described in Berg *et al.* [41]. Read-count matrices, new-to-total RNA ratio matrices, and discrete labeling likelihoods were generated using GRAND3 version 3.0.9 with the options -D, -progress, -profile, -estim Full, and -outputDiscreteLoglik [41]. Cells from the 0 h and 2 h time points were retained for subsequent analyses.

Gene-specific kinetic parameters were estimated separately at 0 h and 2 h. The 0 h samples were analyzed under the steady-state model, whereas the 2 h samples were analyzed using the non-steady-state model with the 0 h total RNA distribution as the initial reference distribution. Genes with confident parameter estimates were retained.

Pseudobulk abundances were estimated separately using the same GRAND3 workflow. Cells were aggregated according to time point using the options -pseudobulkName and -pseudobulkFile, with a pseudobulk definition file based on the sample annotations used by Berg *et al.* [41]. Total and newly synthesized RNA abundances were therefore re-estimated directly from the aggregated conversion-count data rather than obtained by summing cell-level estimates. Pseudobulk log_2_(2 h*/*0 h) fold changes were calculated separately for total and newly synthesized RNA.

#### Processing of MYC scSLAM-seq data

Raw reads were processed with cellranger count (Cell Ranger Martian runtime v4.0.8; STAR v2.7.2a) against a custom human Ensembl 90 reference containing the Tir1 transgene. Sample identities were assigned from MULTI-seq barcode reads using deMULTIplex. Following demultiplexing and negative-cell rescue, 7,044 cells were assigned to the six retained sample barcodes.

The Cell Ranger BAM file was converted to CIT format using Gedi Bam2CIT (GEDI v1.0.6f). GRAND3 v3.0.9 [41] generated count matrices, NTR estimates and discrete labeling likelihoods using -D, -progress, -profile, -estim Full, and -outputDiscreteLoglik. Outputs were imported into grandR [31]. Processing settings, sequencing metrics, demultiplexing and subsequent cell filtering are detailed in Supplementary Section 7.1.

#### MYC ChIP-seq processing

MYC ChIP-seq and matched input-control data from A375 cells were obtained from the dataset reported by Vogt *et al.* [32] and reprocessed from FASTQ files. Reads were aligned to the GRCh38.p10 human genome assembly (GCA_000001405.25). Mouse spike-in reads were used to normalize the ChIP-seq and input-control signal. MYC peaks were called with MACS2 using the MYC DMSO ChIP sample as treatment and the spike-in-normalized, downsampled input signal as control. Peak calling was performed with the human genome size option (-g hs) and a *p*-value threshold of 0.01. This identified 112,952 MYC peaks. Peaks were assigned to nearby transcription start sites (TSSs), and promoter-associated peaks were retained if the distance between the peak summit and assigned TSS was at most 3 kb, resulting in 30,274 promoter-associated MYC peaks. Promoter-level MYC annotations were then integrated with new RNA fold changes and gene-specific changes in bursting kinetics inferred from the scSLAM-seq data.

### Statistical modeling and kinetic inference

#### Read-level labeling evidence

Read-level T-to-C conversion evidence was evaluated using the GRAND-SLAM measurement model [10]. For each gene–cell observation with total count *N_i_*, we calculated a discrete log-likelihood profile over all possible new RNA counts *k* = 0*, …, N_i_*. These profiles retained labeling uncertainty for subsequent kinetic inference rather than replacing the latent new RNA count with a point estimate. Profile construction and storage are described in Supplementary Section 1.

#### Transcriptional bursting model

Expression of RNA molecules (*M*) was modeled as an effective instantaneous bursting process. Burst events occurred at rate *k_m_*. At each event, a random burst size *B* was drawn, and the RNA count increased by its realized value *b*:

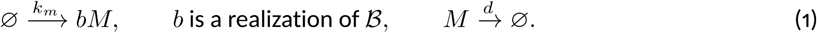

Individual RNA molecules degraded independently at rate *d*, which is related to RNA half-life by

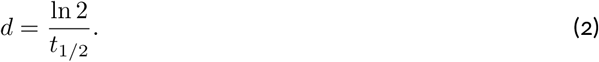

The burst-size random variable *B* followed a geometric distribution on *{*0, 1, 2*, …}*:

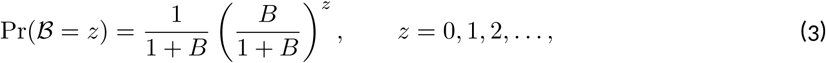

where *B* = E[*B*] is the mean burst size. The mean RNA production rate was therefore

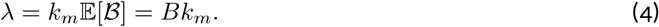

Our treatment of time-dependent RNA count distributions builds on the transient bursting framework of Singh & Bokes [6], which we adapted to the metabolic-labeling setting by distinguishing newly synthesized RNA from surviving pre-existing RNA. This effective compound-Poisson burst model does not include an explicit promoter ON state, OFF state, or active-state duration. It can be interpreted as the short-burst limit of more detailed promoter-switching models. The full probability-generating-function derivation, moment equations, and numerical evaluation of the model distributions are given in Supplementary Sections 2 and 3.

#### New, old, and total RNA count distributions

For labeling duration *t*, the observed total RNA count is

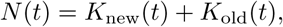

where *K*_new_ counts molecules synthesized after labeling onset and *K*_old_ counts surviving pre-existing molecules. Given ***θ*** = (*B, k_m_, d*) and the initial RNA distribution, these populations are independent, and their PMFs give

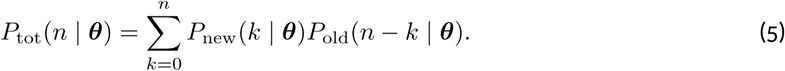

The component distributions were evaluated by FFT inversion and combined by discrete convolution.

For steady-state inference, the initial old RNA population followed the stationary bursting distribution. For non-steady-state inference, it was represented by the empirical total RNA distribution of a reference condition, treated as fixed. Pre-existing RNA was propagated through first-order degradation, while new RNA was generated over the labeling interval. Numerical evaluation and boundary conditions are detailed in Supplementary Sections 3 and 4.5.

#### Joint likelihood for kinetic inference

For a gene–cell observation with total count *N_i_*, the bursting model implies the conditional probability that *k* of the molecules are newly synthesized:

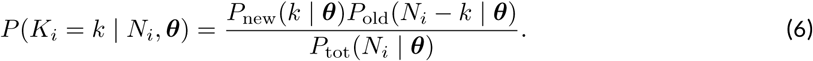

The centered GRAND-SLAM profile provides a relative measurement likelihood

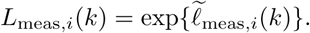

The likelihood contribution of cell *i* was obtained by marginalizing over the latent new RNA count:

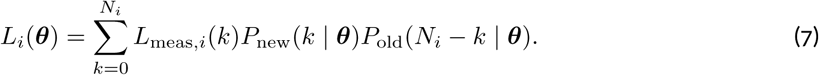

The full gene-level log-likelihood was

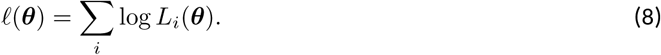

The marginalization was evaluated in log space using the log-sum-exp identity. Cells with zero total counts contributed through *P*_new_(0)*P*_old_(0) and required no additional labeling-evidence term. Component PMFs were normalized over their full, adaptively selected numerical support before probabilities corresponding to observed counts were extracted. Numerical implementation details are provided in Supplementary Section 4.2.

For validation analyses using point estimates of new and old RNA counts, the measurement likelihood was omitted and the component-count likelihood was evaluated directly from *P*_new_ and *P*_old_. For the beta approximation, per-cell beta distributions over the new-to-total RNA ratio were discretized over possible new RNA counts and substituted for the full measurement likelihood in the same marginalization. Details of the profile-based, point-estimate, and beta-approximation likelihoods are given in Supplementary Sections 4 and 6.

#### Parameter estimation and uncertainty

Kinetic parameters were inferred separately for each gene and condition. To improve numerical behavior, optimization was performed in the transformed coordinate system

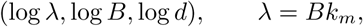

and the burst-arrival frequency was recovered as *k_m_* = *λ/B*. Initial values were obtained from the aggregate new RNA fraction and the first two moments of the estimated new RNA count distribution. For non-steady-state initialization, old RNA abundance in the fitted population and total RNA abundance in the reference population were compared on a per-cell mean scale, accounting for unequal numbers of cells. The log-likelihood was maximized in these coordinates using the Nelder–Mead algorithm with relative tolerance 10*^−^*^6^ and a maximum of 1,000 iterations.

Fits were attempted only for genes with sufficient information in the fitted condition. The default filters required at least five fitted cells, a total count sum of at least 50, more than three distinct total-count values, at least 5% of cells with non-zero total counts, an estimated new-count sum of at least 10, more than three distinct estimated new-count values, and at least 3% of cells with non-zero estimated new counts. For non-steady-state fits, at least five cells were also required in the reference condition.

Uncertainty was summarized by a local Laplace approximation, using the inverse observed information matrix at the maximum-likelihood estimate. A total of 20,000 Gaussian draws in (log *λ,* log *B,* log *d*) coordinates were transformed to *λ*, *B*, *d* and *k_m_*. Maximum-likelihood estimates were reported alongside the median and central 95% interval of the transformed Laplace draws.

Local identifiability was assessed using the eigenvalue ratio and condition number of the observed information matrix, together with parameter correlations in both (log *λ,* log *B,* log *d*) and (log *B,* log *k_m_,* log *d*) coordinates. Fits with pronounced likelihood ridges or strong parameter trade-offs were flagged and retained with their diagnostic information. Numerical initialization, support selection, Hessian regularization, and fit-status definitions are described in Supplementary Sections 3 and 4.

#### Comparison of kinetic parameters between conditions

Condition-specific uncertainty was propagated by independently sampling parameter vectors from the two fitted Laplace approximations while preserving parameter covariance within each condition. For *q ∈ {λ, B, k_m_, d}*, draw-wise log_2_ fold changes were calculated as log_2_(*q_c_*_2_ */q_c_*_1_) and summarized by their median and central 95% interval.

Burst-frequency and burst-size changes were assessed jointly. A kinetic change was considered supported when the squared Mahalanobis distance from the no-change point exceeded 5.99 and at least 90% of joint draws lay outside a no-change circle of radius 0.3. Supported changes were classified as frequency dominated, size dominated, or combined using posterior region probabilities, with opposing responses evaluated separately. Changes lacking sufficient support for mechanistic attribution were designated unresolved. Full classification rules and numerical details are provided in Supplementary Section 4.9.

Fold changes were calculated as 2 h relative to 0 h for sci-fate and auxin relative to DMSO for MYC depletion. MYC-depletion figures display the reverse contrast, log_2_(DMSO*/*Aux), so positive values indicate a reduction following depletion. For visualization, concordant and opposing joint responses were pooled into a single combined category.

### Simulation and benchmarking

#### Simulation of RNA counts and labeling evidence

RNA count trajectories were simulated using an exact Gillespie algorithm implemented in C++ through Rcpp. Burst events occurred with propensity *k_m_*, and degradation occurred with total propensity *d*(*n*_new_ + *n*_old_). At a degradation event, the removed molecule was assigned to the new or old population in proportion to the current population sizes.

The validation simulation was initialized at steady state. For each gene, initial old RNA counts were sampled from the stationary probability mass function calculated by FFT, while the new RNA population was initialized at zero. Independent trajectories were generated for each gene and cell over a 2 h labeling interval.

Gene-specific generating parameters were obtained from buRst fits to the 0 h condition of the sci-fate dataset. Genes whose fits were classified as bad_hessian or strong_tradeoff were excluded. The remaining 8,254 genes were simulated using their estimated burst sizes, burst frequencies, and degradation rates. For each gene, 1,151 cells were simulated using random seed 1234.

Simulated new- and old RNA counts were assembled into sparse gene-by-cell matrices and converted into grandR objects. GRAND-SLAM-like read-level evidence was generated by simulating uridine content and T-to-C conversion counts for old and newly synthesized molecules. The full simulation procedure, read-evidence model, output formats, and reproducibility settings are described in Supplementary Section 5.

#### Simulation of non-steady-state kinetic changes

To validate inference of kinetic changes under non-steady-state conditions, we generated an artificial perturbation benchmark using kinetic parameters fitted to the sci-fate A549 0 h condition. Fits flagged as bad_hessian or strong_tradeoff were excluded.

Each retained gene was assigned one of 25 predefined perturbations: no change; frequency-only or size-only changes with log_2_ fold changes of *±*0.5, *±*1, or *±*2; or joint changes of equal absolute magnitude in the same or opposing directions. Perturbation combinations were distributed approximately evenly across genes and randomly assigned. Degradation rates were unchanged.

For each gene, 1,151 cells were simulated in each condition. The reference population was generated under steady-state kinetics. The perturbed population was initialized directly from the simulated reference total RNA counts and propagated for 2 h under the perturbed parameters. New and pre-existing RNA counts and read-level labeling evidence were generated for both conditions.

The reference and perturbed conditions were analyzed using the steady-state and non-steady-state inference models, respectively, with the simulated reference population providing the initial distribution for the latter. Estimated fold changes, interval coverage, and mechanistic classifications were evaluated against the generating values. Simulation settings and perturbation assignment are detailed in Supplementary Section 5.5.

#### Comparison of full and approximate labeling likelihoods

Simulated data were analyzed using three representations of labeling evidence: the full discrete likelihood, an integer new-count point estimate, and a beta approximation to per-cell NTR uncertainty discretized over possible new RNA counts. The kinetic model, initialization, optimization settings, and uncertainty approximation were otherwise unchanged.

Recovery of *q ∈ {λ, d, k_m_, B}* was quantified by log (*q*_10_*/q*_true_). Empirical coverage was the fraction of genes whose 95% Laplace-approximation interval contained the generating value. Likelihood definitions and evaluation procedures are provided in Supplementary Section 6.

#### Benchmark against *burstMCMC*

*buRst* was benchmarked against *burstMCMC* [8] using the same simulated molecule counts and read-level conversion evidence. The simulated evidence was exported in a *burstMCMC*-compatible format containing, for each gene and cell, the total UMI count together with the number of T-to-C conversions and informative uridine positions for each simulated molecule.

The background conversion probability and the labeled RNA conversion probability were fixed to the values used during simulation, *p*_old_ = 10*^−^*^4^ and *p*_new_ = 0.04. The labeling duration was supplied as 120 min. Because the simulation did not include cell-specific capture efficiency, the burstMCMC capture-efficiency parameter was fixed to 0.01 for all cells. Fitting was initiated independently for each gene using burstMCMC model 2, with a restart under model 3 following repeated numerical likelihood failures. Successful fits from both models were included. Parameter recovery, empirical interval coverage, and per-gene runtime were compared on a shared set of genes passing the method-specific quality filters. Sampling settings, parameter-scale conversions, filtering criteria, and timeout handling are detailed in Supplementary Section 6.5.

### Analysis of MYC-dependent transcriptional regulation

#### MYC-depletion scSLAM-seq analysis

The processed grandR object was converted to Seurat with separate total and new RNA assays. Genes detected in at least 50 cells were retained. Independent PCA and UMAP embeddings were calculated for the two assays and recalculated after removal of low-quality populations and cells with inconsistent cell-line annotations (Supplementary Section 7.1). The Muhar 100-gene MYC signature [26] was scored on the new RNA assay using Seurat::AddModuleScore, case-insensitive gene matching and seed 1. Cell-cycle scoring used the Seurat 2019 S- and G2/M-phase gene sets when at least ten genes from each set were present.

Filtered cells were aggregated by cell line, treatment and 4sU-labeling status. Conversion-count data were reanalyzed with GRAND3 v3.0.9 using -D, -progress, -profile, -estim Full, -pseudobulkFile, and -pseudobulkName. Auxin-versus-DMSO fold changes were calculated separately for total and new RNA in each cell line. Phase-stratified pseudobulks additionally grouped cells by G1, S or G2/M phase. Pseudobulk fold changes were descriptive because the groups lacked independent biological replicates. Normalization is detailed in Supplementary Section 7.3.

Steady-state half-lives were estimated from the 2 h labeling experiment using grandR::ComputeSteadyStateHalfLives. For A375 DMSO, genes with finite half-life and total and new RNA fold changes were grouped by half-life to compare the two RNA responses (Supplementary Section 7.5). A375 and MaMel63 new RNA responses were also compared with the gene-level responses reported by Muhar *et al.* [26], using matched genes, harmonized contrasts, scatterplots and correlation analysis.Pairwise Pearson correlations of new RNA responses were calculated before and after excluding the Muhar top-100 signature genes, using the eligible genes available for each cell-line pair. Confidence intervals were calculated using Fisher’s *z* transformation.

A shared MYC-response signature was derived from new RNA log_2_(MYC present*/*depleted) fold changes in A375, MaMel63, K562 and HCT116 cells using a two-component, four-dimensional multivariate *t* mixture with component-specific scale matrices and degrees of freedom (teigen, model UUUU). Genes were ranked by posterior membership in the component with higher mean responses in all four cell lines. The largest ranked set with an average posterior background probability of at most 0.01 defined the BFDR01 signature. Cross-cell-line background correlations were derived from the covariance matrix of the near-zero component (Supplementary Section 7.4).

Cell-cycle comparisons were restricted to BFDR01 genes with finite new RNA fold changes in all three phases within each cell line. Pairwise comparisons used two-sided paired Wilcoxon signed-rank tests across genes, with Benjamini–Hochberg correction across six comparisons. Effect sizes were median paired differences in log_2_ fold change.

Kinetic inference was restricted to 4sU-labeled cells; no-4sU controls were used for labeling quality control. Bursting kinetics were inferred separately for each cell line, using steady-state fits for DMSO and non-steady-state fits for auxin. Both labeling and modeled propagation lasted 2 h. The DMSO total RNA distribution approximated the population at labeling onset; changes during the preceding 2 h auxin pretreatment were not explicitly modeled. Downstream comparisons required successful fits in both conditions and excluded bad_hessian and strong_tradeoff fits. Fitting and figure-subset selection are detailed in Supplementary Section 8.

#### MYC ChIP-seq annotation and integration with transcriptional responses

Promoter-associated MYC ChIP-seq peaks were annotated with quantitative MYC binding, E-box motif content, CpG-island overlap, and TATA-like promoter motifs. Input-control coverage was rescaled before comparison with the MYC ChIP-seq signal. For each promoter-associated peak, a background-subtracted MYC signal and a MYC-to-input coverage ratio were calculated. Peaks with zero input coverage were assigned missing quantitative ratios rather than infinite values.

For promoters associated with multiple MYC peaks, one representative peak was selected for downstream gene-level analyses. Because the scSLAM-seq and bursting results were estimated at gene level, one representative promoter annotation was then selected per gene while retaining the total number of distinct promoter-associated MYC peaks as a separate feature. Exact motif definitions, genomic windows, representative-peak selection rules, and promoter annotation procedures are provided in Supplementary Section 9.

The promoter-level MYC annotations were integrated with A375 pseudobulk new RNA fold changes and gene-specific changes in burst size and burst frequency. Ensembl version suffixes were removed before joining datasets. Quantitative MYC binding was represented as the log_10_ MYC-to-input coverage ratio of the representative promoter-associated peak. For interpretability, response signs were oriented such that positive values indicate a decrease after MYC depletion:

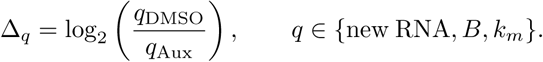

The association between quantitative MYC binding and new RNA response was summarized using Spearman’s rank correlation. Separate univariable ordinary least-squares regressions related quantitative binding to new RNA, burst-size, and burst-frequency responses. These three regressions used a common gene set with finite values for all three responses and a finite, positive MYC-to-input coverage ratio. Associations between new RNA responses and kinetic responses were additionally summarized using ordinary least-squares regression and Pearson’s correlation. Additional categorical analyses compared genes according to peak number and compared E-box-containing and non-E-box promoters within tertiles of quantitative MYC binding, using two-sided Wilcoxon rank-sum tests with Benjamini–Hochberg correction within each analysis. The ability of promoter-binding features to identify strongly MYC-dependent genes, defined by a new RNA log_2_(DMSO*/*Aux) fold change of at least one, was evaluated by receiver-operating-characteristic analysis, including a five-fold cross-validated logistic-regression predictor combining quantitative occupancy and E-box presence. Details are provided in Supplementary Section 10.

#### Association of promoter features with transcriptional and kinetic responses

Associations between promoter features and new RNA, burst-size, and burst-frequency responses were evaluated using separate multivariable linear models. Each model included quantitative promoter-associated MYC binding, E-box presence, TATA-box presence, CpG-island annotation, log-transformed absolute distance to the transcription start site, and MYC-peak position encoded as upstream or downstream. Quantitative MYC binding was represented by the log_10_ MYC-to-input coverage ratio, and E-box annotations were collapsed into a binary presence/absence variable. All three models used a common gene set with finite new RNA, burst-size, and burst-frequency responses and complete predictor information. The input bursting estimates had passed the condition-fit exclusions described above; no additional precision filtering was applied for the regression analyses. Predictor-specific *P* values were adjusted using the Benjamini–Hochberg procedure across the 18 non-intercept coefficients from the three models. Further details are provided in Supplementary Section 10.3.

Burst-size and burst-frequency responses were additionally compared between other genes and genes belonging to the Muhar or BFDR01 signatures using two-sided Wilcoxon rank-sum tests. These comparisons were restricted to the four-cell-line expression-analysis universe. Other genes excluded both signatures, while genes shared between the signatures were included in both signature groups. Benjamini–Hochberg correction was applied across the four comparisons. Further details are provided in Supplementary Section 10.4.

## Supporting information

Supplementary Note: Mathematical and computational details for buRst

## Data and code availability

The buRst R package is available at https://burst.erhard-lab.de, with source code hosted at https://github.com/erhard-lab/buRst.

Analysis workflows, simulation code, processed inputs, and saved results underlying the figures are publicly available on Zenodo at https://doi.org/10.5281/zenodo.24362854.

Sequencing data will be made publicly available through GEO upon publication of the peer-reviewed article.

## Use of AI-assisted tools

OpenAI’s ChatGPT (ChatGPT Plus subscription, accessed via the web interface at https://chatgpt.com), primarily with GPT-5-series models, was used to assist with text formulation, improving spelling, grammar, and sentence structure, and code development, debugging, and optimization during manuscript preparation. All AI-assisted text and code were reviewed by the authors, and code was validated before use in the analyses. The authors take full responsibility for the scientific content, analyses, interpretations, and conclusions presented in this manuscript.

## Acknowledgements

We gratefully acknowledge the support of all participating research groups and institutions.

---

**Figure S1:**
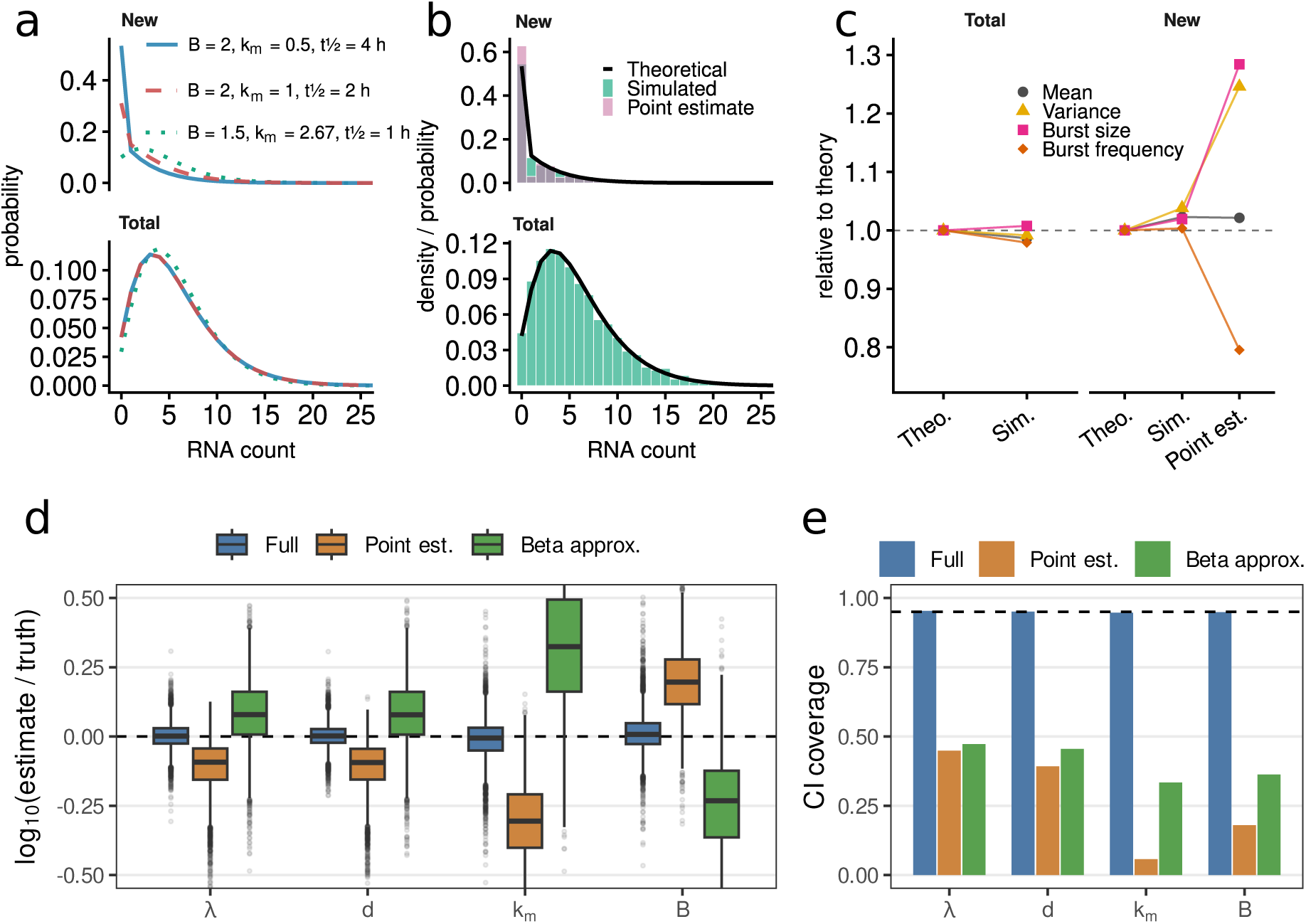
Simulation-based validation of bursting-parameter inference. (**a**) Representative theoretical RNA-count distributions for a reference model (blue), a turnover-alternative model (red), and a bursting-parameter-alternative model (green), shown separately for newly synthesized and total RNA. (**b**) Comparison of theoretical RNA-count distributions (black lines), simulated RNA counts (green), and point estimates of newly synthesized RNA counts (pink), shown separately for newly synthesized and total RNA. (**c**) Relative mean, variance, burst-size, and burst-frequency estimates for theoretical values, simulated data, and point estimates, normalized to the theoretical expectation. (**d**) Accuracy of parameter estimation across simulations, shown as the log_10_ ratio of estimated to true values for the full model, point-estimate approach, and beta approximation. (**e**) 95% approximate posterior credible interval coverage for RNA production rate (*λ*), degradation rate (*d*), burst frequency (*k_m_*), and burst size (*B*) across the evaluated inference approaches. The dashed horizontal line indicates nominal coverage.

**Figure S2:**
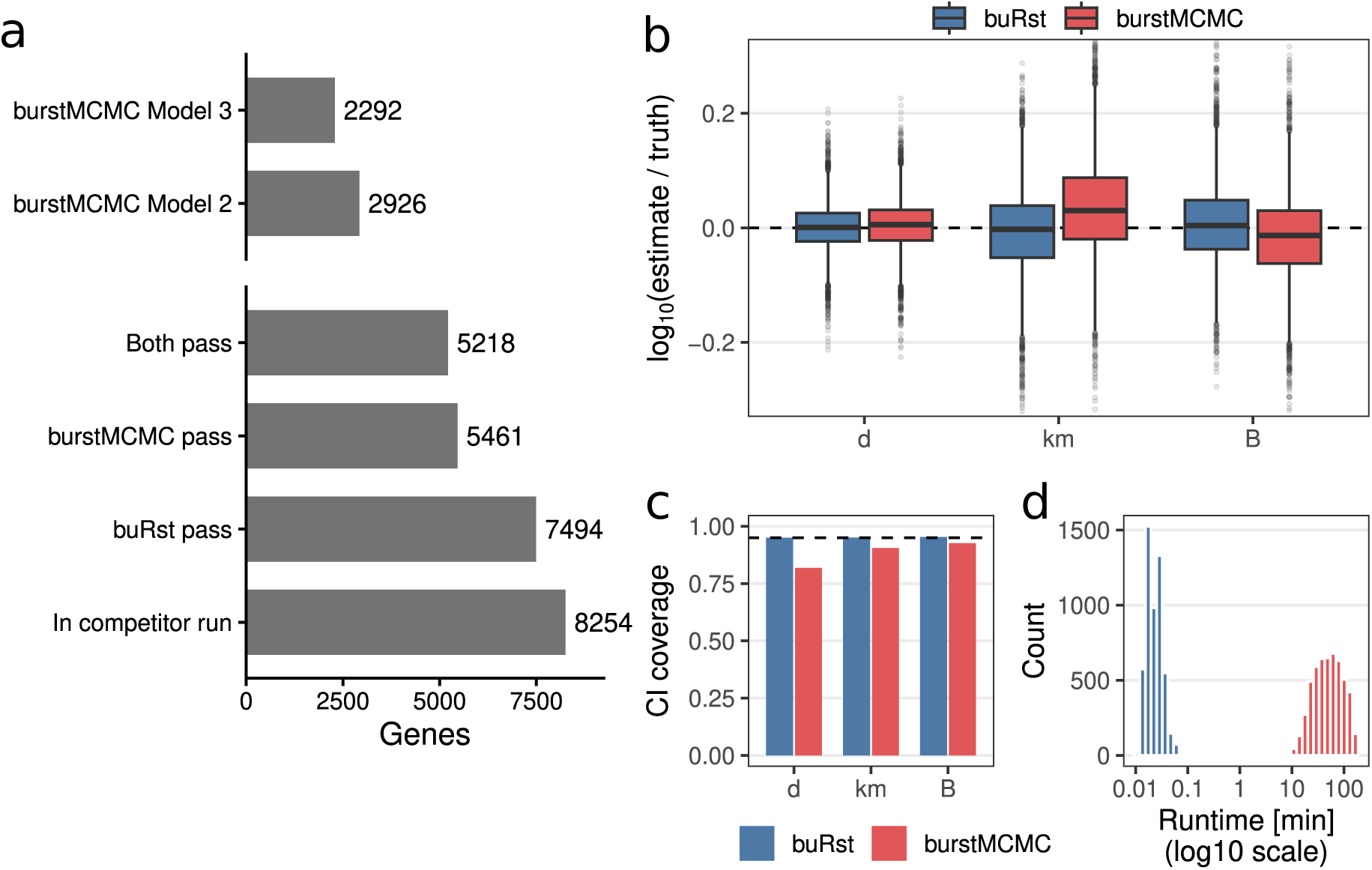
Benchmarking burst-kinetic inference against burstMCMC. (**a**) Number of genes retained at successive filtering and model-selection steps for the competitor run, the buRst analysis, burstMCMC, the intersection of genes passing both methods, and burstMCMC model classes. (**b**) Accuracy of inferred degradation rate (*d*), burst frequency (*k_m_*), and burst size (*B*), shown as the log_10_ ratio of estimated to true parameter values for buRst and burstMCMC. (**c**) 95% approximate posterior credible interval coverage for inferred kinetic parameters. The dashed horizontal line indicates nominal coverage. (**d**) Runtime comparison between buRst and burstMCMC, shown on a log_10_ scale.

**Figure S3:**
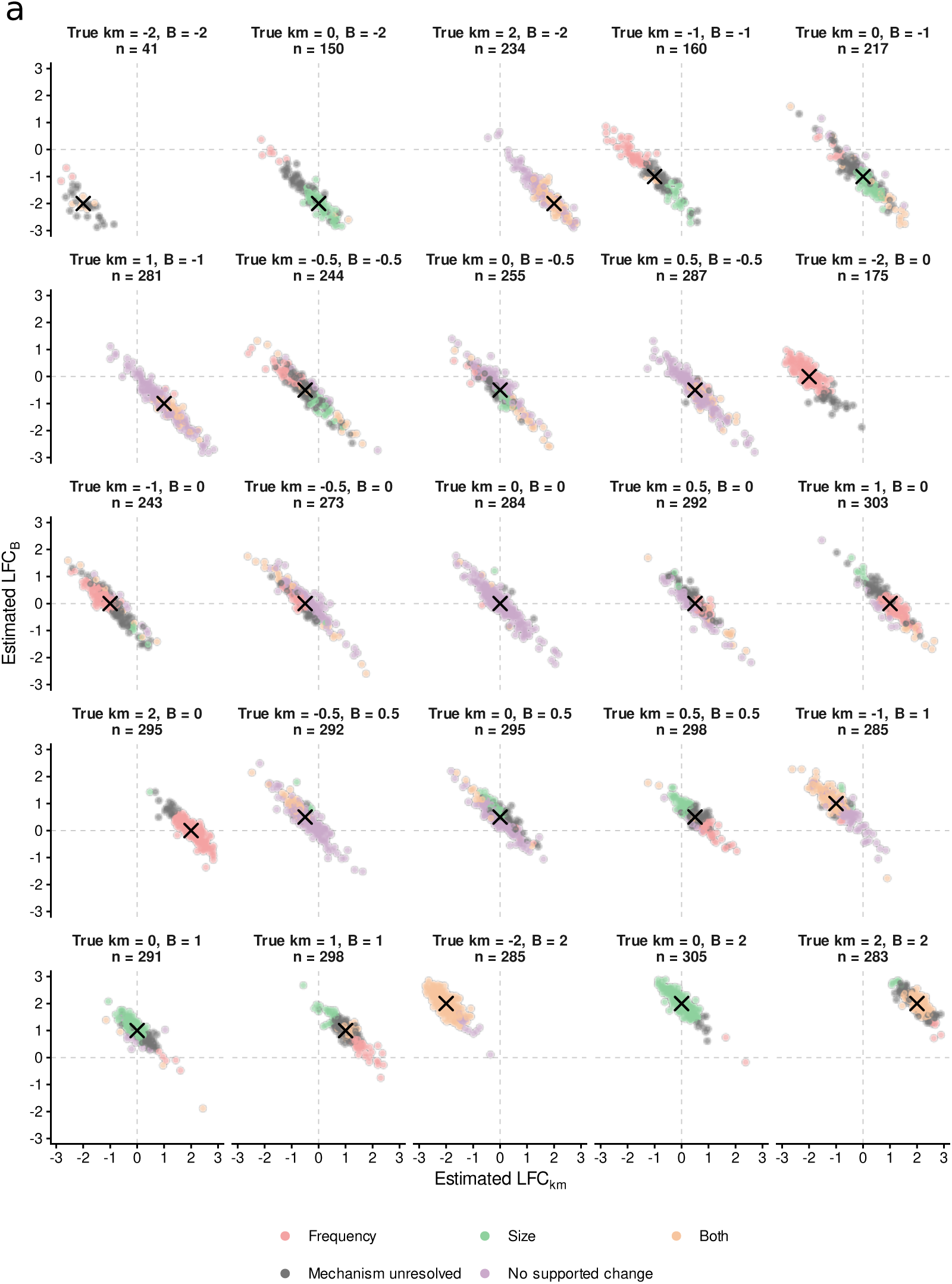
Joint recovery across simulated burst-frequency and burst-size perturbations. (**a**)Joint posterior median estimates of log_2_ fold changes in burst frequency (*k_m_*) and burst size (*B*) across the artificial perturbation grid. Each panel corresponds to one predefined combination of true burst-frequency and burst-size changes. Points represent individual genes after re-estimation with the complete buRst workflow and are colored according to their inferred kinetic response category. The true perturbation is indicated by a black *×*.

**Figure S4:**
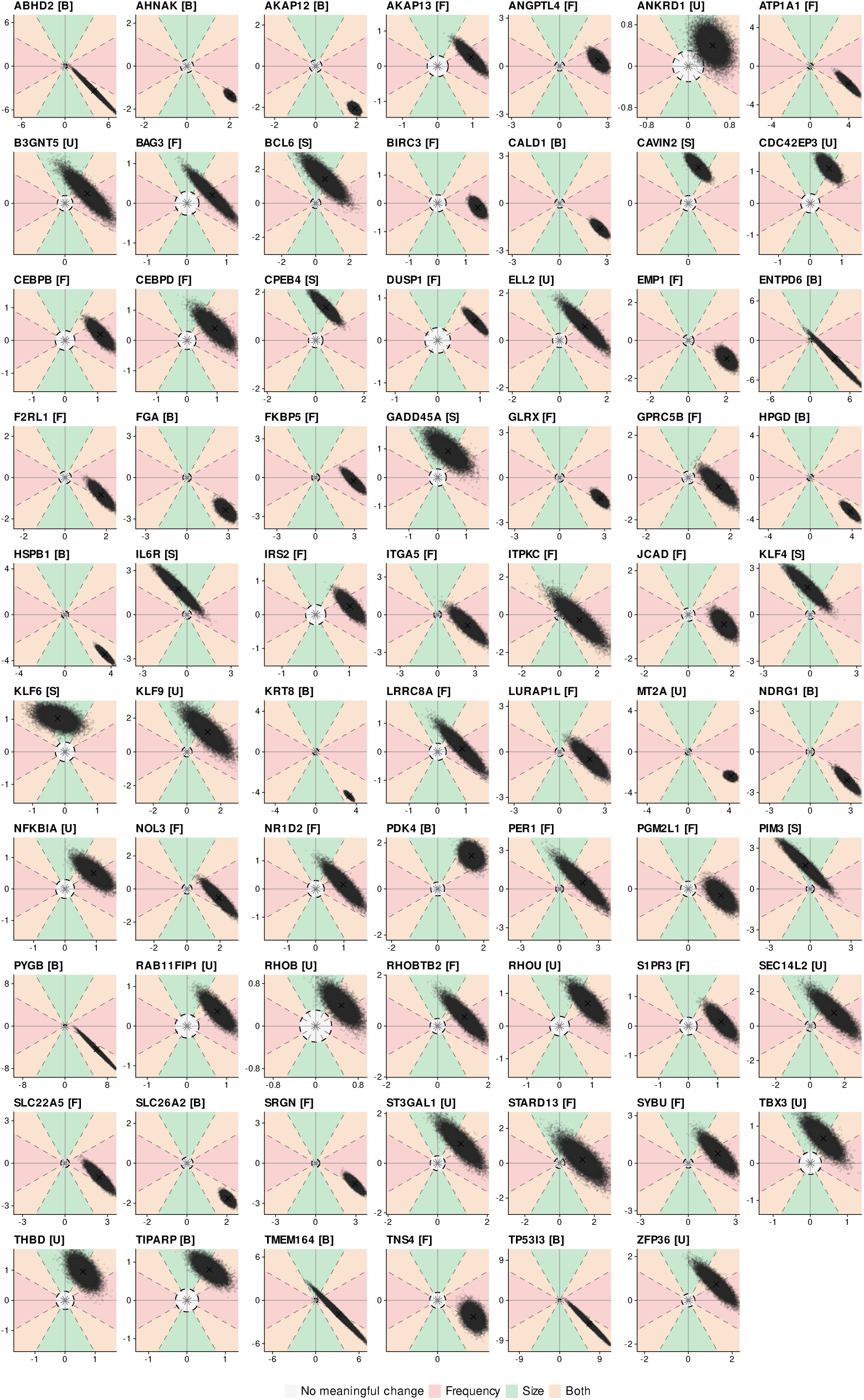
Gene-level posterior distributions of DEX-induced bursting changes. Joint posterior distributions of burst-frequency and burst-size log_2_(2h*/*0h) fold changes for the 69 genes with pseudobulk new RNA log_2_ fold change greater than 1 and evaluable bursting comparisons in the sci-fate dataset. Each panel represents one gene, with burst-frequency changes on the horizontal axis and burst-size changes on the vertical axis. Points show joint draws from the Laplace-approximated posteriors; contours indicate estimated posterior density, and crosses mark coordinate-wise posterior medians. Background colors indicate regions attributed to frequency changes (pink), size changes (green), or changes in both parameters (orange). The central grey region denotes no meaningful change (radius 0.3 in joint log_2_ fold-change space); dashed lines mark region boundaries. Letters in panel titles indicate the inferred classification: F, frequency; S, size; B, both; U, mechanism unresolved; N, no supported change. Axis ranges vary between genes.

**Figure S5:**
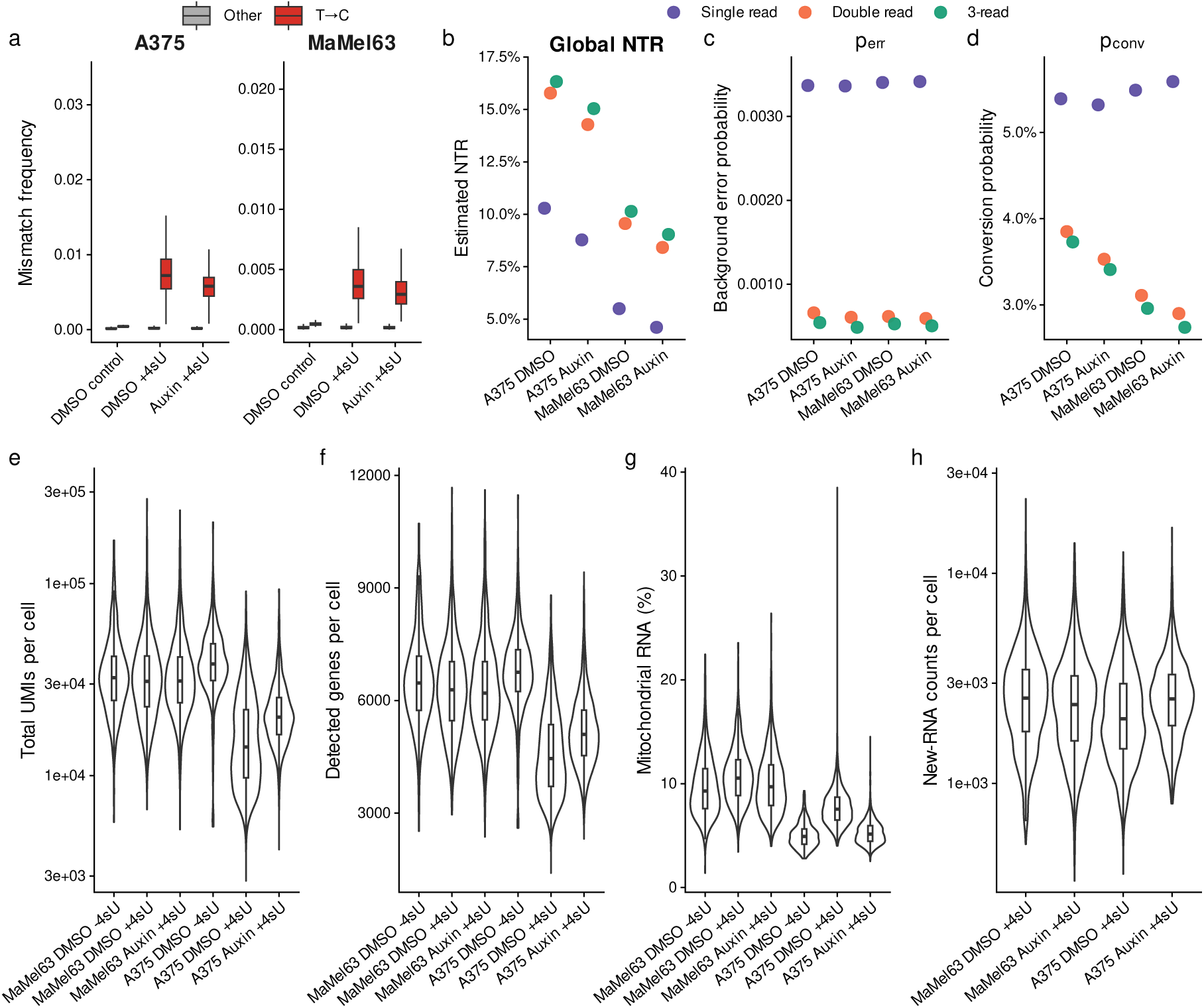
Quality control of 4sU labeling and single-cell RNA measurements. (**a**) Observed exonic mismatch frequencies in A375 and MaMel63 cells for unlabeled DMSO controls and 4sU-labeled DMSO- and auxin-treated cells, based on double-read mismatch calls to reduce sequencing errors. T*→*C mismatches are shown separately from all other mismatch types. (**b**–**d**) Global parameters estimated by GRAND3 for each cell line and treatment: global new-to-total RNA ratio (**b**), background-error probability *p*_err_ (**c**), and T*→*C conversion probability *p*_conv_ (**d**). Estimates are shown separately for UMIs supported by one, two, or at least three reads. (**e**–**g**) Distributions of total UMI counts per cell (**e**), detected genes per cell (**f**), and the percentage of counts assigned to mitochondrial genes (**g**) across cell lines, treatments, and labeling conditions. (**h**) Distribution of newly synthesized RNA counts per cell in the 4sU-labeled DMSO- and auxin-treated samples. Violin plots show the full cell-level distributions, with embedded boxplots indicating the median and interquartile range.

**Figure S6:**
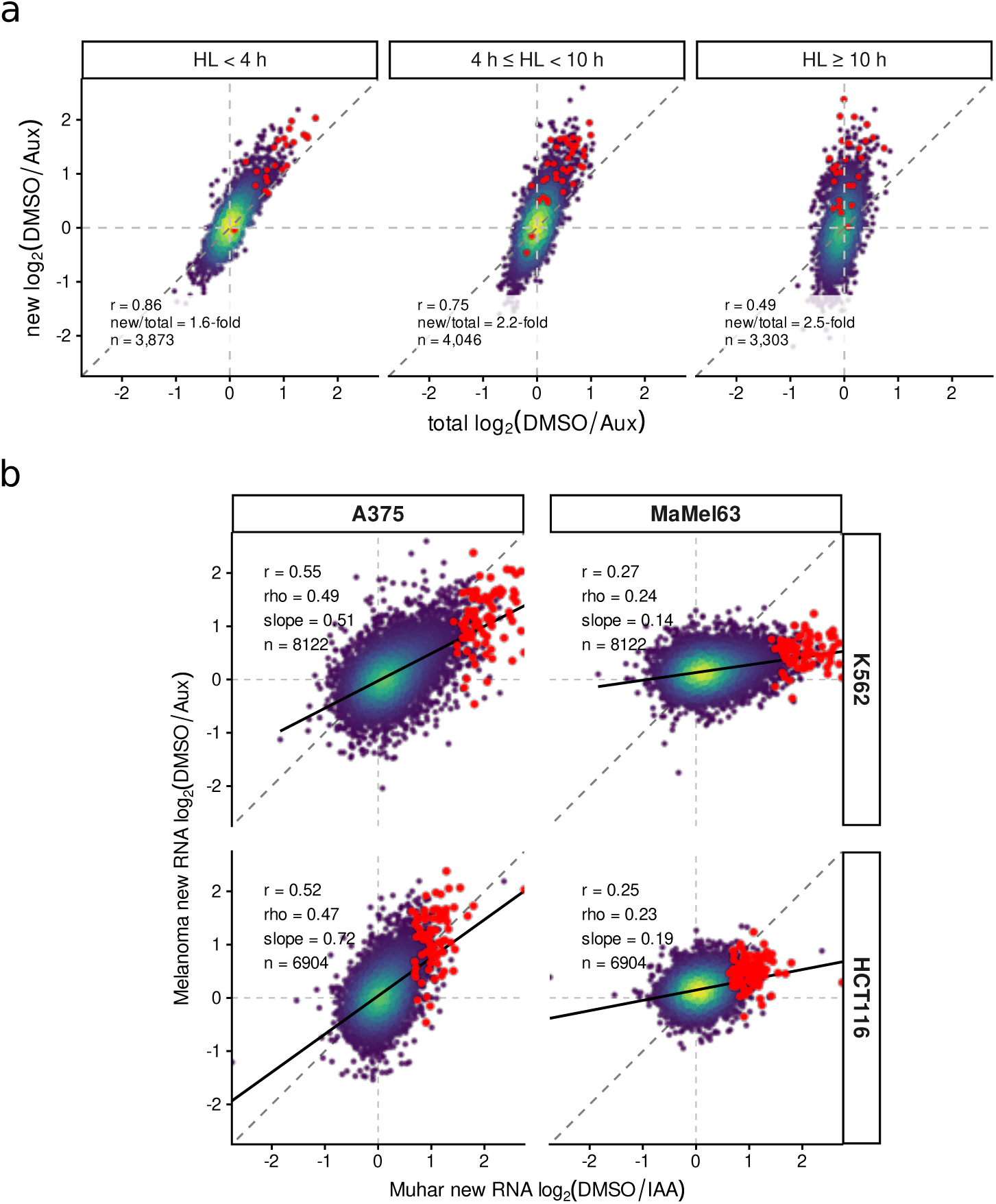
Validation and cross-dataset comparison of MYC-dependent transcriptional responses. (**a**) Relationship between total RNA and new RNA log_2_(DMSO*/*Aux) fold changes in A375 cells, stratified by transcript half-life (HL). Genes belonging to the MYC-response signature are highlighted in red. Dashed lines mark zero on each axis and the diagonal indicates equal total- and new RNA responses. Reported values indicate Pearson’s correlation coefficient (*r*), the ratio of the median absolute new RNA to total RNA fold-change magnitude (new/total), and the number of genes included in each half-life group (*n*). (**b**) Gene-wise comparison of new RNA log_2_ fold changes between the published K562 and HCT116 datasets from Muhar *et al.* [26] and the melanoma pseudobulk responses in A375 and MaMel63 cells. Genes belonging to the MYC-response signature are highlighted in red. Dashed lines mark zero on each axis and the diagonal indicates equal fold-change estimates. Reported values indicate Pearson’s correlation coefficient (*r*), Spearman’s rank correlation coefficient (*ρ*), the slope of the fitted linear regression, and the number of genes included in the comparison (*n*).

**Figure S7:**
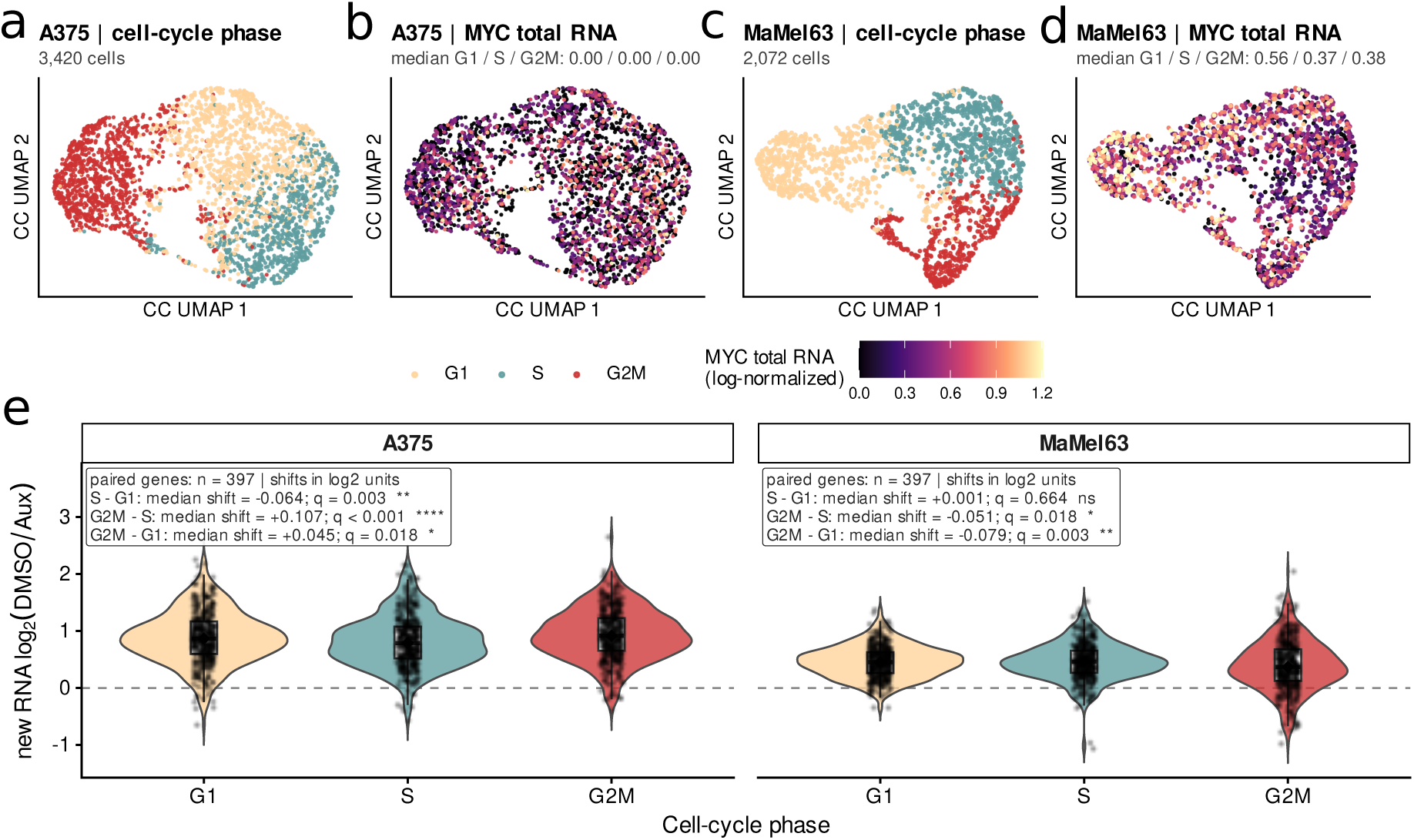
MYC expression and transcriptional activity across the cell cycle. (**a**,**c**) Cell-cycle UMAPs of 4sU-labeled A375 (*n* = 3,420 cells; **a**) and MaMel63 (*n* = 2,072 cells; **c**) colored by the assigned G1, S, or G2M phase. (**b**,**d**) The same UMAPs colored by log-normalized total MYC RNA abundance in A375 (**b**) and MaMel63 (**d**) cells. Median MYC RNA abundance in the G1, S, and G2M phases is reported above each panel. (**e**) New RNA log_2_(DMSO*/*Aux) fold changes of genes in the 438-gene BFDR01 signature, calculated separately for cells in the G1, S, and G2M phases in A375 (left) and MaMel63 (right). Positive values indicate reduced RNA synthesis following auxin-induced MYC depletion. Violin plots show the fold-change distributions across the 397 evaluable signature genes per cell line,quantified in all three phases within each cell line; embedded boxplots show the median and interquartile range. Points represent individual genes. Annotations report median paired differences in log_2_ units, with the subtraction direction indicated, and Benjamini–Hochberg-adjusted P values from paired Wilcoxon signed-rank tests across all six phase comparisons. Significance levels denote adjusted P values: ns, *P ≥* 0.05; *, *P <* 0.05; **, *P <* 0.01; ***, *P <* 0.001; ****, *P <* 0.0001.

**Figure S8:**
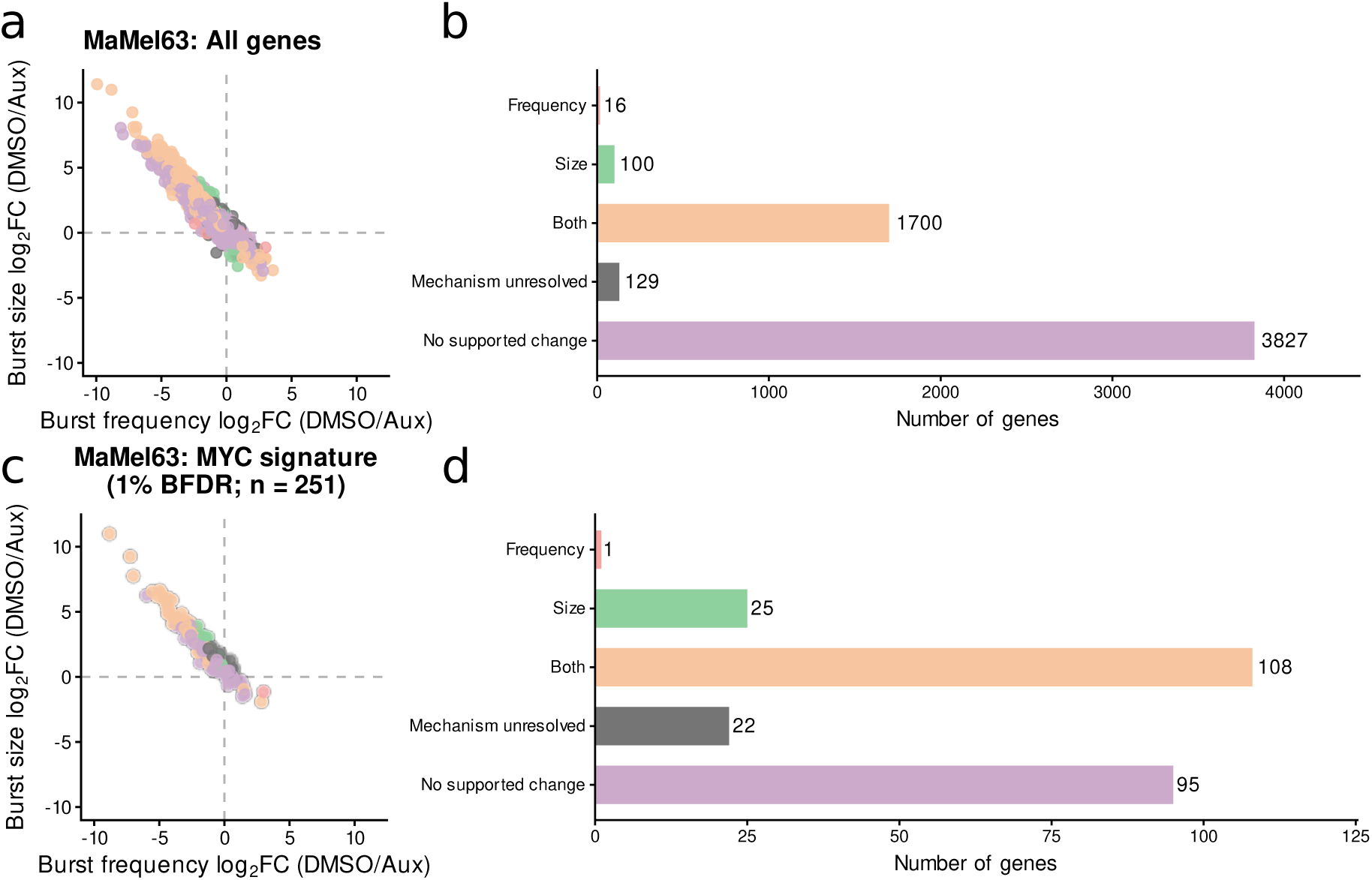
Classification of gene-level bursting responses after MYC depletion in MaMel63 cells. (**a**) Gene-wise log_2_(DMSO/Aux) fold changes in inferred burst frequency (*k_m_*) and burst size (*B*). Positive values indicate reduced burst frequency or burst size after auxin-induced MYC depletion. Point colours indicate whether the response was classified as predominantly involving burst frequency, burst size, or both parameters. (**b**) Number of genes assigned to each kinetic response category. (**c**) Burst-size and burst-frequency changes for the 251 genes from the shared BFDR01 MYC-response signature included in the bursting analysis. (**d**) Number of BFDR05 signature genes assigned to each kinetic response category.

**Figure S9:**
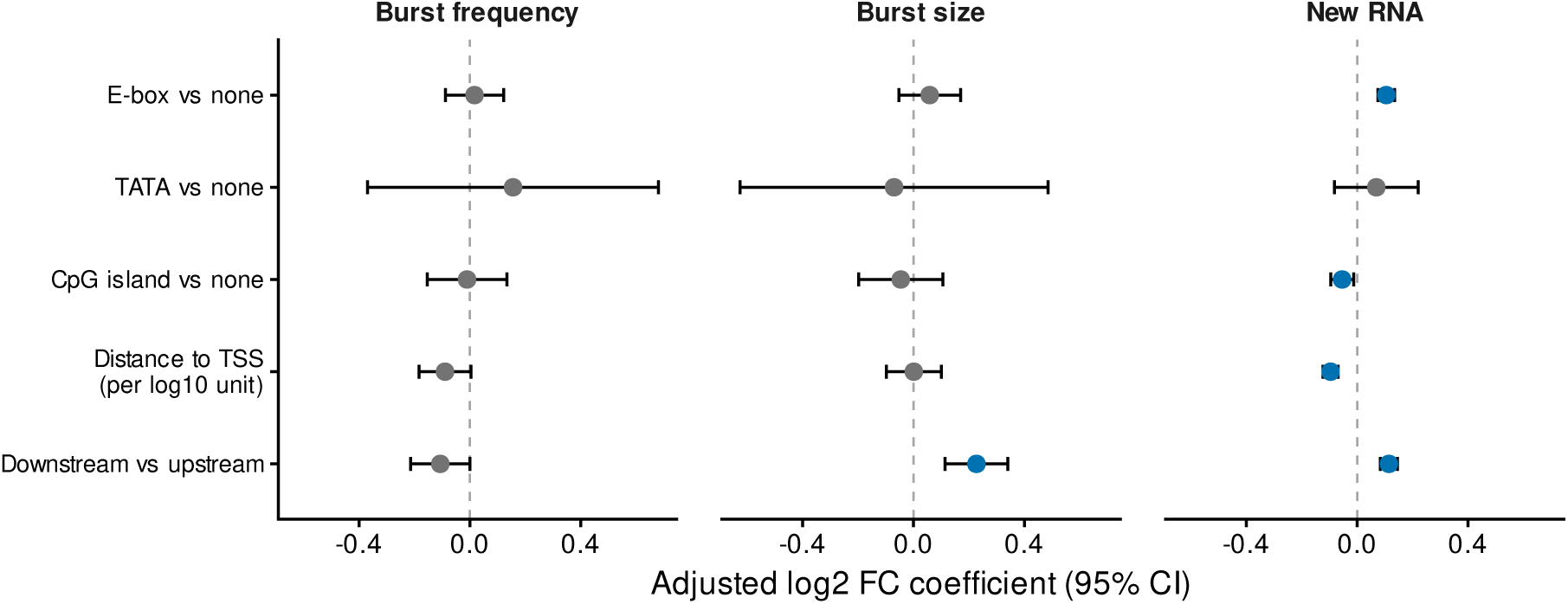
Adjusted associations between promoter features and transcriptional responses in A375 cells. Columns show associations with burst-frequency, burst-size, and new RNA log_2_(DMSO*/*Aux) fold changes. Separate multivariable linear models included quantitative MYC binding and all displayed promoter features. Absolute peak-to-TSS distance was transformed as log_10_(1 + *|*distance*|*), with distance expressed in base pairs. All models used the same eligible genes with complete response and predictor information, without additional condition-fit precision filtering. Points indicate regression coefficients, and bars show nominal 95% confidence intervals. Dashed vertical lines indicate no association. Blue points indicate Benjamini–Hochberg-adjusted *q <* 0.05, with correction across all 18 non-intercept coefficients from the three models; grey points indicate *q ≥* 0.05.

## References

1. Elowitz, M. B., Levine, A. J., Siggia, E. D. & Swain, P. S. Stochastic Gene Expression in a Single Cell. Science 297, 1183–1186 (Aug. 16, 2002).

2. Paulsson, J. Models of Stochastic Gene Expression. Physics of Life Reviews 2, 157–175 (June 2005).

3. Raj, A., Peskin, C. S., Tranchina, D., Vargas, D. Y. & Tyagi, S. Stochastic mRNA Synthesis in Mammalian Cells. PLoS biology 4, e309 ((Oct 2006).

4. Peccoud, J. & Ycart, B. Markovian Modeling of Gene-Product Synthesis. Theoretical Population Biology 48, 222–234 (Oct. 1995).

5. Suter, D. M. et al. Mammalian Genes Are Transcribed with Widely Different Bursting Kinetics. Science 332, 472–474 (Apr. 22, 2011).

6. Singh, A. & Bokes, P. Consequences of mRNA Transport on Stochastic Variability in Protein Levels. Bio-physical Journal 103, 1087–1096 (Sept. 5, 2012).

7. Larsson, A. J. M. et al. Genomic Encoding of Transcriptional Burst Kinetics. Nature 565, 251–254 (Jan. 2019).

8. Edwards, D. M., Davies, P. & Hebenstreit, D. Synergising Single-Cell Resolution and 4sU Labelling Boosts Inference of Transcriptional Bursting. Genome Biology 24, 138 (June 16, 2023).

9. Herzog, V. A. et al. Thiol-Linked Alkylation of RNA to Assess Expression Dynamics. Nature Methods 14, 1198–1204 (Dec. 1, 2017).

10. Jürges, C., Dölken, L. & Erhard, F. Dissecting Newly Transcribed and Old RNA Using GRAND-SLAM. Bioin-formatics 34, i218–i226 (July 1, 2018).

11. Erhard, F. et al. Time-Resolved Single-Cell RNA-Seq Using Metabolic RNA Labelling. Nature Reviews Methods Primers 2, 77 (Sept. 29, 2022).

12. Erhard, F. et al. scSLAM-Seq Reveals Core Features of Transcription Dynamics in Single Cells. Nature 571, 419–423 (July 18, 2019).

13. Cao, J., Zhou, W., Steemers, F., Trapnell, C. & Shendure, J. Sci-Fate Characterizes the Dynamics of Gene Expression in Single Cells. Nature Biotechnology 38, 980–988 (Aug. 2020).

14. Battich, N. et al. Sequencing Metabolically Labeled Transcripts in Single Cells Reveals mRNA Turnover Strategies. Science 367, 1151–1156 (Mar. 6, 2020).

15. Ramsköld, D. et al. Single-Cell New RNA Sequencing Reveals Principles of Transcription at the Resolution of Individual Bursts. Nature Cell Biology 26, 1725–1733 (Oct. 2024).

16. Meyer, N. & Penn, L. Z. Reflecting on 25 Years with MYC. Nature Reviews. Cancer 8, 976–990 (Dec. 2008).

17. Dang, C. V. MYC on the Path to Cancer. Cell 149, 22–35 (Mar. 2012).

18. Kress, T. R., Sabò, A. & Amati, B. MYC: Connecting Selective Transcriptional Control to Global RNA Production. Nature Reviews Cancer 15, 593–607 (Oct. 2015).

19. Wolf, E., Lin, C. Y., Eilers, M. & Levens, D. L. Taming of the Beast: Shaping Myc-Dependent Amplification. Trends in Cell Biology 25, 241–248 (Apr. 2015).

20. Rahl, P. B. et al. C-Myc Regulates Transcriptional Pause Release. Cell 141, 432–445 (Apr. 2010).

21. Lin, C. Y. et al. Transcriptional Amplification in Tumor Cells with Elevated C-Myc. Cell 151, 56–67 (Sept. 2012).

22. Nie, Z. et al. C-Myc Is a Universal Amplifier of Expressed Genes in Lymphocytes and Embryonic Stem Cells. Cell 151, 68–79 (Sept. 2012).

23. Baluapuri, A. et al. MYC Recruits SPT5 to RNA Polymerase II to Promote Processive Transcription Elongation. Molecular Cell 74, 674–687.e11 (May 16, 2019).

24. Sabò, A. et al. Selective Transcriptional Regulation by Myc in Cellular Growth Control and Lymphomagenesis. Nature 511, 488–492 (July 24, 2014).

25. Walz, S. et al. Activation and Repression by Oncogenic MYC Shape Tumour-Specific Gene Expression Profiles. Nature 511, 483–487 (July 24, 2014).

26. Muhar, M. et al. SLAM-Seq Defines Direct Gene-Regulatory Functions of the BRD4-MYC Axis. Science 360, 800–805 (May 18, 2018).

27. Patange, S. et al. MYC Amplifies Gene Expression through Global Changes in Transcription Factor Dynamics. Cell Reports 38, 110292 (Jan. 25, 2022).

28. Shahrezaei, V. & Swain, P. S. Analytical Distributions for Stochastic Gene Expression. Proceedings of the National Academy of Sciences of the United States of America 105, 17256–17261 (Nov. 11, 2008).

29. Gu, J. et al. Scalable Inference and Identifiability of Kinetic Parameters for Transcriptional Bursting from Single Cell Data. Bioinformatics 41 (ed Mathelier, A.) btaf581 (Nov. 1, 2025).

30. Kim, J. K. & Marioni, J. C. Inferring the Kinetics of Stochastic Gene Expression from Single-Cell RNA-Sequencing Data. Genome Biology 14, R7 (Jan. 28, 2013).

31. Rummel, T., Sakellaridi, L. & Erhard, F. grandR: A Comprehensive Package for Nucleotide Conversion RNA-Seq Data Analysis. Nature Communications 14, 3559 (June 15, 2023).

32. Vogt, M. et al. Targeting MYC Effector Functions in Pancreatic Cancer by Inhibiting the ATPase RUVBL1/2. Gut 73, 1509–1528 (Aug. 8, 2024).

33. Lorenzin, F. et al. Different Promoter Affinities Account for Specificity in MYC-Dependent Gene Regulation. eLife 5, e15161 (July 27, 2016).

34. Weikum, E. R., Knuesel, M. T., Ortlund, E. A. & Yamamoto, K. R. Glucocorticoid Receptor Control of Transcription: Precision and Plasticity via Allostery. Nature Reviews Molecular Cell Biology 18, 159–174 (Mar. 2017).

35. Popp, A. P., Hettich, J. & Gebhardt, J. C. M. Altering Transcription Factor Binding Reveals Comprehensive Transcriptional Kinetics of a Basic Gene. Nucleic Acids Research 49, 6249–6266 (June 21, 2021).

36. Carroll, P. A., Freie, B. W., Mathsyaraja, H. & Eisenman, R. N. The MYC Transcription Factor Network: Balancing Metabolism, Proliferation and Oncogenesis. Frontiers of Medicine 12, 412–425 (Aug. 2018).

37. Stine, Z. E., Walton, Z. E., Altman, B. J., Hsieh, A. L. & Dang, C. V. MYC, Metabolism, and Cancer. Cancer Discovery 5, 1024–1039 (Oct. 2015).

38. Campbell, K. J. & White, R. J. MYC Regulation of Cell Growth through Control of Transcription by RNA Polymerases I and III. Cold Spring Harbor Perspectives in Medicine 4, a018408–a018408 (May 1, 2014).

39. Guo, J. et al. Sequence Specificity Incompletely Defines the Genome-Wide Occupancy of Myc. Genome Biology 15, 482 (2014).

40. Wendlinger, S. et al. Blood Eosinophils Are Associated with Efficacy of Targeted Therapy in Patients with Advanced Melanoma. Cancers 14, 2294 (May 4, 2022).

41. Berg, K., et al. Identifying Modulators of Cellular Responses by Heterogeneity-Sequencing http://biorxiv.org/lookup/doi/10.1101/2024.10.28.620481 (2026). Pre-published.

