## Supplementary Note: Mathematical and computational details for buRst for "Inferring transcriptional bursting dynamics from scSLAM-seq data"

### Overview

This Supplementary Note provides mathematical derivations, numerical implementation details, simulation procedures, and analysis-specific reproducibility information for the `buRst` framework. The notation follows the main Methods: $B$ denotes the mean burst size, $k_m$ the burst-arrival frequency, $d$ the RNA degradation rate, and $\lambda = Bk_m$ the mean production rate. In code, these quantities correspond to `B`, `km`, and `d`.

#### 1 Discrete labeling likelihoods from read-level conversion evidence

##### 1.1 GRAND-SLAM measurement model

Read-level T→C conversion evidence was evaluated using the GRAND-SLAM measurement model [1]. For each molecule, the model uses the number of informative uridine positions, the number of observed T→C conversions, and conversion/error parameters to evaluate the likelihood of the observed read evidence under old RNA and newly synthesized RNA states.

For a gene–cell observation with total count $N_i$, the number of newly synthesized molecules is latent:

$$K_i \in \{0, 1, \dots, N_i\}.$$

For $N_i > 0$, the mixture log-likelihood was evaluated at new-RNA fractions $\theta = k/N_i$, giving a discrete log-likelihood profile

$$\ell_i(k), \qquad k = 0, \dots, N_i.$$

This profile was used as an approximate measurement likelihood over possible new RNA counts during kinetic inference.

##### 1.2 Profile centering, clipping, and quantization

Only relative measurement likelihoods are needed during kinetic inference. Each profile was centered at its maximum:

$$\widetilde{\ell}_i(k) = \ell_i(k) - \max_{r \in \{0,\dots,N_i\}} \ell_i(r). \tag{1}$$

The corresponding non-negative likelihood loss was

$$D_i(k) = \min\left[\max\left\{-\widetilde{\ell}_i(k), 0\right\}, R_{\text{clip}}\right]. \tag{2}$$

In the analyses described here, $R_{\text{clip}} = 60$ unless stated otherwise. Clipping bounds the range that must be stored while affecting only values with a likelihood less than $\exp(-R_{\text{clip}})$ relative to the profile maximum. For $R_{\text{clip}} = 60$, this corresponds to a relative likelihood below approximately $8.8 \times 10^{-27}$, so these values make a negligible contribution during inference. To reduce storage requirements and memory-transfer costs, the clipped profile was quantized to an unsigned 16-bit scale:

$$q_i(k) = \text{round}\left[65535 \frac{D_i(k)}{R_{\text{clip}}}\right]. \tag{3}$$

During inference, the stored integer value was converted back to

$$D_i(k) = q_i(k) R_{\text{clip}}/65535.$$

This representation requires 2 bytes per profile entry, compared with 4 bytes for single-precision or 8 bytes for double-precision floating-point storage. Thus, for $M$ total profile entries, the profiles require approximately $2M$ bytes, excluding file-format overhead. Because all profiles were centered, quantization approximately preserves the relative likelihood information required by the kinetic likelihood.

##### 1.3 HDF5 layout

Profiles were stored gene-wise in an HDF5 file. For each gene, the cell-specific profiles were concatenated into a single integer vector under

`/genes/<gene>`.

A cell with count $N_i > 0$ contributed $N_i + 1$ entries, whereas a cell with zero counts contributed an empty profile. The corresponding total counts and profile lengths were stored under

`/genes_meta/<gene>/counts_reads` and `/genes_meta/<gene>/lens_reads_plus1`.

Gene and cell identifiers were stored in the HDF5 metadata group. Datasets were chunked and compressed to permit efficient gene-wise retrieval during downstream inference.

#### 2 Effective transcriptional bursting model

##### 2.1 Model definition

Expression of RNA molecules ($M$) was modeled as an effective instantaneous bursting process. Burst events occurred according to a Poisson process with rate $k_m$. At each event, a random burst size $\mathcal{B}$ was drawn, and the RNA count increased by its realized value $b$:

$$\varnothing \xrightarrow{k_m} bM, \qquad b \text{ is a realization of } \mathcal{B}, \qquad M \xrightarrow{d} \varnothing. \tag{4}$$

The burst-size random variable $\mathcal{B}$ followed a geometric distribution on $\{0, 1, 2, \ldots\}$:

$$\Pr(\mathcal{B} = z) = \frac{1}{1+B}\left(\frac{B}{1+B}\right)^z, \qquad z = 0, 1, 2, \ldots, \tag{5}$$

where

$$B = \mathbb{E}[\mathcal{B}] \tag{6}$$

is the mean burst size. The mean RNA production rate was therefore

$$\lambda = k_m \mathbb{E}[\mathcal{B}] = B k_m. \tag{7}$$

Equivalently, when the current RNA count is $m$, a burst of size $z$ produces the transition

$$m \longrightarrow m + z \tag{8}$$

at rate

$$k_m \Pr(\mathcal{B} = z). \tag{9}$$

Each RNA molecule degraded independently at rate $d$, so that the total degradation rate in a state containing $m$ RNA molecules was $dm$:

$$m \longrightarrow m - 1 \qquad \text{at rate } dm. \tag{10}$$

The degradation rate was related to the RNA half-life by

$$d = \frac{\ln 2}{t_{1/2}}. \tag{11}$$

The model is an effective compound-Poisson burst model rather than an explicit two-state promoter-switching model. In particular, it does not contain separate promoter ON and OFF states or an explicit active-state duration. Instead, all RNA molecules associated with a burst are taken to be produced instantaneously. The model can therefore be interpreted as the short-burst limit of a more detailed promoter-switching process, in which the duration of an active transcriptional episode is short relative to the RNA lifetime and the timescales resolved by the experiment.

##### 2.2 Burst-size distribution

The burst size $\mathcal{B}$ followed a geometric distribution on $\{0, 1, 2, \ldots\}$. Using the standard success-probability parameter $p \in (0, 1]$, its probability mass function is

$$\mathbb{P}(\mathcal{B} = z) = p(1-p)^z, \qquad z = 0, 1, 2, \ldots. \tag{12}$$

Under this convention, $\mathcal{B}$ counts the number of RNA molecules produced before termination of a burst. Its mean is

$$\mathbb{E}[\mathcal{B}] = \frac{1-p}{p}. \tag{13}$$

For the kinetic model, it is more convenient to parameterize the distribution by the mean burst size

$$B = \mathbb{E}[\mathcal{B}] = \frac{1-p}{p}, \qquad p = \frac{1}{1+B}. \tag{14}$$

This parameterization is used below because $B$ has a direct biological interpretation and enters the mean RNA production rate as $\lambda = Bk_m$. Substituting $p = 1/(1+B)$ gives

$$\mathbb{P}(\mathcal{B} = z) = \frac{1}{1+B}\left(\frac{B}{1+B}\right)^z, \qquad z = 0, 1, 2, \ldots. \tag{15}$$

The probability-generating function is

$$F(x) = \mathbb{E}[x^{\mathcal{B}}] = \frac{p}{1-(1-p)x} = \frac{1}{1+B(1-x)}. \tag{16}$$

The moments used below are

$$\mathbb{E}[\mathcal{B}] = B, \qquad \mathbb{E}[\mathcal{B}^2] = B + 2B^2, \qquad \mathbb{E}[\mathcal{B}(\mathcal{B}-1)] = 2B^2. \tag{17}$$

#### 2.3 Newly synthesized RNA

The generating-function derivation in this and the following subsection adapts the analytical approach of Singh and Bokes [2] to the time-dependent distributions of newly synthesized and pre-existing RNA after labeling onset. Let $P_{\rm new}(m,t)$ denote the probability of observing $m$ molecules synthesized after labeling onset. The chemical master equation is

$$\frac{\partial P_{\rm new}(m,t)}{\partial t} = k_m \left[ \sum_{z=0}^{m} \mathbb{P}(\mathcal{B}=z) P_{\rm new}(m-z,t) - P_{\rm new}(m,t) \right] + d\left[(m+1)P_{\rm new}(m+1,t) - mP_{\rm new}(m,t)\right]. \quad (18)$$

At labeling onset,

$$P_{\rm new}(m,0) = \delta_{m0}.$$

Define the probability-generating function

$$G_{\rm new}(x,t) = \sum_{m=0}^{\infty} x^m P_{\rm new}(m,t). \quad (19)$$

Multiplying Eq. (18) by $x^m$ and summing over $m$ gives

$$\frac{\partial G_{\rm new}}{\partial t} = k_m\big(F(x)-1\big)G_{\rm new} + d(1-x)\frac{\partial G_{\rm new}}{\partial x}. \quad (20)$$

With $u = x-1$ and

$$\varphi_{\rm new}(u,t) = \log G_{\rm new}(1+u,t),$$

the factorial-cumulant generating function satisfies

$$\frac{\partial \varphi_{\rm new}}{\partial t} + du\frac{\partial \varphi_{\rm new}}{\partial u} = k_m\left(M(u)-1\right), \qquad M(u) = F(1+u). \quad (21)$$

Solving by characteristics gives

$$\varphi_{\rm new}(u,t) = k_m \int_0^t \left[ M\left(ue^{-ds}\right) - 1 \right] ds. \quad (22)$$

For the geometric burst-size distribution,

$$M(u) = \frac{1}{1-Bu}, \quad (23)$$

and therefore

$$\varphi_{\rm new}(u,t) = \frac{k_m}{d}\log\left(\frac{1-Bue^{-dt}}{1-Bu}\right). \quad (24)$$

Returning to $x = 1+u$,

$$G_{\rm new}(x,t) = \left[\frac{1-B(x-1)e^{-dt}}{1-B(x-1)}\right]^{k_m/d}. \quad (25)$$

In the limit $t\to\infty$, the newly synthesized distribution approaches the stationary total RNA distribution:

$$G_{\rm ss}(x) = \left[1-B(x-1)\right]^{-k_m/d}. \quad (26)$$

#### 2.4 Pre-existing RNA

Let $P_{\text{old}}(m,t)$ denote the distribution of molecules already present at labeling onset and surviving until time $t$. No new molecules enter this population, so its chemical master equation is

$$\frac{\partial P_{\text{old}}(m,t)}{\partial t} = d\left[(m+1)P_{\text{old}}(m+1,t) - mP_{\text{old}}(m,t)\right]. \tag{27}$$

If $G_{\text{old}}^{(0)}(x)$ is the initial generating function, then binomial thinning gives

$$G_{\text{old}}(x,t) = G_{\text{old}}^{(0)}\left[1 + (x-1)e^{-dt}\right]. \tag{28}$$

If the initial population is stationary, then

$$G_{\text{old}}(x,t) = \left[1 - Be^{-dt}(x-1)\right]^{-k_m/d}. \tag{29}$$

#### 2.5 Moments

Let

$$A = \frac{k_m B}{d} = \frac{\lambda}{d}, \qquad r(t) = e^{-dt}.$$

For newly synthesized RNA,

$$\mathbb{E}[K_{\text{new}}(t)] = A\left(1 - r(t)\right) \tag{30}$$

and

$$\text{Var}[K_{\text{new}}(t)] = A\left[1 - r(t) + B\left(1 - r(t)^2\right)\right]. \tag{31}$$

If the old RNA population has initial mean $\mu_0$ and variance $V_0$, then binomial thinning gives

$$\mathbb{E}[K_{\text{old}}(t)] = \mu_0 r(t) \tag{32}$$

and

$$\text{Var}[K_{\text{old}}(t)] = \mu_0 r(t)\left(1 - r(t)\right) + V_0 r(t)^2. \tag{33}$$

Under a stationary initial condition,

$$\mu_0 = A, \qquad V_0 = A(1+B),$$

so

$$\mathbb{E}[K_{\text{old}}(t)] = Ar(t), \qquad \text{Var}[K_{\text{old}}(t)] = A\left[r(t) + Br(t)^2\right]. \tag{34}$$

The old and new RNA populations are independent under the model. Therefore, when the system starts at steady state, the total RNA distribution remains stationary while molecules are progressively partitioned between old and new RNA.

#### 3 Numerical evaluation of RNA count distributions

##### 3.1 FFT inversion of probability-generating functions

The probability mass functions of newly synthesized, pre-existing, and total RNA were evaluated numerically from their probability-generating functions. For a distribution with PGF

$$G(x,t) = \sum_{m=0}^{\infty} P(m,t)x^m,$$

we evaluated $G$ at $N$ roots of unity

$$x_j = \exp\left(-\frac{2\pi ij}{N}\right), \qquad j = 0,\dots,N-1. \tag{35}$$

The first $N$ coefficients were recovered by inverse discrete Fourier transform:

$$\widetilde{P}(m,t) = \frac{1}{N}\sum_{j=0}^{N-1} G(x_j,t)\exp\left(\frac{2\pi ijm}{N}\right), \qquad m = 0,\dots,N-1. \tag{36}$$

For finite $N$, the coefficients satisfy

$$\widetilde{P}(m,t) = \sum_{\ell=0}^{\infty} P(m+\ell N,t), \tag{37}$$

so probability mass above $N-1$ can wrap around. The support was therefore selected adaptively.

##### 3.2 Numerically stable log-PGF evaluation

For each Fourier point, the implementation evaluated the log-PGF

$$\ell_j = \log G(x_j,t).$$

To avoid numerical overflow or underflow, the largest real component was subtracted:

$$c = \max_j \mathrm{Re}(\ell_j), \qquad A_j = \exp(\ell_j - c). \tag{38}$$

The inverse FFT was applied to $A_j$. The common scaling factor does not affect the normalized PMF. After inversion, the real component was retained, negative values were set to zero, and the coefficients were normalized to sum to one. Invalid or non-finite numerical evaluations were reported as errors; absolute-value and uniform-distribution fallbacks were not used. The minimum signed coefficient and the total magnitude of negative coefficients were recorded as numerical diagnostics before clipping. For the stationary distribution, the log-PGF was evaluated analytically:

$$\log G_{\mathrm{ss}}(x) = -\frac{k_m}{d}\log\left[1 - B(x-1)\right]. \tag{39}$$

For finite labeling times, the real and imaginary parts of the integral in Eq. (22) were evaluated numerically. For arbitrary empirical initial old RNA distributions with support values $n_r$ and probabilities $p_r^{(0)}$, the old RNA log-PGF was evaluated as

$$\log G_{\mathrm{old}}(x,t) = \operatorname{logsumexp}_r \left\{ n_r \log \left[ 1 - e^{-dt} + e^{-dt} x \right] + \log p_r^{(0)} \right\}. \tag{40}$$

Initial support points with zero probability were excluded before log-PGF evaluation. For a support point with $n_r = 0$, the term $n_r \log[1 - e^{-dt} + e^{-dt}x]$ was defined as zero, including when its logarithm argument was zero. The resulting numerical PMF was normalized after inversion.

##### 3.3 Adaptive support selection

The initial support size was selected from the analytical mean $\mu(t)$ and standard deviation $\sigma(t)$:

$$N_{\mathrm{start}} = \max \left\{ N_{\mathrm{user}}, \lceil \mu(t) + 4\sigma(t) \rceil + 1 \right\}. \tag{41}$$

After inversion, a tail diagnostic was calculated as the summed probability in the final $L$ support bins:

$$\epsilon_{\mathrm{tail}}^{(N)} = \sum_{m=N-L}^{N-1} \widetilde{P}(m,t). \tag{42}$$

The tail diagnostic used the final ten support bins, or all bins when the support contained fewer than ten entries. During kinetic fitting, the tolerance was set to $10^{-8}$ for both newly synthesized and pre-existing RNA distributions. If the diagnostic was not below this tolerance, the support was enlarged according to

$$N \leftarrow \lceil 1.5 N \rceil$$

and the PMF was recalculated. At most four support evaluations were performed, including the initial evaluation. The support-size stopping thresholds were $2^{20}$ for newly synthesized RNA and $2^{18}$ for pre-existing RNA. These thresholds were checked after evaluation, so the initial support or a support enlargement could exceed them. If the tolerance was not reached before an iteration or support-size limit, the last normalized PMF was returned together with its tail diagnostic; this did not automatically reject the fit. The tail diagnostic is not a strict bound on omitted probability, because circular aliasing can move probability mass to lower counts. It was used as a practical diagnostic of support adequacy.

##### 3.4 Total RNA distribution

The total count is

$$K_{\mathrm{tot}}(t) = K_{\mathrm{new}}(t) + K_{\mathrm{old}}(t).$$

Because old and new RNA are independent under the model, the total distribution is the convolution

$$P_{\mathrm{tot}}(n,t) = \sum_{k=0}^{n} P_{\mathrm{new}}(k,t) P_{\mathrm{old}}(n-k,t). \tag{43}$$

The convolution was evaluated by zero-padding the two PMFs, multiplying their discrete Fourier transforms, and applying an inverse FFT. For input vectors of lengths $N_1$ and $N_2$, the transform length was

$$N_{\mathrm{conv}} = 2^{\lceil \log_2(N_1 + N_2 - 1) \rceil}.$$

##### 3.5 Analytical checks

As a consistency check of the preceding derivations, we verified that the decomposition into pre-existing and newly synthesized RNA preserves the stationary total RNA distribution. When the total RNA count is initially at stationarity, the probability-generating function of the sum of the old and new RNA populations is the product of their generating functions:

$$\begin{aligned} G_{\text{old}}(x,t)G_{\text{new}}(x,t) &= \left[1 - Be^{-dt}(x-1)\right]^{-k_m/d} \left[\frac{1 - Be^{-dt}(x-1)}{1 - B(x-1)}\right]^{k_m/d} \\ &= [1 - B(x-1)]^{-k_m/d} = G_{\text{ss}}(x). \end{aligned} \tag{44}$$

Thus, the total RNA distribution remains stationary at every labeling time. This identity was used as an analytical check of FFT inversion and convolution.

### 4 Kinetic likelihood, initialization, and uncertainty

##### 4.1 Parameterization

For each gene and condition, the biological parameter vector is

$$\boldsymbol{\theta} = (B, k_m, d).$$

Optimization was performed in transformed coordinates

$$\boldsymbol{\eta} = (\log \lambda, \log B, \log d), \qquad \lambda = Bk_m. \tag{45}$$

The burst-arrival frequency was recovered as

$$k_m = \lambda / B.$$

This transformation enforces positivity and places the mean production rate directly in the optimization vector.

##### 4.2 Profile-based likelihood

Let $N_i$ denote the observed total RNA count in cell $i$. The number of newly synthesized RNA molecules, denoted by $k$, is not observed directly. Instead, the measurement procedure provides a likelihood profile over the possible values $k = 0, \ldots, N_i$. We write the centered relative log-likelihood as $E_i(k)$, with

$$\max_{0 \le k \le N_i} E_i(k) = 0. \tag{46}$$

Thus, $\exp E_i(k)$ is proportional to the likelihood of the measurement in cell $i$ conditional on the cell containing $k$ newly synthesized RNA molecules. Centering removes a cell-specific multiplicative constant that does not depend on the kinetic parameters. For a proposed kinetic parameter vector $\boldsymbol{\theta}$, the model defines the probability mass functions

$$P_{\text{new}}(k \mid \boldsymbol{\theta}) \qquad \text{and} \qquad P_{\text{old}}(r \mid \boldsymbol{\theta}) \tag{47}$$

for the new and old RNA populations. Because the observed total count satisfies

$$N_i = k + r, \tag{48}$$

the likelihood contribution of cell $i$, up to the profile-specific constant removed by centering, is

$$L_i(\boldsymbol{\theta}) = \sum_{k=0}^{N_i} \exp E_i(k) P_{\text{new}}(k \mid \boldsymbol{\theta}) P_{\text{old}}(N_i - k \mid \boldsymbol{\theta}). \tag{49}$$

This expression marginalizes over all possible allocations of the observed total count into newly synthesized and pre-existing RNA. The model probability of observing total count $N_i$, irrespective of the measurement profile, is the convolution

$$P_{\text{tot}}(N_i \mid \boldsymbol{\theta}) = \sum_{k=0}^{N_i} P_{\text{new}}(k \mid \boldsymbol{\theta}) P_{\text{old}}(N_i - k \mid \boldsymbol{\theta}). \tag{50}$$

Conditional on observing the total count $N_i$, the model-implied probability that $k$ of these molecules are newly synthesized is

$$\pi_{\boldsymbol{\theta}}(k \mid N_i) = \frac{P_{\text{new}}(k \mid \boldsymbol{\theta}) P_{\text{old}}(N_i - k \mid \boldsymbol{\theta})}{P_{\text{tot}}(N_i \mid \boldsymbol{\theta})}. \tag{51}$$

This conditional distribution is introduced only to separate the information provided by the total RNA count from the additional information provided by the measurement profile. Substituting Eq. (51) into Eq. (49) gives

$$L_i(\boldsymbol{\theta}) = P_{\text{tot}}(N_i \mid \boldsymbol{\theta}) \sum_{k=0}^{N_i} \exp E_i(k) \pi_{\boldsymbol{\theta}}(k \mid N_i). \tag{52}$$

The first factor evaluates how well the model explains the observed total count. The second factor evaluates how well the model-implied allocation between new and old RNA agrees with the measurement profile. Summing the logarithm of Eq. (52) over cells gives

$$\ell(\boldsymbol{\theta}) = \ell_{\text{tot}}(\boldsymbol{\theta}) + \ell_{\text{meas}}(\boldsymbol{\theta}), \tag{53}$$

where

$$\ell_{\text{tot}}(\boldsymbol{\theta}) = \sum_{N} c_N \log \left( P_{\text{tot}}(N \mid \boldsymbol{\theta}) \right), \tag{54}$$

and

$$\ell_{\text{meas}}(\boldsymbol{\theta}) = \sum_{i:N_i>0} \log \left( \sum_{k=0}^{N_i} \exp E_i(k) \pi_{\boldsymbol{\theta}}(k \mid N_i) \right). \tag{55}$$

Here, $c_N$ is the number of cells with observed total count $N$, and only observed count values with $c_N > 0$ need to be included in the sum. The factorization above describes the information supplied by total counts and labeling evidence. Numerically, the joint likelihood was evaluated directly as

$$\ell(\boldsymbol{\theta}) = \sum_{i} \operatorname{logsumexp}_{0 \le k \le N_i} \left\{ E_i(k) + \log P_{\text{new}}(k \mid \boldsymbol{\theta}) + \log P_{\text{old}}(N_i - k \mid \boldsymbol{\theta}) \right\}. \tag{56}$$

This avoids division by the total-count probability and does not add probability floors to the profile-based likelihood.

The new and old RNA PMFs were normalized over their full, adaptively selected FFT supports before entries corresponding to counts from zero to the largest observed total count were extracted. When the total RNA PMF was required, it was obtained by convolution of the full component PMFs and normalized on its full convolution support before extracting entries.

Cells were grouped by observed total count for efficient evaluation of the measurement-profile matrices. For $N_i = 0$, the contribution was $\log P_{\text{new}}(0) + \log P_{\text{old}}(0)$, with no additional labeling-evidence term.

##### 4.3 MAP or known component-count likelihood

For point-estimate analyses, the integer new RNA count was reconstructed using the maximum-a-posteriori NTR estimate from the labeling data

$$\widehat{K}_{\text{new},i} = \operatorname{round}\left(\widehat{\text{NTR}}_i N_i\right).$$

The corresponding old count was

$$\widehat{K}_{\text{old},i} = \max\left(N_i - \widehat{K}_{\text{new},i}, 0\right).$$

For truth-count analyses, the exact simulated new and old RNA counts were used. The component-count likelihood was

$$\ell_{\text{count}}(\boldsymbol{\theta}) = \sum_i \log P_{\text{new}}\left(K_{\text{new},i} \mid \boldsymbol{\theta}\right) + \sum_i \log P_{\text{old}}\left(K_{\text{old},i} \mid \boldsymbol{\theta}\right). \tag{57}$$

A probability floor of $10^{-300}$ was used. The total-count PMF was not included by default because the total count is determined by the two component counts.

##### 4.4 Beta approximation to labeling uncertainty

For the beta-approximation analysis, the uncertainty in the new-to-total RNA ratio of cell $i$ was represented as

$$X_i \sim \text{Beta}(\alpha_i, \beta_i), \tag{58}$$

where $X_i$ denotes the new-to-total ratio and $\alpha_i$ and $\beta_i$ are the cell-specific shape parameters. To obtain a discrete likelihood over the possible newly synthesized RNA counts $k = 0, \ldots, N_i$, the beta distribution was integrated over the interval of ratio values that would round to count $k$. For cells with $N_i > 0$, the resulting probability mass was

$$w_i(k) = F_{\text{Beta}}\left(\min\left(1, \frac{k+1/2}{N_i}\right); \alpha_i, \beta_i\right) - F_{\text{Beta}}\left(\max\left(0, \frac{k-1/2}{N_i}\right); \alpha_i, \beta_i\right). \tag{59}$$

where $F_{\text{Beta}}$ is the beta cumulative distribution function. The boundary intervals for $k = 0$ and $k = N_i$ were extended to $0$ and $1$, respectively. For numerical evaluation, the masses were floored at $10^{-300}$

before taking logarithms. The discrete masses were converted to a centered relative log-likelihood profile,

$$E_i^{\mathrm{Beta}}(k) = \log w_i(k) - \max_{0 \le r \le N_i} \log w_i(r), \tag{60}$$

and the corresponding negative log-likelihood losses were clipped at $R_{\mathrm{clip}} = 60$, as for the full measurement profiles. The resulting profile was then supplied to the same profile-based kinetic likelihood used for the full-likelihood analysis. Thus, the beta approximation differed from the full analysis only in the representation of per-cell labeling uncertainty, while the kinetic model and fitting procedure were unchanged. Cells with $N_i = 0$ have only the possible new RNA count $k = 0$ and therefore did not contribute a labeling-uncertainty profile.

#### 4.5 Steady-state and non-steady-state models

For steady-state fits, the RNA population immediately before labeling was assumed to follow the stationary distribution. This distribution was used as the initial old RNA population. During the labeling interval, each pre-existing RNA molecule was retained independently with probability

$$\exp\left(-dt_{\mathrm{label}}\right), \tag{61}$$

corresponding to survival without degradation. Thus, conditional on an initial old RNA count of $n$, the number remaining after the labeling interval followed a binomial distribution with parameters $n$ and $\exp(-dt_{\mathrm{label}})$. Newly synthesized RNA was generated over the same labeling interval according to the bursting model.

For non-steady-state fits, the initial old RNA distribution was constructed from total RNA counts in a reference condition:

$$\widehat{p}_0(n) = \frac{1}{C_{\mathrm{ref}}} \sum_{i=1}^{C_{\mathrm{ref}}} \mathbf{1}\{N_i^{\mathrm{ref}} = n\}. \tag{62}$$

The formulation described here uses equal modeled propagation and labeling intervals, $t_{\mathrm{diff}} = t_{\mathrm{label}}$. The old RNA distribution at the fitted observation was obtained by thinning the empirical reference distribution over this interval:

$$G_{\mathrm{old}}(x, t_{\mathrm{label}}) = \widehat{G}_0\left[1 + (x-1)e^{-dt_{\mathrm{label}}}\right]. \tag{63}$$

New RNA was generated over the same interval using the kinetic parameters of the fitted condition. The empirical reference distribution was treated as fixed.

For the MYC-depletion analysis, $t_{\mathrm{diff}} = t_{\mathrm{label}} = 2$ h. The empirical total RNA distribution from the corresponding DMSO condition was used to approximate the RNA population at labeling onset. Changes in this distribution during the preceding 2 h auxin pretreatment were not modeled explicitly.

#### 4.6 Initialization of numerical optimization

The following procedure was used to obtain starting parameter values for numerical optimization. These initial values were used for all likelihood representations, including the full-profile, point-estimate, and beta approximation analyses. Estimated new RNA counts were used only for initialization and did not replace the measurement profiles in the fitted likelihood.

Let

$$\widehat{f} = \frac{\sum_i \widehat{K}_{\text{new},i}}{\max(1, \sum_i N_i)}$$

denote the aggregate estimated new RNA fraction. Under the steady-state model, the initial degradation-rate estimate was

$$d_0 = -\frac{1}{t_{\text{label}}} \log(1 - \widehat{f}). \tag{64}$$

For non-steady-state fits, fitted and reference abundances were compared on a per-cell mean scale. Let $C_{\text{fit}}$ and $C_{\text{ref}}$ denote the numbers of cells in the fitted and reference populations, respectively, and define

$$\overline{O} = \frac{1}{C_{\text{fit}}} \sum_{i=1}^{C_{\text{fit}}} \left(N_i - \widehat{K}_{\text{new},i}\right), \qquad \overline{F}_0 = \frac{1}{C_{\text{ref}}} \sum_{j=1}^{C_{\text{ref}}} N_j^{\text{ref}}.$$

For the equal modeled propagation and labeling intervals used here, and a positive reference mean, the initial degradation-rate estimate was

$$d_0 = -\frac{1}{t_{\text{label}}} \log\left[\min\left(\frac{\overline{O}}{\overline{F}_0}, 1\right)\right]. \tag{65}$$

Using per-cell means accounts for unequal cell numbers in the fitted and reference populations. This calculation supplies only a starting value for subsequent likelihood optimization.

If the initial degradation-rate estimate was non-finite or non-positive, it was replaced by $\ln 2/t_{\text{label}}$. The resulting value was rounded to five decimal places and restricted to

$$\frac{\ln 2}{50} \leq d_0 \leq \frac{\ln 2}{5/60}.$$

Initial $B$ and $k_m$ values were obtained by matching the empirical mean and second moment of estimated new RNA counts. For fixed $d_0$, the candidate burst frequency required by the mean equation was

$$k_m(B) = \frac{d_0 \widehat{m}}{B\left(1 - e^{-d_0 t_{\text{label}}}\right)}, \tag{66}$$

where $\widehat{m}$ is the empirical mean of estimated new RNA counts. The candidate $B_0$ was selected by minimizing the squared discrepancy between the observed and analytical transient second raw moments. For the empty initial condition, exact matching gives

$$B_0 = \frac{\widehat{v} - \widehat{m}}{\widehat{m}\left(1 + e^{-d_0 t_{\text{label}}}\right)}, \tag{67}$$

where $\widehat{v}$ is the empirical variance of estimated new RNA counts, when this expression is positive. Moment estimates were rejected when they were non-finite or non-positive, when $B_0$ was outside $[10^{-3}, 100]$, when $k_{m,0}$ was outside $[10^{-3}, 20]$, or when they implied $B_0 k_{m,0} < 10^{-2}$. The fallback was

$$B_0 = 0.1, \qquad \lambda_0 = \max(\widehat{m} d_0, 10^{-2}), \qquad k_{m,0} = \lambda_0 / B_0.$$

Accepted moment estimates of $B_0$ and $k_{m,0}$ were rounded to three decimal places. Before fitting, $B_0$ was restricted to $[10^{-3}, 100]$ and $k_{m,0}$ to $[10^{-3}, 10]$, and $\lambda_0 = B_0 k_{m,0}$ was recomputed after these restrictions.

#### 4.7 Optimization and fit filters

A conservative minimum FFT support was selected once from the initial parameters. The support covered the largest observed count and at least

$$\mu_{0,\mathrm{ss}} + 10\sigma_{0,\mathrm{ss}},$$

where $\mu_{0,\mathrm{ss}}$ and $\sigma_{0,\mathrm{ss}}$ are the stationary mean and standard deviation implied by the initial parameters. The initial minimum-support heuristic was capped at 20,000 states. The likelihood builder nevertheless enforced coverage of every observed count, requiring at least $1 + \max_i N_i$ states, and adaptive PMF evaluation could enlarge the support further. The transformed log-likelihood was maximized using Nelder–Mead with relative tolerance $10^{-6}$ and at most 1,000 iterations. Parameter proposals were rejected if any of

$$\lambda, \quad B, \quad d, \quad k_m$$

was non-finite, no larger than $10^{-12}$, or greater than $10^3$. Fits were attempted only for genes satisfying the information filters described in the main Methods. These filters were not applied to fits based on exact simulation truth.

#### 4.8 Laplace approximation and identifiability diagnostics

A numerical Hessian $H$ of the log-likelihood was evaluated at the optimum. The local covariance approximation in transformed space was

$$\widehat{\Sigma} = (-H)^{-1}. \tag{68}$$

The Hessian and covariance matrices were symmetrized. Eigenvalues of the covariance matrix smaller than $10^{-10}$ were replaced by $10^{-10}$ before simulation. Gaussian draws were generated as

$$\boldsymbol{\eta}^{(s)} \sim \mathcal{N}\left(\widehat{\boldsymbol{\eta}}, \widehat{\Sigma}\right), \tag{69}$$

and transformed to $\lambda^{(s)}$, $B^{(s)}$, $d^{(s)}$, and $k_m^{(s)} = \lambda^{(s)}/B^{(s)}$. Invalid transformed draws were discarded. The 2.5th, 50th, and 97.5th percentiles were reported. Under a locally flat prior in the transformed coordinates, these Gaussian distributions were interpreted as approximate posteriors. Identifiability was assessed from the observed information matrix $-H$. A ridge score was defined as

$$R_{\mathrm{ridge}} = \frac{\lambda_{\min}(-H)}{\lambda_{\max}(-H)}. \tag{70}$$

Parameter correlations were calculated in both $(\log \lambda, \log B, \log d)$ and $(\log B, \log k_m, \log d)$ coordinates.

Diagnostic flags were assigned hierarchically. Non-finite or non-positive observed-information eigenvalues, or failed covariance or fitting-coordinate correlation calculations, were flagged as `bad_hessian`. An observed-information eigenvalue ratio below 0.001 was flagged as `strong_tradeoff`. Otherwise, if the most strongly correlated pair in fitting coordinates was $\log \lambda$ and $\log d$, with absolute correlation at least 0.95, the fit was flagged as `d_related_tradeoff`. If the most strongly correlated pair in biological coordinates was $\log B$ and $\log k_m$, the fit was flagged as `strong_tradeoff_Bkm` for absolute correlation at least 0.97, or as `B_km_tradeoff` for absolute correlation at least 0.90. Remaining fits with an eigenvalue ratio below 0.01 were flagged as `moderate_tradeoff`; other fits were flagged as `ok`. Where fit-quality exclusions were applied, fits flagged as `bad_hessian` or `strong_tradeoff` were excluded. Other diagnostic flags did not independently trigger exclusion.

#### 4.9 Classification of kinetic fold-change responses

Kinetic changes between two conditions or time points were classified from the joint approximate posterior distribution of burst-frequency and burst-size $\log_2$ fold changes. For each Monte Carlo draw $s$, the joint fold-change vector was defined as

$$\boldsymbol{\Delta}^{(s)} = \begin{pmatrix} \Delta_{k_m}^{(s)} \\ \Delta_B^{(s)} \end{pmatrix}. \tag{71}$$

Evidence for a kinetic change was assessed using both the displacement and the posterior mass of the joint fold-change distribution. Let $\widehat{\boldsymbol{\mu}}_\Delta$ and $\widehat{\boldsymbol{\Sigma}}_\Delta$ denote the empirical mean vector and covariance matrix of the joint fold-change draws. For numerical stability, the covariance matrix was regularized as

$$\widehat{\boldsymbol{\Sigma}}_{\Delta,\mathrm{reg}} = \widehat{\boldsymbol{\Sigma}}_\Delta + \delta I_2, \qquad \delta = \max\left\{ 10^{-8}\frac{\mathrm{tr}\left(\widehat{\boldsymbol{\Sigma}}_\Delta\right)}{2},\ 10^{-10} \right\}, \tag{72}$$

where $I_2$ is the $2 \times 2$ identity matrix. The squared Mahalanobis distance from the no-change point was then calculated as

$$D^2 = \widehat{\boldsymbol{\mu}}_\Delta^\mathsf{T} \widehat{\boldsymbol{\Sigma}}_{\Delta,\mathrm{reg}}^{-1} \widehat{\boldsymbol{\mu}}_\Delta. \tag{73}$$

Joint evidence for a kinetic change required $D^2 > \chi^2_{2,0.95} \approx 5.99$, corresponding to the 95th percentile of a $\chi^2$ distribution with two degrees of freedom, and at least 90% of draws lay outside a no-change circle of radius 0.3:

$$\sqrt{\left(\Delta_{k_m}^{(s)}\right)^2 + \left(\Delta_B^{(s)}\right)^2} > 0.3. \tag{74}$$

For genes with a supported kinetic change, joint draws outside the no-change circle were assigned to mechanistic regions. Draws were assigned to the burst-frequency region when

$$|\Delta_B^{(s)}| < \frac{|\Delta_{k_m}^{(s)}|}{\sqrt{3}},$$

to the burst-size region when

$$|\Delta_B^{(s)}| > \sqrt{3}|\Delta_{k_m}^{(s)}|,$$

and to the combined region otherwise. These boundaries correspond to angles of $30^\circ$ and $60^\circ$ in the plane of absolute fold changes. Mechanistic probabilities were calculated conditional on lying outside the no-change circle, with at least 60% required to support a region. Opposing responses were evaluated separately. An opposing response was supported when at least 60% of draws outside the no-change circle had opposite-sign fold changes and the combined opposing region was the most populated individual directional region. Among opposing draws, a component was considered dominant when its absolute fold change was at least 1.2 times that of the other component in at least 70% of draws. Supported opposing responses were classified as frequency dominated with an opposing size shift, size dominated with an opposing frequency shift, or as an opposing trade-off without supported dominance. These assignments took precedence over the ordinary mechanistic-region classification. Other genes with a supported kinetic change were assigned to the frequency, size, or combined class when the corresponding conditional probability reached 60%; otherwise, their mechanism was considered unresolved. Genes failing either the joint-evidence or practical-change criterion were classified as showing no supported change.

### 5 Simulation framework

#### 5.1 Gillespie simulation of RNA counts

RNA counts were simulated using an exact stochastic simulation algorithm implemented in C++ through `Rcpp`. For each simulated cell, the state was

$$n_{\text{new}}(t), \qquad n_{\text{old}}(t), \qquad n_{\text{total}}(t) = n_{\text{new}}(t) + n_{\text{old}}(t).$$

Burst events occurred with propensity $k_m$. At a burst event,

$$n_{\text{new}} \to n_{\text{new}} + \mathcal{B},$$

where $\mathcal{B}$ was sampled from Eq. (15). Degradation propensities were

$$dn_{\text{new}}, \qquad dn_{\text{old}}.$$

Equivalently, degradation occurred with total propensity $d(n_{\text{new}} + n_{\text{old}})$, and the degraded molecule was assigned to the new or old population in proportion to the current population sizes. Waiting times were sampled from an exponential distribution with rate $k_m + d(n_{\text{new}} + n_{\text{old}})$. If the next event time exceeded the end of the simulation interval, the trajectory was terminated. At labeling onset, $n_{\text{new}}(0) = 0$. For stationary simulations, $n_{\text{old}}(0)$ was sampled from the stationary PMF calculated by FFT. For non-stationary simulations, initial old RNA counts were supplied directly.

#### 5.2 Construction of simulated `grandR` objects

Simulated new and old RNA counts were assembled into sparse gene-by-cell matrices. Total counts and true NTR values were calculated as

$$C_{gc} = C_{gc}^{\text{new}} + C_{gc}^{\text{old}}$$

and

$$\text{NTR}_{gc} = \frac{C_{gc}^{\text{new}}}{C_{gc}}, \qquad C_{gc} > 0. \tag{75}$$

The matrices were converted into a `grandR` object containing `count`, `ntr`, `new`, and `old` slots. Gene metadata included the true values of $B$, $k_m$, and $d$, and cells were assigned unique 20-nucleotide barcodes.

#### 5.3 Simulation of read-level labeling evidence

To generate GRAND-SLAM-like labeling evidence, the total molecule count of each gene–cell observation was used as the number of simulated reads. Read-specific uridine fractions were sampled as

$$\rho_r \sim \text{Beta}(\alpha_U, \beta_U),$$

where the shape parameters were calculated from the requested mean $\mu_U$ and standard deviation $\sigma_U$:

$$\alpha_U = \mu_U \left[ \frac{\mu_U(1-\mu_U)}{\sigma_U^2} - 1 \right], \qquad \beta_U = \frac{1-\mu_U}{\mu_U} \alpha_U. \tag{76}$$

For a read of length $L$, the number of uridine positions was sampled as

$$U_r \sim \text{Binomial}(L, \rho_r).$$

The validation and perturbation benchmarks used the settings $L = 100$ nucleotides, $\mu_U = 0.25$, and $\sigma_U = 0.05$, corresponding to $\alpha_U = 18.5$ and $\beta_U = 55.5$. Reads with no uridine positions did not contribute conversion evidence. Conversion-count evidence was generated using `grandR::CreateMixMatrix`, parameterized by the simulated new-to-total RNA ratio, the uridine-count histogram, background error parameter $p_{\text{old}} = 10^{-4}$, and new RNA conversion parameter $p_{\text{new}} = 0.04$. The conversion parameter used during fitting was also set to 0.04 in the primary validation and perturbation benchmark. For each observation with positive total count $N_i$, the measurement likelihood was evaluated over all possible new RNA counts $k = 0, \ldots, N_i$. Profiles were centered, clipped, and quantized as described in Section 1. Observations with positive counts but no informative uridine positions were assigned a flat measurement profile; zero-count observations contributed an empty profile. NTR estimates were obtained using `grandR::fit.ntr`. When requested, beta-approximation parameters were fitted using `beta.approx=TRUE` and stored in the corresponding `grandR` slots. The perturbation benchmark did not request beta-approximation fitting.

#### 5.4 Self-contained output formats

Simulated data were exported as a self-contained, grand3-style `<prefix>.targets` directory. The directory contained

- `4sU.Binom.discrete.h5`, containing quantized discrete likelihood profiles;
- `matrix.mtx.gz`, containing the sparse total-count matrix;
- `barcodes.tsv.gz`, containing cell identifiers; and
- `features.tsv.gz`, containing gene identifiers, gene symbols, and the feature type `Gene Expression`.

The matrix, barcode, and feature files form a 10x-compatible reference used to align count data with the HDF5 likelihood profiles. For comparison with `burstMCMC`, an additional tab-separated file was generated. Each line contained the gene identifier, cell barcode, total count, and simulated read evidence represented as repeated `k/n` entries, where $n$ is the number of uridine positions and $k$ is the number of observed conversions. In bundled output mode, this file was stored as `burstMCMC.txt` inside the `.targets` directory.

#### 5.5 Validation simulation based on sci-fate-derived parameters

Gene-specific generating parameters for the primary validation simulation were obtained from fits to the A549 0 h condition of the sci-fate dataset. Genes with fits flagged as `bad_hessian` or `strong_tradeoff` were excluded. The remaining 8,254 genes were simulated using their estimated burst sizes $B$, burst frequencies $k_m$, and degradation rates $d$. For each gene, 1,151 cells were simulated over a 2 h labeling interval. The simulation used random seed 1234. A separate perturbation benchmark assessed inference of kinetic changes under non-steady-state conditions. Reference parameters were taken from A549 0 h fits after excluding the same two diagnostic flags. Genes were ordered by Ensembl identifier before perturbation assignment. The benchmark contained 25 predefined perturbation combinations:

- one no-change combination, $(\Delta k_m, \Delta B) = (0, 0)$;
- six frequency-only combinations, $(\Delta k_m, \Delta B) = (\pm a, 0)$;
- six size-only combinations, $(\Delta k_m, \Delta B) = (0, \pm a)$;
- six concordant joint combinations, $(\Delta k_m, \Delta B) = (a, a)$ or $(-a, -a)$;
- six opposing joint combinations, $(\Delta k_m, \Delta B) = (a, -a)$ or $(-a, a)$,

where $a \in \{0.5, 1, 2\}$ and all changes are on the $\log_2$ scale. Each gene received one perturbation combination. The ordered list of 25 combination identifiers was repeated to the number of retained genes and randomly permuted using seed 1001. Consequently, the numbers of genes assigned to individual combinations differed by at most one. Assignment was balanced across combinations rather than across the five broad perturbation classes. Perturbed parameters were defined as

$$k_{m,2} = k_{m,0}\, 2^{\Delta k_m}, \qquad B_2 = B_0\, 2^{\Delta B}, \qquad d_2 = d_0.$$

The reference condition was simulated under steady-state kinetics for a 2 h labeling interval using 1,151 cells per gene. The perturbed condition was initialized directly from the gene-by-cell total RNA count matrix generated for the reference condition, with all initial molecules designated as pre-existing RNA. It was then propagated for 2 h under the perturbed kinetic parameters, using the same number of cells. Read-level labeling evidence was generated for both conditions using the settings described in Section 5.3. Beta-approximation fitting was disabled, and both discrete likelihood profiles and `burstMCMC`-compatible evidence were exported. Both condition-generation calls used random seed 1234 and parallel execution. The worker count was set through the workflow configuration and recorded in the archived run provenance. The simulated reference condition was fitted under the steady-state model. The perturbed condition was fitted under the non-steady-state model using the simulated reference total RNA distribution as its initial condition. Recovered burst-frequency and burst-size fold changes, uncertainty intervals, and mechanistic classifications were compared with their generating values.

##### 5.6 Reproducibility and parallel execution

When a seed was supplied, R's random-number generator was used throughout. The simulation wrapper selected the `L'Ecuyer-CMRG` generator to support reproducibility in simulations parallelized across genes. Read evidence calculations were parallelized across genes where applicable, while HDF5 writing was performed gene by gene.

#### 6 Validation likelihoods and benchmark against `burstMCMC`

##### 6.1 Full discrete likelihood

In the full-likelihood analysis, the stored Grand3 profile was used directly. For each cell with total count $N_i$, the profile over $k = 0, \ldots, N_i$ was combined with the model-predicted new and old RNA

PMFs as in Eq. (49). This approach marginalizes over labeling uncertainty instead of first estimating a single new RNA count.

##### 6.2 Point-estimate likelihood

In the point-estimate analysis, labeling uncertainty was replaced by a single integer estimate $K_{\mathrm{new},i}$, based on the NTR calculated by *grandR*. The old RNA count was set to

$$K_{\mathrm{old},i} = \max(N_i - K_{\mathrm{new},i}, 0).$$

The kinetic parameters were fitted using the component-count likelihood in Eq. (57).

##### 6.3 Beta approximation

In the beta-approximation analysis, per-cell NTR uncertainty was represented by

$$\Theta_i \sim \mathrm{Beta}(\alpha_i, \beta_i).$$

For a cell with total count $N_i > 0$, this continuous distribution was converted into a discrete probability mass over possible new RNA counts $k = 0, \ldots, N_i$. The mass assigned to $k$ was

$$\pi_i(k) = \int_{a_{ik}}^{b_{ik}} \mathrm{Beta}\left(\theta; \alpha_i, \beta_i\right)\, d\theta, \tag{77}$$

where

$$a_{ik} = \max\left(0, \frac{k - \frac{1}{2}}{N_i}\right), \qquad b_{ik} = \min\left(1, \frac{k + \frac{1}{2}}{N_i}\right). \tag{78}$$

Boundary intervals for $k = 0$ and $k = N_i$ were extended to 0 and 1, respectively. Masses were floored at $10^{-300}$ before taking logarithms. The discrete masses were converted to relative negative-log-likelihoods

$$g_i(k) = \max_r \log \pi_i(r) - \log \pi_i(k), \tag{79}$$

clipped at $R_{\mathrm{clip}} = 60$, and supplied to the same marginalized kinetic likelihood used for the full profiles.

##### 6.4 Parameter recovery and coverage

Parameter recovery was evaluated for

$$q \in \{\lambda, d, k_m, B\}$$

using

$$\varepsilon_q = \log_{10}\left(\frac{\widehat{q}}{q_{\mathrm{true}}}\right). \tag{80}$$

Empirical coverage was calculated as the fraction of genes for which the generating parameter value fell within the 95% Laplace-approximation interval.

##### 6.5 Benchmark against `burstMCMC`

The same simulated molecule counts and read-level conversion evidence were used for the `burstMCMC` benchmark [3]. The `burstMCMC`-compatible input file contained, for each gene and cell, the total UMI count together with the number of T→C conversions and informative uridine positions for each simulated molecule. The background and labeled-RNA conversion probabilities were fixed to the simulation values,

$$p_{\text{old}} = 10^{-4}, \qquad p_{\text{new}} = 0.04.$$

The labeling duration was supplied as 120 min. Because the `buRst` simulation did not include capture efficiency, the `burstMCMC` capture-efficiency parameter was fixed to

$$\alpha_i = 0.01$$

for all cells. The data were formatted using `burstMCMC::formatData`. A custom runner used the package functions for likelihood evaluation, prior specification, chain initialization, and proposal generation. Fitting was initiated under model 2 independently for each gene. Chains contained 5,000 stored iterations, reduced to 1,500 when the estimated expression scale, $\mathrm{ME} = \overline{\mathrm{UMI}}/\overline{\alpha}$, was at least 1,000. The corresponding numerical-failure monitoring windows were 500 and 100 iterations. When the recent-iteration check detected at least 25 or 5 iterations, respectively, for which the likelihood routine returned an `f_n_failure` indicator, the chain was discarded and restarted under model 3. Successful final chains from both models were included in downstream comparisons only when their final status was `ok`. Incomplete timeout or error chains were excluded, even if they contained some completed sampling iterations. Initialization used an expression parameter fixed at $\mathrm{ME}$, a log-uniform burst-rate prior between 1 and 10, and a normal lifetime prior centered on a labeling-based lifetime estimate with standard deviation equal to one-fifth of that estimate. Subsequent sampling used uniform priors on the expression and burst-rate parameters between 0 and 100,000 and on the lifetime parameter between 1 and 100,000 min. Genes were processed in parallel using the worker count specified in the workflow configuration and recorded in the archived run provenance. Elapsed time was checked during initialization, warm-up, and sampling against a 10,000 s limit per gene. Fits exceeding this limit at a check were recorded as timeouts and excluded from posterior comparisons. Recorded per-gene runtime included initialization and any chain restarts. Per-gene results were checkpointed so that an interrupted run could reuse saved attempts; reloading and summarizing a cached attempt did not replace its original sampling runtime. Before comparison with simulation truth, burst-size posterior draws were multiplied by the fixed capture efficiency, $\alpha = 0.01$, to express them on the simulated count scale. Decay-rate and burst-frequency draws were multiplied by 60 to convert rates from $\mathrm{min}^{-1}$ to $\mathrm{h}^{-1}$. Posterior summaries used a nominal burn-in of 2,500 stored samples and thinning by a factor of two. For shorter chains, the burn-in was reduced to retain at least 100 stored samples before thinning. Consequently, a 5,000-iteration chain contributed 1,250 samples after burn-in and thinning, whereas a 1,500-iteration chain contributed 50. Non-finite draws were excluded from posterior interval and coefficient-of-variation calculations. Point estimates were posterior means, and central 95% intervals were calculated from the 2.5th and 97.5th percentiles of the retained draws. For direct method comparisons, `burstMCMC` results were restricted to successful fits with posterior coefficients of variation no greater than 0.45 for each of the degradation rate, burst size, and burst frequency. The same gene set was used for both methods by intersecting these genes with those passing the `buRst` fit-quality criteria, excluding fits flagged as `bad_hessian` or `strong_tradeoff`. Parameter recovery, interval coverage, and per-gene runtime were compared on this shared gene set. For `buRst`, the stored per-gene elapsed time covered the condition-specific fit or fits and any requested posterior comparison. This included work performed inside the fit, such as profile retrieval, optimization, Hessian evaluation and Laplace draws. Shared batch preparation and parallel-worker startup occurred outside this timer. The validation comparison used a single steady-state fit per gene. Empirical coverage was calculated as the proportion of evaluable genes whose central 95% interval contained the generating parameter value.

### 7 MYC scSLAM-seq preprocessing and pseudobulk analysis

#### 7.1 Read processing, demultiplexing, and cell filtering

Raw MYC scSLAM-seq reads were processed with `cellranger count` using the Cell Ranger Martian runtime v4.0.8 and STAR v2.7.2a against a custom human Ensembl 90 reference containing the Tir1 transgene. Cell Ranger was run with an expected cell number of 6,000 and 24 local cores. The output contained 10,013 estimated cells and 1.68 billion reads, with a mean of 167,908 reads per cell, a median of 26,373 UMIs per cell, and a median of 6,220 genes per cell. Quality metrics were 97.9% valid barcodes, 89.7% Q30 bases in the RNA read, 72.0% of reads assigned to cells, 97.4% mapped to the genome, and 73.5% mapped confidently to the transcriptome.

Sample identities were assigned using MULTI-seq barcode reads and `deMULTIplex`. Cell barcodes from the Cell Ranger filtered feature-barcode matrix were used after removal of the `-1` suffix. Reads from two sequencing lanes were preprocessed with `MULTIseq.preProcess`; cell barcodes and UMIs were extracted from read 1 at positions 1–16 and 17–28, respectively, and sample tags from read 2 at positions 1–8. The lanes were combined and aligned with `MULTIseq.align`. Six sample barcodes were retained: Bar8, Bar9, Bar10, Bar11, Bar13, and Bar14.

Assignment thresholds were determined by sweeping quantiles from 0.01 to 0.99 using `findThresh`. Negative cells were removed over three iterative rounds of `classifyCells`. Negative-cell rescue used `findReclassCells` and `rescueCells`, retaining stable reclassifications. After demultiplexing and rescue, 7,044 cells were assigned to retained sample barcodes; 1,217 were classified as doublets and 1,752 as negative. Assigned cells comprised 1,970 A375 auxin 4sU cells, 1,780 A375 DMSO 4sU cells, 324 A375 DMSO no-4sU cells, 1,341 MaMel63 auxin 4sU cells, 1,141 MaMel63 DMSO 4sU cells, and 488 MaMel63 DMSO no-4sU cells. These counts precede the downstream filtering described below.

The Cell Ranger BAM file was converted to CIT format using `Gedi Bam2CIT` (GEDI v1.0.6f). GRAND3 v3.0.9 generated read-count matrices, NTR estimates, new and old RNA count matrices, and discrete labeling likelihoods using `-D`, `-progress`, `-profile`, `-estim Full`, and `-outputDiscreteLoglik`. Outputs were loaded using `grandR::ReadGRAND3` and genes were filtered using `grandR::FilterGenes`. The object was converted to Seurat with separate total and new RNA assays. Genes were retained when detected with at least one count in at least 50 cells.

Total and new RNA embeddings were calculated independently. Counts were normalized with scale factors of 10,000 for total RNA and 1,000 for new RNA, followed by variable-feature selection, scaling, PCA and UMAP. The first 20 principal components were used for UMAP, with seed 1 for PCA and UMAP.

Initial embeddings were inspected for low-quality tails and isolated populations. Cells in these regions were selected manually in the total RNA UMAP and, for 4sU-labeled cells, additionally in the new RNA UMAP. Selected cells were removed and PCA and UMAP were recalculated.

Cells with inconsistent cell-line annotations were identified using distances to cell-line-specific PCA centroids, defined by component-wise medians. Total RNA PCA was used for all cells; new RNA PCA provided additional evidence for 4sU-labeled cells. A cell was flagged when its nearest centroid belonged to the alternative cell line and the ratio of its distance to the annotated centroid to its distance to the nearest centroid exceeded 1.08 for total RNA or 1.05 for new RNA. Filtering was repeated for up to three rounds, recalculating PCA and UMAP after each round.

The final total RNA UMAP was calculated jointly across all retained cells. New RNA PCA and UMAP were recalculated separately for 4sU-labeled A375 and MaMel63 cells using the same 20-component workflow. Manual-tail filtering removed 657 cells, and cell-line-mismatch filtering removed a further 122 cells, leaving 6,265 cells for downstream analyses. The excluded cell barcodes and retained cell metadata were saved with the analysis outputs. No-4sU controls were retained for labeling quality control but excluded from kinetic fitting.

#### 7.2 Cell-cycle and MYC signature scoring

Cell-cycle scores were calculated using the Seurat 2019 S- and G2/M-phase gene sets when at least ten genes from each set were present in the data. The MYC signature was defined from the gene set reported by [4]. Signature genes were matched to the new RNA assay without regard to letter case. For each cell line, a MYC module score was calculated from the new RNA assay using `Seurat::AddModuleScore` with random seed 1.

For cell-cycle visualizations, separate embeddings were calculated for 4sU-labeled cells in each cell line using the total RNA assay. S- and G2/M-phase genes were used for scaling and PCA, and the first five principal components were supplied to UMAP with 25 neighbors, minimum distance 0.15, cosine distance, and seed 1. Existing saved cell-cycle embeddings were reused when present.

Phase-specific transcriptional responses were assessed using BFDR01 genes with finite new RNA fold changes in all three phases within each cell line. Pairwise phase comparisons used two-sided paired Wilcoxon signed-rank tests across genes, with Benjamini–Hochberg correction across the six comparisons (three phase pairs in each of two cell lines). Effect sizes were median paired differences in $\log_2$ fold change.

#### 7.3 Pseudobulk analysis

Following cell-quality and barcode-mismatch filtering, cells were aggregated into pseudobulk samples according to cell line, treatment, and 4sU-labeling status. The cell-to-pseudobulk mapping was supplied to Grand3, and data were reloaded using `grandR::ReadGRAND3`. Genes were filtered using the standard `grandR::FilterGenes` procedure. Treatment-associated $\log_2$ fold changes were calculated separately for A375 and MaMel63 cells by contrasting auxin-treated and DMSO-treated pseudobulk samples. Fold changes were calculated independently for total and newly synthesized RNA using `grandR::LFC`. For these comparisons, both total and new RNA fold changes were normalized using the total RNA normalization factors. For the cell-cycle-stratified analysis, cells with an assigned G1, S, or G2/M phase were additionally aggregated by cell-cycle phase. Auxin-versus-DMSO $\log_2$ fold changes in total and newly synthesized RNA were then calculated separately for each cell line and cell-cycle phase using the same filtering and normalization procedure. These phase-specific estimates were used to compare the MYC-dependent transcriptional response across G1, S, and G2/M cells. Because the pseudobulk groups did not contain independent biological replicates, fold-change estimates were used descriptively rather than for replicate-based differential-expression testing.

##### 7.4 Derivation of the shared MYC-response signature

A shared MYC-response signature was derived by combining gene-level new RNA responses from A375, MaMel63, K562, and HCT116 cells. A375 and MaMel63 responses were obtained from the pseudobulk analyses described above; K562 and HCT116 responses were taken from the processed DESeq2 result tables accompanying Muhar et al. [4] (GSE100708). All responses were oriented as $\log_2(\mathrm{MYC\ present}/\mathrm{MYC\ depleted})$, so positive values indicate reduced new RNA abundance following MYC depletion. Gene identifiers were harmonized by trimming whitespace, removing terminal numeric version suffixes, and converting symbols to uppercase. Duplicate entries were collapsed within each cell line using the median fold change. Genes with finite responses in all four cell lines were retained, defining a four-cell-line expression-analysis universe of 6,487 genes. No positive minimum baseline-expression threshold or adjusted-$P$-value filter was applied to the Muhar data.

For each retained gene $g$, the response vector was

$$\mathbf{x}_g = (\Delta_{\mathrm{A375},g}, \Delta_{\mathrm{MaMel63},g}, \Delta_{\mathrm{K562},g}, \Delta_{\mathrm{HCT116},g})^{\mathsf{T}} .$$

The unstandardized response vectors were fitted with a two-component, four-dimensional multivariate $t$ mixture using `teigen::teigen`, with component-specific scale matrices and degrees of freedom (model UUUU; `scale = FALSE`). Fitting used emEM initialization with ten initializations of ten iterations each, numerical updating of the degrees of freedom, and random seeds 123 and 456. The iterated M-step and EM convergence tolerances were $10^{-6}$ and $10^{-5}$, with maximum iteration counts of 1,000 and 20,000, respectively. The converged fit with the highest log-likelihood was retained.

The component whose centroid was closest to zero, after dividing each coordinate by its across-gene response standard deviation, was designated the near-zero or background component. The other component was designated the strong-response component; its mean response was required to be positive and greater than the background mean in all four cell lines. This scaling was used only to identify the component, not to fit the response distributions.

Gene-level posterior membership probabilities were evaluated from the final fitted model using `predict`. Let $p_g$ denote the posterior probability that gene $g$ belongs to the strong-response component. The expected number of genes belonging to this component was calculated as $\sum_g p_g$.

To define the BFDR01 signature, genes were ranked by decreasing $p_g$. For the first $k$ ranked genes, the estimated Bayesian false discovery rate was

$$\widehat{\mathrm{BFDR}}(k) = \frac{1}{k}\sum_{j=1}^{k}\left(1 - p_{(j)}\right) ,$$

where $p_{(j)}$ is the posterior probability of the gene at rank $j$. The largest ranked set satisfying $\widehat{\mathrm{BFDR}}(k) \leq 0.01$ was selected. This criterion limits the average posterior background probability within the selected set under the fitted model; it does not require every selected gene to have $p_g \geq 0.99$. No additional requirement for positive responses in every cell line was imposed.

For main Figure 6d, component curves represent mixture-weighted marginal $t$ densities from the same joint four-dimensional model, rather than separate cell-line-specific fits. Signature membership was determined from the joint posterior probabilities. For main Figure 6e, cross-cell-line background correlations were derived from the separately estimated covariance matrix of the near-zero component. For a component with scale matrix $\mathbf{S}$ and degrees of freedom $\nu > 2$, the covariance is

$$\mathrm{Cov}(\mathbf{x}) = \frac{\nu}{\nu - 2}\mathbf{S}.$$

The scalar correction does not affect the corresponding correlation matrix. These values describe correlations within the fitted component, rather than empirical correlations among genes assigned to it by a posterior-probability threshold.

Separately, pairwise Pearson correlations of new RNA responses were calculated using all eligible genes and after excluding the Muhar top-100 signature genes (main Figure 6c). Each comparison used the genes available for that cell-line pair; 95% confidence intervals were obtained using Fisher's $z$ transformation.

##### 7.5 RNA half-life analysis

Steady-state RNA half-lives were estimated from the 2 h labeling experiment using `grandR::ComputeSteadyStateHalfLives`. For the A375 DMSO condition, genes with finite half-life, total RNA fold-change, and new RNA fold-change estimates were grouped into

$$t_{1/2} < 4\,\mathrm{h}, \qquad 4 \leq t_{1/2} < 10\,\mathrm{h}, \qquad t_{1/2} \geq 10\,\mathrm{h}.$$

Within each group, agreement between total and new RNA responses to MYC depletion was summarized using Pearson's correlation coefficient.

#### 8 Genome-wide inference after MYC depletion

Bursting kinetics were inferred separately for 4sU-labeled A375 and MaMel63 cells; no-4sU controls were excluded from kinetic fitting. Before model fitting, genes were retained when they had at least one observed count in at least 50 cells. Only genes with unique gene-symbol annotations were included. For each cell line, DMSO-treated and auxin-treated cells were analyzed as a paired condition comparison. The DMSO condition was fitted under the steady-state model. The auxin condition was fitted under the non-steady-state model, using the corresponding DMSO population as reference. Both the 4sU-labeling duration and the modeled propagation interval were set to 2 h. As described in Section 4.5, the DMSO total RNA distribution was used to approximate the RNA population at labeling onset, without explicitly modeling changes during the preceding 2 h auxin pretreatment. For each condition-specific fit, 20,000 parameter draws were generated from the local Gaussian approximation. Gene-wise condition comparisons were then performed from the corresponding draws. The random seed was set to 123. Genes were processed in parallel using the configured worker count, recorded in the archived run provenance, with a configured timeout of 300 s per condition-specific fit. The paired comparison was performed after the two condition fits. Downstream burst-change analyses retained genes with successful fits in both conditions. Fits classified as `bad_hessian` or `strong_tradeoff` were excluded. Kinetic changes were classified using the procedure described in Section 4.9. For visualization, detailed classifications were grouped into five display categories: frequency, size, both, mechanism unresolved, and no supported change. Concordant joint responses and all opposing-response subclasses were pooled into the "both" display category. Scatterplots retained genes with successful comparisons and finite posterior-median fold changes, including genes with unresolved mechanisms or no supported change. Genes without a successful comparison were excluded. For visualization, fold changes were expressed in the DMSO-relative-to-auxin direction:

$$\Delta_q = \log_2\left(\frac{q_{\mathrm{DMSO}}}{q_{\mathrm{Aux}}}\right), \qquad q \in \{B, k_m\}.$$

Positive values therefore indicate a reduction following MYC depletion. For signature-restricted visualizations, genes were selected by membership in the shared four-cell-line MYC-response signature defined at a Bayesian false discovery rate of 1% (BFDR01). Signature membership was matched after trimming whitespace, removing terminal numeric version suffixes, and converting gene symbols to uppercase. After condition-fit filtering and exclusion of unsuccessful comparisons or non-finite posterior-median fold changes, 3,485 genes were available for visualization in A375 and 5,772 in MaMel63, including 158 and 251 BFDR01 signature genes, respectively.

### 9 MYC ChIP-seq processing and promoter annotation

#### 9.1 MYC ChIP-seq processing

Data was preprocessed as described in Methods. The downstream analyses used the resulting promoter-associated peak tables containing MYC and input coverage for each peak–gene assignment. Input-control coverage was rescaled using the factor

$$\frac{940008}{1512647}$$

before comparison with the MYC ChIP-seq signal. For each promoter-associated peak $p$, a background-subtracted signal was calculated as

$$S_p = C_{\mathrm{MYC},p} - C^*_{\mathrm{input},p}, \tag{81}$$

where $C^*_{\mathrm{input},p}$ denotes rescaled input coverage. A MYC-to-input coverage ratio was calculated as

$$R_p = \frac{C_{\mathrm{MYC},p}}{C^*_{\mathrm{input},p}}. \tag{82}$$

Peaks with zero input coverage were assigned a missing ratio rather than an infinite value.

#### 9.2 Motif and promoter annotations

Peak coordinates were restricted to chromosomes 1–22, X, and Y and converted to the hg38 chromosome naming convention. The 500-bp extension introduced during the initial peak-to-promoter intersection was removed before calculating each peak center. A 500-bp sequence window extending from 250 bp upstream to 249 bp downstream of the peak center was extracted from the hg38 reference genome using `BSgenome.Hsapiens.UCSC.hg38`. MYC-associated E-box motifs were identified by exact sequence matching within the peak-centered windows. The primary E-box annotation counted occurrences of

$$\mathrm{CACGTG},\ \mathrm{CACGCG},\ \mathrm{CATGCG},\ \mathrm{CATGTG},\ \mathrm{CACGAG},\ \mathrm{CACATG},$$

together with their reverse-complement sequences. A broader count comprising all $\mathrm{CANNTG}$ sequences and selected non-canonical E-box variants was also calculated for exploratory annotation but was not used for the primary E-box classification. CpG-island coordinates for hg38 were obtained from the UCSC `cpgIslandExt` table. Peak-centered windows were classified as overlapping a CpG island when they shared at least one genomic position with a UCSC CpG-island interval. TATA-like motifs were searched in a strand-aware interval extending from 100 bp upstream to 50 bp downstream of the annotated transcription start site. The exact motifs

$$\mathrm{TATAAAA},\ \mathrm{TATATAA},\ \mathrm{TATAAAT},\ \mathrm{TATATAT}$$

were considered. Motif positions were transformed into transcription-oriented coordinates, such that negative positions denote locations upstream of the transcription start site. A canonical promoter TATA motif was defined as a motif located between 50 and 15 bp upstream of the transcription start site. When several canonical motifs occurred in one promoter, the motif nearest to $-30$ bp was selected.

##### 9.3 Representative peak and gene-level annotation

For promoters associated with multiple MYC peaks, the peak with the largest MYC-to-input coverage ratio was selected as the representative peak for that promoter. Because the bursting and new RNA results were estimated at gene level, one representative promoter was subsequently selected for each gene. This selection was based primarily on the largest background-subtracted MYC signal, with the MYC-to-input ratio and original peak score used to break ties. The total number of distinct promoter-associated MYC peaks was retained separately for each gene before representative-peak selection.

#### 10 Integration of MYC binding with transcriptional and kinetic responses

##### 10.1 Dataset integration

Promoter-level MYC annotations were integrated with A375 pseudobulk new RNA fold changes and gene-specific changes in burst size and burst frequency. Ensembl version suffixes were removed before joining datasets. Bursting and pseudobulk tables were joined by gene identifier, while promoter annotations were added by left join so that genes without assigned MYC peaks remained available for categorical peak-presence analyses. For quantitative analyses, genes were required to have a finite positive MYC-to-input coverage ratio and finite estimates of new RNA, burst-size, and burst-frequency responses. Quantitative MYC binding was represented as

$$X_g = \log_{10}(R_g), \tag{83}$$

where $R_g$ is the coverage ratio of the representative promoter-associated peak for gene $g$. Response signs were oriented so that positive values indicate a decrease following MYC depletion:

$$\Delta_q = \log_2\left(\frac{q_{\text{DMSO}}}{q_{\text{Aux}}}\right), \qquad q \in \{\text{new RNA}, B, k_m\}. \tag{84}$$

For the quantitative binding scatterplots, separate univariable ordinary least-squares regressions related $X_g$ to new RNA, burst-size, and burst-frequency responses. These regressions used the common complete-case set defined above. Fitted slopes, coefficients of determination ($R^2$), and two-sided $P$ values for the slope were reported.

Associations between pseudobulk new RNA responses and kinetic responses were also assessed by ordinary least-squares regression and Pearson's correlation using this common gene set. Kinetic fold changes for these regressions were calculated from condition-specific maximum-likelihood estimates; the classification scatterplots described in Section 8 used posterior-median fold changes. Regression lines were displayed with 95% confidence intervals, and BFDR01 genes were highlighted in the new-RNA-versus-kinetics scatterplots.

##### 10.2 Categorical and predictive analyses

The gene-level new RNA response was defined as

$$D_g = \log_2\left(\frac{\text{new RNA}_{\text{DMSO},g}}{\text{new RNA}_{\text{Aux},g}}\right). \tag{85}$$

Larger positive values represent a stronger reduction in new RNA synthesis following MYC depletion. Among genes with an assigned peak and finite quantitative binding value, genes were divided into low, middle, and high MYC-binding tertiles according to $\log_{10}(R_g)$. The monotonic association between quantitative binding and the new RNA response was summarized using Spearman's rank correlation. Genes were also grouped according to the total number of distinct promoter-associated MYC peaks:

$$0, \qquad 1, \qquad \geq 2.$$

To assess whether E-box presence was associated with transcriptional response at comparable MYC occupancy, genes with an assigned MYC peak and finite quantitative binding values were divided into low, middle, and high binding tertiles. Within each tertile, new RNA responses were compared between promoters with and without an E-box using two-sided Wilcoxon rank-sum tests. Benjamini–Hochberg correction was applied across the three within-tertile comparisons. For the peak-count analysis, correction was applied across the three pairwise comparisons between the zero-, one-, and multiple-peak groups. Jittered points were subsampled to at most 300 genes per group for display, while all available genes were retained for boxplots and statistical tests. The ability of promoter-binding features to identify strongly MYC-dependent genes was evaluated by receiver-operating-characteristic analysis. A gene was classified as strongly MYC dependent when

$$D_g \geq 1.$$

Four predictors were considered: presence of at least one MYC peak, total MYC peak count, quantitative MYC-to-input coverage ratio, and a combined model containing quantitative coverage and E-box presence. Genes without an assigned peak were assigned zero quantitative occupancy. The combined predictor was fitted by logistic regression:

$$\operatorname{logit}\Pr(Z_g = 1) = \gamma_0 + \gamma_1 \operatorname{scale}[\log(1 + R_g)] + \gamma_2 E_g, \tag{86}$$

where $Z_g$ indicates strong MYC dependence and $E_g$ indicates the presence of at least one E-box in the assigned peak window. Predictions were generated using stratified five-fold cross-validation. ROC curves and AUCs were calculated using `pROC`, together with 95% confidence intervals for the AUC.

##### 10.3 Associations of promoter architecture with transcriptional and kinetic responses

To assess associations between promoter features and MYC-dependent responses, separate multivariable linear models were fitted for new RNA, burst-size, and burst-frequency $\log_2(\mathrm{DMSO}/\mathrm{Aux})$ fold changes. Positive responses therefore indicate higher RNA synthesis or larger kinetic parameter values in the presence of MYC. For each response $Y_g$, the model was

$$Y_g = \beta_0 + \beta_1 X_{\mathrm{MYC},g} + \beta_2 X_{\mathrm{E-box},g} + \beta_3 X_{\mathrm{TATA},g} + \beta_4 X_{\mathrm{CpG},g} + \beta_5 X_{\mathrm{distance},g} + \beta_6 X_{\mathrm{position},g} + \varepsilon_g. \tag{87}$$

Quantitative MYC binding was represented by the $\log_{10}$ MYC-to-input coverage ratio. E-box annotations were encoded as presence or absence of at least one E-box. TATA-box presence and CpG-island overlap were also encoded as binary variables. Absolute peak-to-TSS distance was expressed in base pairs and transformed as $\log_{10}(1 + |\mathrm{distance}|)$. Continuous predictors were not standardized. Peak position distinguished downstream from upstream peaks, with upstream as the reference category; peaks located exactly at the TSS were excluded. All three response models used a common gene set with finite new RNA, burst-size, and burst-frequency responses and complete predictor information. The input bursting estimates had passed the condition-fit exclusions for `bad_hessian` and `strong_tradeoff`; no additional precision filtering was applied. Predictor-specific $P$ values were adjusted jointly using the Benjamini–Hochberg procedure across the 18 non-intercept coefficients from the three models.

##### 10.4 Signature comparisons

Burst-size and burst-frequency responses were compared between other genes and genes belonging to the Muhar or BFDR01 signatures. Analyses were restricted to genes in the four-cell-line expression-analysis universe with finite estimates for the kinetic response being compared. Other genes excluded the union of both signatures; genes shared between the signatures appeared in both signature groups. Each signature was compared with other genes using a two-sided Wilcoxon rank-sum test, with Benjamini–Hochberg correction across the four comparisons.

#### Software and reproducibility

Workflow configurations specify random seeds and worker counts. Workflow configurations and R session information, including loaded package versions, were recorded alongside analysis results. Parallel worker counts were held fixed within each simulation run because changing the number of workers can change simulated random streams even when the same seed is supplied.

### References

[1] Christopher Jürges, Lars Dölken, and Florian Erhard. "Dissecting Newly Transcribed and Old RNA Using GRAND-SLAM". In: *Bioinformatics* 34.13 (July 1, 2018), pp. i218–i226. ISSN: 1367-4803, 1367-4811. DOI: 10.1093/bioinformatics/bty256. URL: https://academic.oup.com/bioinformatics/article/34/13/i218/5045735 (visited on 09/09/2025).

[2] Abhyudai Singh and Pavol Bokes. "Consequences of mRNA Transport on Stochastic Variability in Protein Levels". In: *Biophysical Journal* 103.5 (Sept. 5, 2012), pp. 1087–1096. ISSN: 1542-0086. DOI: 10.1016/j.bpj.2012.07.015. PMID: 23009859.

[3] David M. Edwards, Philip Davies, and Daniel Hebenstreit. "Synergising Single-Cell Resolution and 4sU Labelling Boosts Inference of Transcriptional Bursting". In: *Genome Biology* 24.1 (June 16, 2023), p. 138. ISSN: 1474-760X. DOI: 10.1186/s13059-023-02977-y. PMID: 37328900.

[4] Matthias Muhar et al. "SLAM-Seq Defines Direct Gene-Regulatory Functions of the BRD4-MYC Axis". In: *Science* 360.6390 (May 18, 2018), pp. 800–805. ISSN: 0036-8075, 1095-9203. DOI: 10.1126/science.aao2793. URL: https://www.science.org/doi/10.1126/science.aao2793 (visited on 09/09/2025).
